# Genetic activation of canonical RNA interference in mice

**DOI:** 10.1101/2023.12.29.573630

**Authors:** Valeria Buccheri, Josef Pasulka, Radek Malik, Zuzana Loubalova, Eliska Taborska, Filip Horvat, Marcos Iuri Roos Kulmann, Irena Jenickova, Jan Prochazka, Radislav Sedlacek, Petr Svoboda

## Abstract

Canonical RNA interference (RNAi) is sequence-specific mRNA degradation guided by small interfering RNAs (siRNAs) made from double-stranded RNA (dsRNA) by RNase III Dicer. RNAi has different roles including gene regulation, antiviral immunity or defense against transposable elements. In mammals, RNAi is constrained by Dicer, which is adapted to produce microRNAs, another class of small RNAs. However, RNAi exists in mouse oocytes, which employs a truncated Dicer variant. A homozygous mutation to express only the truncated variant (ΔHEL1) causes dysregulation of microRNAs and perinatal lethality in mice. Here, we report the phenotype and RNAi activity in *Dicer^ΔHEL1/wt^*mice, which are viable, show minimal miRNome changes but their endogenous siRNA levels are increased by an order of magnitude. We show that siRNA abundance is limited by available dsRNA but not by PKR, a dsRNA sensor of innate immunity. Expressing dsRNA from a transgene, functional RNAi *in vivo* was induced in heart. *Dicer^ΔHEL1/wt^* mice thus represent a new model for researching mammalian canonical RNAi *in vivo* and offer an unprecedented platform for addressing claims about its biological roles.

## Introduction

RNase III Dicer generates small RNAs, which guide ribonucleoprotein complexes in microRNA (miRNA) and RNA interference (RNAi) pathways (reviewed in (1)). In the gene-regulating miRNA pathway, Dicer cleaves genome-encoded small hairpin miRNA precursors (pre-miRNA) into ∼21-23 nt duplexes; one of the strands (miRNA) is loaded onto an AGO protein and guides gene repression while the other one is discarded. In RNAi, which functions in gene regulations, retrotransposon repression, and antiviral immunity, long double-stranded RNA (dsRNA) is cut by Dicer into 22 nt small interfering RNA (siRNA) duplexes, then one siRNA strand is bound by an endonucleolytic AGO protein and guides sequence-specific cleavage of RNAs.

Co-existence of small RNA biogenesis and RNA-induced silencing complex (RISC) formation in RNAi and miRNA pathways evolved in different ways. In *D. melanogaster*, the pathways diverged such that each employs dedicated Dicer and AGO proteins (2). *C. elegans* utilizes a single Dicer for miRNA and RNAi pathways and small RNAs are sorted onto different AGO proteins (3). Invertebrate RNAi provides defense against parasitic nucleic acids – this role appears largely lost in vertebrates where genomic and phylogenetic data support the primary role of Dicer in the miRNA pathway (reviewed in (4)). Vertebrate Dicers are highly conserved when compared to Dicer divergence across animals, particularly those utilizing antiviral RNAi (5), and lack ATP-dependent Dicer processivity facilitating siRNA production (6). Similarly, vertebrate AGO proteins associate primarily with miRNAs and have lower rates of molecular evolution than siRNA-associated AGO proteins from other taxa (7).

Canonical RNAi (i.e. long dsRNA-induced sequence-specific mRNA degradation (8)) was observed in mammalian somatic cells in several early studies (9–14) suggesting that RNAi might exist under favorable conditions. The endogenous canonical RNAi in mammals, however, appears severely constrained by (I) scarcity of cellular long dsRNA and its inefficient processing by Dicer (15) and (II) innate immunity mechanisms, which respond to dsRNA and restrict RNAi (16,17). The inhibition may be reciprocal (18,19) but there is no simple antagonistic relationship among RNAi and other dsRNA-responding pathways (20,21). An exceptional instance of mammalian RNAi evolved in mouse oocyte where an N-terminally truncated oocyte-specific Dicer variant (Dicer^O^) supports highly active and essential endogenous RNAi (22). RNAi is associated with the loss of the N-terminal DExD/H helicase domain (HEL1 helicase subdomain), which otherwise provides functional adaptation of the mammalian Dicer to miRNA biogenesis and inhibits processing of long dsRNA into siRNAs (23). Removal of HEL1 enhances long dsRNA processing *in vivo* and supports both, miRNA biogenesis and endogenous RNAi (15,22).

We have developed a mouse mutant where the endogenous *Dicer* gene was genetically engineered to express the more active Dicer variant lacking the HEL1 domain (Dicer^ΔHEL1^) (23). As a control, we generated an allele denoted *Dicer^SOM^*, which expresses an HA-tagged full-length Dicer variant (24) but lacks the same intronic sequences like the *Dicer^ΔHEL1^*allele (Fig.1A). Both engineered *Dicer* alleles express Dicer proteins of expected sizes and at comparable levels (23). *Dicer^SOM/SOM^*mice are viable but females are sterile because this modification eliminates expression of Dicer^O^, which is essential for female fertility (22,24). *Dicer^ΔHEL1/ΔHEL1^* mice exhibit severe miRNome dysregulation, growth retardation, defects in the cardiopulmonary system, and perinatal lethality (23). Consequently, the lethal phenotype of *Dicer^ΔHEL1/ΔHEL1^* animals precludes obtaining the maximum Dicer activity *in vivo* originating from natural transcriptional control of the gene. At the same time, *Dicer^ΔHEL1/wt^*heterozygotes are viable and fertile, demonstrating that one allele expressing *Dicer^ΔHEL1^* is well tolerated *in vivo*.

**Figure 1.**
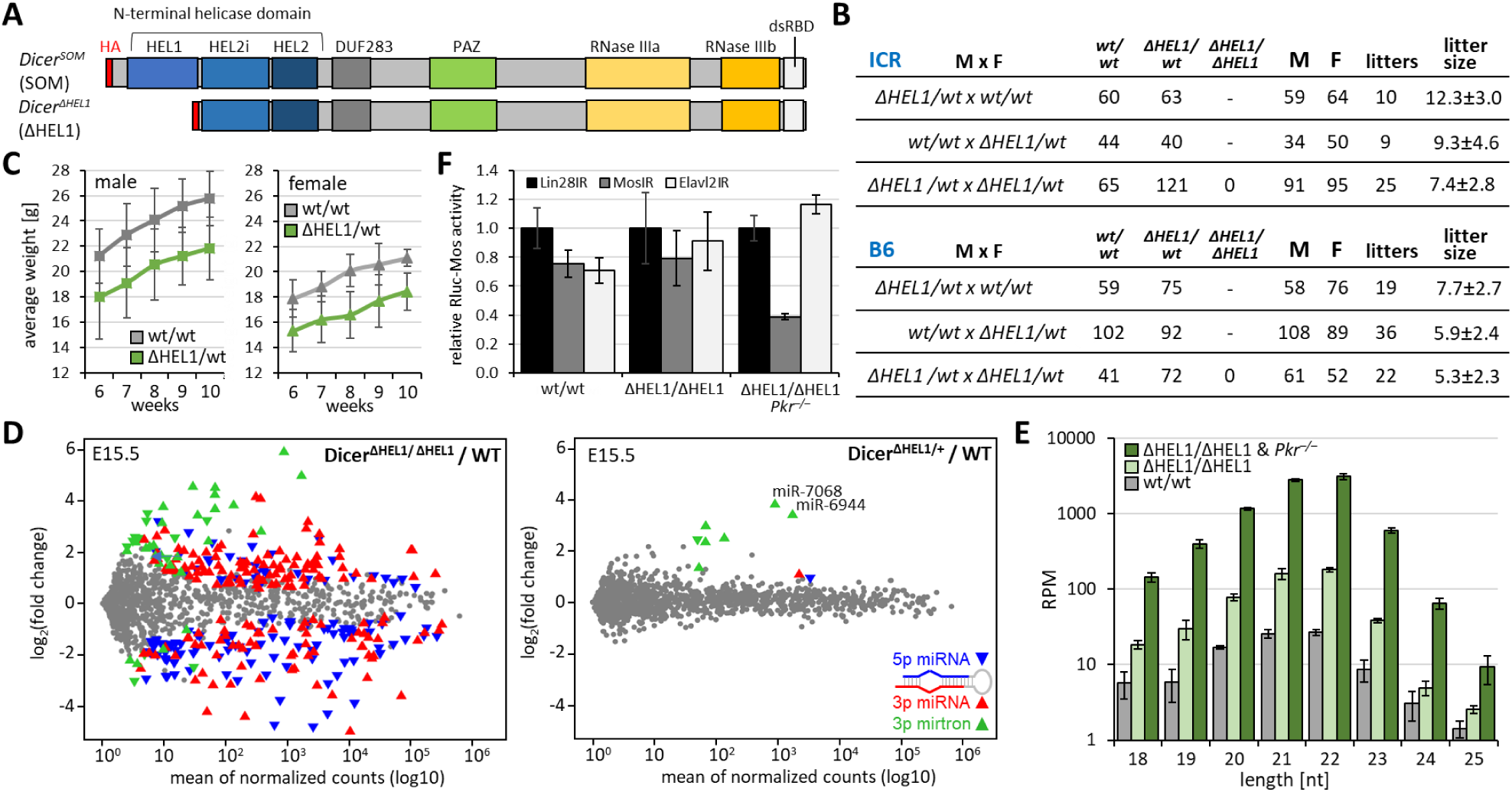
Extended analysis of *Dicer^ΔHEL1^*mutants. (A) Schematic depiction of Dicer^SOM^ and Dicer^ΔHEL1^protein variants. (B) Breeding performance of heterozygous mutants. (C) Weight of 6-10 weeks old *Dicer^ΔHEL1/wt^* and wild type littermates on the C57Bl/6NCrl background. (D) MA plots of small RNA-seq analysis of whole *Dicer^ΔHEL1/ΔHEL1^*and *Dicer^ΔHEL1/wt^* E15.5 embryos compared to wild type (WT) E15.5 embryos. Depicted are changes in levels of annotated murine miRNAs (miRBase 22.1 (61)). Significantly dysregulated 5p and 3p miRNAs (DESeq p-value 0.05) are shown as oriented blue ▾and red ▴ triangles, respectively. Significantly dysregulated mirtrons are represented by green triangles whose orientation indicates 5p and 3p miRNA strands. The published MA plot for *Dicer^ΔHEL1/ΔHEL1^* (23) is included here for convenient comparison of changes in homo- and heterozygotes. (E) In ESCs, the loss of PKR enhances production of endo-siRNAs from a dsRNA-expressing plasmid MosIR (22). The MosIR plasmid was transfected into ESCs with indicated genotypes in triplicates and small RNAs were analyzed by RNA-seq. The y-axis depicts reads per million (RPMs) calculated for all 18-32 nt mapped reads. Error bars = SD. (F) Endogenous RNAi activity in genetically modified ESCs. RNAi was analyzed using luciferase-based RNAi assay (15). Cells were co-transfected with plasmids expressing firefly luciferase (non-targeted reporter), *Renilla* luciferase reporter with *Mos* sequence in the 3′ UTR (RNAi-targeted reporter), and one of the plasmids expressing a long hairpin RNA (Lin28IR, MosIR or Elavl2IR). Results are presented as a ratio of a *Renilla* luciferase activity divided by a non-targeted firefly luciferase activity scaled to Lin28IR, which was set to 1. The transfection experiments were performed in a triplicate, error bars = SD.

Here, we investigated the phenotype of *Dicer^ΔHEL1/wt^*mice and assessed their ability to mount functional RNAi response induced by long dsRNA. We report that a single *Dicer^ΔHEL1^* allele affects canonical miRNA biogenesis much less than expected while it is sufficient to increase biogenesis of specific mirtrons, a non-canonical miRNA class (25,26). Furthermore, siRNA biogenesis is enhanced but robust endogenous RNAi is rarely observed, even in the Protein kinase R (*Pkr*, official gene symbol *Eif2ak2*) null background. Consequently, RNAi effect was observed in heart where high siRNA abundance was experimentally achieved with high level of expression of long dsRNA. Altogether, we provide an important framework for understanding limits of mammalian canonical RNAi *in vivo* and a unique set of genetically modified mouse models for further research on mammalian RNAi and other dsRNA-response pathways.

## Results and Discussion

### Dicer^ΔHEL1/wt^ mice have a subtle but discernible phenotype

*Dicer^ΔHEL1/ΔHEL1^* mice were growth retarded, had developmental defects, and died perinatally (23). In contrast, *Dicer^ΔHEL1/wt^*animals appeared normal. Fertility of *Dicer^ΔHEL1/wt^* was normal in ICR and C57Bl/6NCrl genetic backgrounds and there was no significant difference in the birth rate and survival of wild types and heterozygotes (Fig. 1B). For detailed phenotype assessment, a standard phenotype screening of *Dicer^ΔHEL1/wt^* animals was performed at the Czech Centre for Phenogenomics (the full report is available in the Supplementary File 1). The phenotyping pipeline included assessment of embryonic development, anatomy, histopathology, metabolism, hematology, immunology, cardiopulmonary function, and neurobiology (including vision, hearing, and behavior). We have found that *Dicer^ΔHEL1/wt^* animals were smaller than wild type littermates (Fig. 1C and S1A). Similar shapes of growth curves suggests that growth retardation occurred earlier, possibly between E12.5 and E15.5 when major growth retardation of *Dicer^ΔHEL1/ΔHEL1^*occurred and a slight deviation was also observed in *Dicer^ΔHEL1/wt^*animals (23). *Dicer^ΔHEL1/wt^* had overall normal anatomy (Supplementary File 1) including that of the cardiopulmonary system, which was severely affected in *Dicer^ΔHEL1/ΔHEL1^*mice (23). Heart and lung functions were normal (Supplementary File 1 and Fig. S1B). Hematopoietic parameters were also normal, including those, which were significantly different in *Dicer^ΔHEL1/ΔHEL1^*mutants (Fig. S1C). Taken together, *Dicer^ΔHEL1/wt^* animals were viable, fertile and exhibited a minor growth reduction as the major phenotypic feature.

### Mirtrons dominate miRNome changes in *Dicer^ΔHEL1/wt^* mutants

The *Dicer^ΔHEL1/ΔHEL1^* genotype was associated with massive miRNA dysregulation, which was consistent with altered function of Dicer lacking the N-terminal HEL1 domain (23). Because *Dicer^ΔHEL1/ΔHEL1^* animals died perinatally, we analyzed small RNAs in E15.5 embryos where we could compare miRNome changes in *Dicer^ΔHEL1^* homozygotes and heterozygotes. Consistent with the minor phenotype, *Dicer^ΔHEL1/wt^*E15.5 embryos showed minimal miRNome changes when compared to wild type littermates (Fig. 1D, right panel). Majority of significantly dysregulated miRNAs (7/9 miRNAs) were mirtrons, miRNAs with non-canonical biogenesis arising from small spliced introns (27). Increased levels of a mirtron subset were consistent with previously reported increased cleavage of their precursors by the Dicer^ΔHEL1^ variant (23). Notably, mirtrons miR-7068-3p (19-fold) and mir-6944-3p (18-fold) achieved considerable abundance, which could be physiologically relevant (Fig. 1D).

Further analysis of miRNome changes in adult organs was done in animals that were also used for siRNA analysis. Thus, in addition to the *Dicer^ΔHEL1/wt^* or control *Dicer^SOM/wt^* genotypes, these mice carried a dsRNA-expressing transgene Tg(CAG-EGFP-MosIR), for simplicity referred to as MosIR transgene, and one or two mutant alleles of *Pkr*. The MosIR transgene is ubiquitously expressing a long dsRNA hairpin with *Mos* gene sequence. This expressed long dsRNA is well tolerated and able to induce RNAi in oocytes but not in soma (28). Analysis of MosIR-derived small RNAs in *Dicer^ΔHEL1/ΔHEL1^*ESCs showed that replacement of the full-length Dicer with *Dicer^ΔHEL1^*variant increases MosIR-derived 21-23 nt small RNAs by an order of magnitude. Notably, the loss of PKR in *Dicer^ΔHEL1/ΔHEL1^* ESCs added another order of magnitude in MosIR-derived 21-23 nt small RNA abundance (Fig. 1E). With this small RNA abundance, we observed sequence-specific knock-down of a luciferase reporter carrying a complementary sequence (Fig. 1F). It is unclear what accounts for the additional increase in siRNA abundance in PKR mutants in ESCs. It was reported that PKR inhibits expression of transiently transfected plasmids in mammalian cultured cells (29,30). We have observed that expression of transiently transfected plasmids but not of genome-integrated genes was negatively affected in a PKR-dependent manner when one of the transiently transfected plasmids generated dsRNA (30). PKR knock-down resulted in two-fold increase of reporter mRNA levels 24 hours post-transfection (30). We thus analyzed the abundance of 18-32 nt RNA fragments from MosIR in *Dicer^ΔHEL1/ΔHEL1^ Pkr^-/-^* ESCs and observed a moderate increase (40% higher) of EGFP-derived fragments (Fig. S1D). At the same time, MosIR sequences of other lengths than 21-23 nt nucleotides (e.g. 18, 19, 24, and 25 nt in Fig. 1E) showed almost an order of magnitude higher abundance in *Dicer^ΔHEL1/ΔHEL1^ Pkr^-/-^* ESCs. Since the sequencing analysis and RNAi assessment were performed 48 h post-transfection, we hypothesize, that PKR may negatively affect the initial plasmid expression while the effect weakens at later timepoints. At the same time, MosIR dsRNA fragmentation may leave behind duplexes of various sizes, which are likely more stable than degradation of single-stranded EGFP RNA. Consequently, we would observe increased levels of MosIR fragments in the whole range of 18-25 nucleotide fragments in *Dicer^ΔHEL1/ΔHEL1^ Pkr^-/-^* ESCs. Because PKR binds MosIR dsRNA (30) and Dicer interacts with PKR (21), we cannot exclude that the positive effect of PKR loss on RNAi includes improved dsRNA substrate recognition or that PKR binding reduces Dicer^ΔHEL1^ activity. Since *Pkr* knockout (*Pkr*^−/−^) mice are viable and phenotypically indistinguishable from wild type animals (31), we thus took the opportunity and included *Pkr* knock-out background into our experimental analysis.

Altogether, RNA-seq experiments provided small RNA profiles from brain, heart, liver, spleen and thymus of *Dicer^ΔHEL1/wt^ Pkr^+/–^*Tg(CAG-EGFP-MosIR) animals, which could be compared with small RNA profiles of normal age-matched C57Bl/6NCrl (Fig. 2A) and other specific genotypes to investigate miRNome changes and activation of siRNA biogenesis. In adult organs of *Dicer^ΔHEL1/wt^* mice, we observed a similar picture as in E15.5 *Dicer^ΔHEL1/wt^* embryos: canonical miRNAs showed small changes while mirtrons were among the most upregulated miRNAs (Fig. 2A). When comparing an age-matched laboratory stock of C57Bl/6NCrl mice with *Dicer^ΔHEL1/wt^ Pkr^+/–^* Tg(CAG-EGFP-MosIR) animals, many more canonical miRNAs appeared slightly but significantly differentially expressed than in E15.5 embryos. However, most of these differences may not originate specifically from the loss of the helicase domain because comparison of *Dicer^ΔHEL1/wt^ Pkr^+/–^* Tg(CAG-EGFP-MosIR) organs with those from *Dicer^SOM/wt^ Pkr^+/–^* Tg(CAG-EGFP-MosIR) showed minimal differential miRNA expression as well as miRNA expression variability, especially in heart and spleen where mirtrons were over-represented again (Fig. S2A).

**Figure 2.**
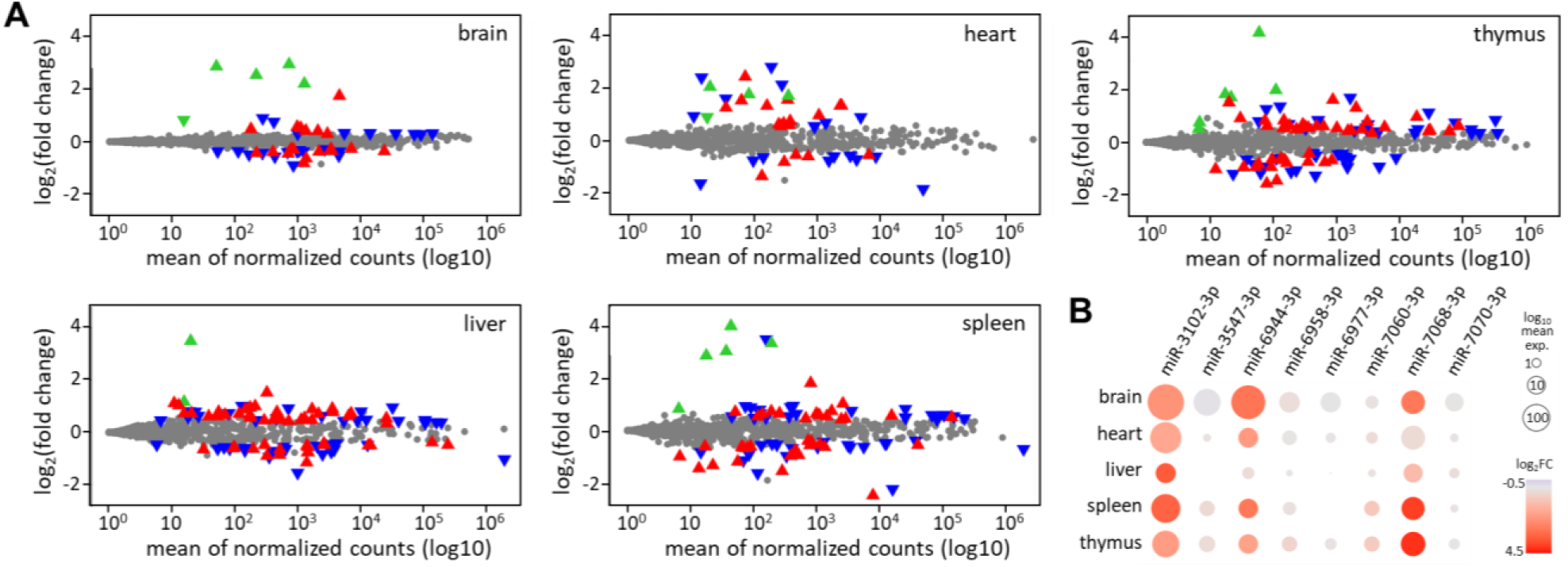
miRNA dysregulation in organs of *Dicer^ΔHEL1/wt^* mice. (A) MA plots depicting changes of annotated murine miRNAs (miRBase 22.1 (61)) in *Dicer^ΔHEL1/wt^ Pkr^+/–^* Tg(CAG-EGFP-MosIR) animals relatively to age-matched normal C57Bl/6NCrl controls. Significantly dysregulated 5p and 3p miRNAs (DESeq p-value 0.05) are shown as oriented blue ▾and red ▴ triangles, respectively. Mirtrons are represented by green triangles whose orientation indicates 5p and 3p miRNA strands. (B) Abundance and expression changes of selected mirtrons in different organs of *Dicer^ΔHEL1/wt^ Pkr^+/–^* Tg(CAG-EGFP-MosIR) relative to C57Bl/6NCrl mice.

Taken together, small RNA analysis from *Dicer^ΔHEL1/wt^*E15.5 embryos as well as from adult organs implies that presence of a single full-length Dicer allele reduces aberrant effects of the *Dicer^ΔHEL1^*allele on canonical miRNA biogenesis more than expected. We hypothesize that the pre-dicing state formed by the full-length Dicer (23,32) facilitates recognition of canonical miRNA precursors *in vivo* while the ΔHEL1 variant interacts with a broader range of substrates. Consequently, the full-length Dicer would contribute to miRNA biogenesis significantly more than the promiscuous truncated Dicer expressed from the *Dicer^ΔHEL1^* allele. At the same time, suboptimal substrates, such as mirtrons and dsRNA, would be still more efficiently processed by the truncated Dicer and would remain upregulated. Accordingly, a set of mirtrons identified in our earlier E15.5 analysis (Fig. 1D and (23)), was consistently upregulated in *Dicer^ΔHEL1/wt^*tissues (Fig. 2B and S2B). Upregulated mirtrons were a subset of a much larger mirtron population (25,27). With their extended stems (the extended miR-3102 stem is cleaved twice by Dicer (Fig. S2B)), they are suboptimal full-length Dicer substrates and are low expressed in wild type tissues. Their biological roles, if any, remain unclear. It is possible that their activity becomes significant when they become abundant in *Dicer^ΔHEL1^*mutants (Fig. 2B) and could contribute to reduced growth. Thus, *Dicer^ΔHEL1/wt^* animals provide an interesting model to further investigate mirtron significance *in vivo*, especially in combination with mutations disrupting above-mentioned mirtrons.

### siRNA biogenesis is stimulated *in vivo* by *Dicer^ΔHEL1^* but not by the loss of PKR

To evaluate siRNA production *in vivo*, we first analyzed 21-23 nt RNAs from *Optn* and *Anks3* loci, which give rise to 21-23 nt RNAs in Dicer-dependent manner, i.e. bona fide endo-siRNAs (22,33,34). The *Optn* locus carries an inverted repeat at the 3’ end of the gene, which is predicted to form a stem loop with ∼300 bp stem and ∼ 1 kb loop and gives rise to low-abundant 21-23 nt small RNAs from the base-pairing region (Fig. 3A). RNA sequencing data from wild type mouse organs show that low levels of mRNA and endo-siRNAs from the *Optn* exist in many organs (Fig. 3B,C and S3A). Most endo-siRNAs are found in heart, notably much more than in intestine, where *Optn* mRNA is expressed at the same level. Analysis of heart and liver from *Dicer^ΔHEL1/wt^* mice showed that a single allele of *Dicer^ΔHEL1^* increased abundance of endo-siRNAs by an order of magnitude but loss of PKR did not have a statistically significant effect (Fig. 3D, p-value >0.05, one-tailed t-test). Robust endo-siRNA upregulation was also observed in brain and thymus (Fig. S3B). *Anks3*, which harbors an inverted repeat in an intron, showed minimal expression in most organs but upregulation of *Anks3* endo-siRNAs was observed in heart, spleen, and thymus of *Dicer^ΔHEL1/wt^ Pkr^+/–^* Tg(CAG-EGFP-MosIR) animals (Fig. S3B). Altogether, analysis of small RNA sequencing of embryos and organs showed that endo-siRNAs from endogenous dsRNA are rare, which is consistent with analyses of Dicer^O^ and Dicer^ΔHEL1^ variants (15,22), which suggested that long dsRNA abundance may be the limiting factor.

**Figure 3.**
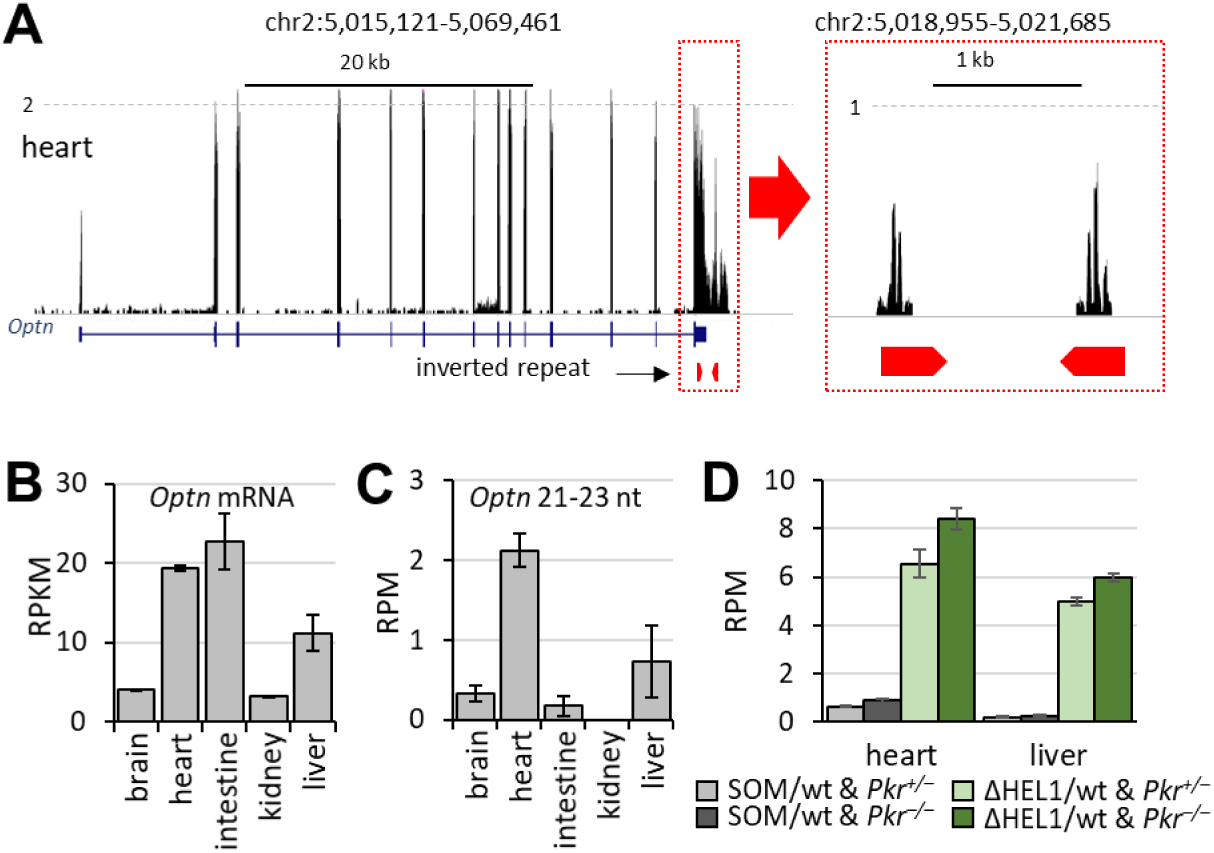
Analysis of endo-siRNAs from a natural inverted repeat at the 3’ end of *Optn* gene. (A) USCS browser snapshot of the *Optn* locus displaying mRNA expression in heart using published RNA-seq data (63). Next to it are displayed 21-23 nt RNAs from small RNA sequencing data from heart of a normal mouse (64) mapped into the *Optn* inverted repeat (depicted by red pentagons). The vertical scale is in counts per million 18-32 nt reads (CPM). (B) Quantification of *Optn* mRNA expression in organs based on a published dataset (63). The vertical scale is in reads per kilobase per million (RPKM). (C) Quantification of *Optn* endo-siRNAs in the same set of organs as in the panel B using a published RNA-seq dataset (64). (D) Effect of ΔHEL1 and *Pkr* deletion on endo-siRNA abundance in heart and liver. All error bars = SEM.

Because of the expected negligible amounts of naturally arising endo-siRNAs, we used the MosIR transgene (28), which is expressing long dsRNA hairpin from a strong synthetic promoter (Fig. 4A). The CAG promoter (35) yields different levels of transgene expression in different organs (Fig. 4B). This expression is consistent with previously observed expression of CAG-EGFP transgenes (28,36) and allows for investigating capacity of siRNA production and efficient induction of RNAi. Analysis of small RNAs from *Dicer^SOM/wt^* and *Dicer^ΔHEL1/wt^* animals carrying the MosIR transgene showed strong increase of MosIR-derived endo-siRNA in animals carrying a single *Dicer^ΔHEL^*allele (Fig. 4C). MosIR siRNA abundance in *Dicer^ΔHEL1/wt^* organs ranged over three orders of magnitude and reached > 10 000 RPMs in heart, which also had the highest MosIR expression, as indicated by the highest EGFP fluorescence in this organ (Fig. 4B) and qPCR analysis (Fig. 4D). This suggests that siRNA biogenesis in *Dicer^ΔHEL1/wt^* is primarily limited by dsRNA abundance and achieving high siRNA levels from endogenously expressed dsRNA requires very high expression: >10 000 RPM Mos siRNAs level remarkably contrasts with negligible abundance of *Optn* endo-siRNAs (Fig. 3C).

**Figure 4.**
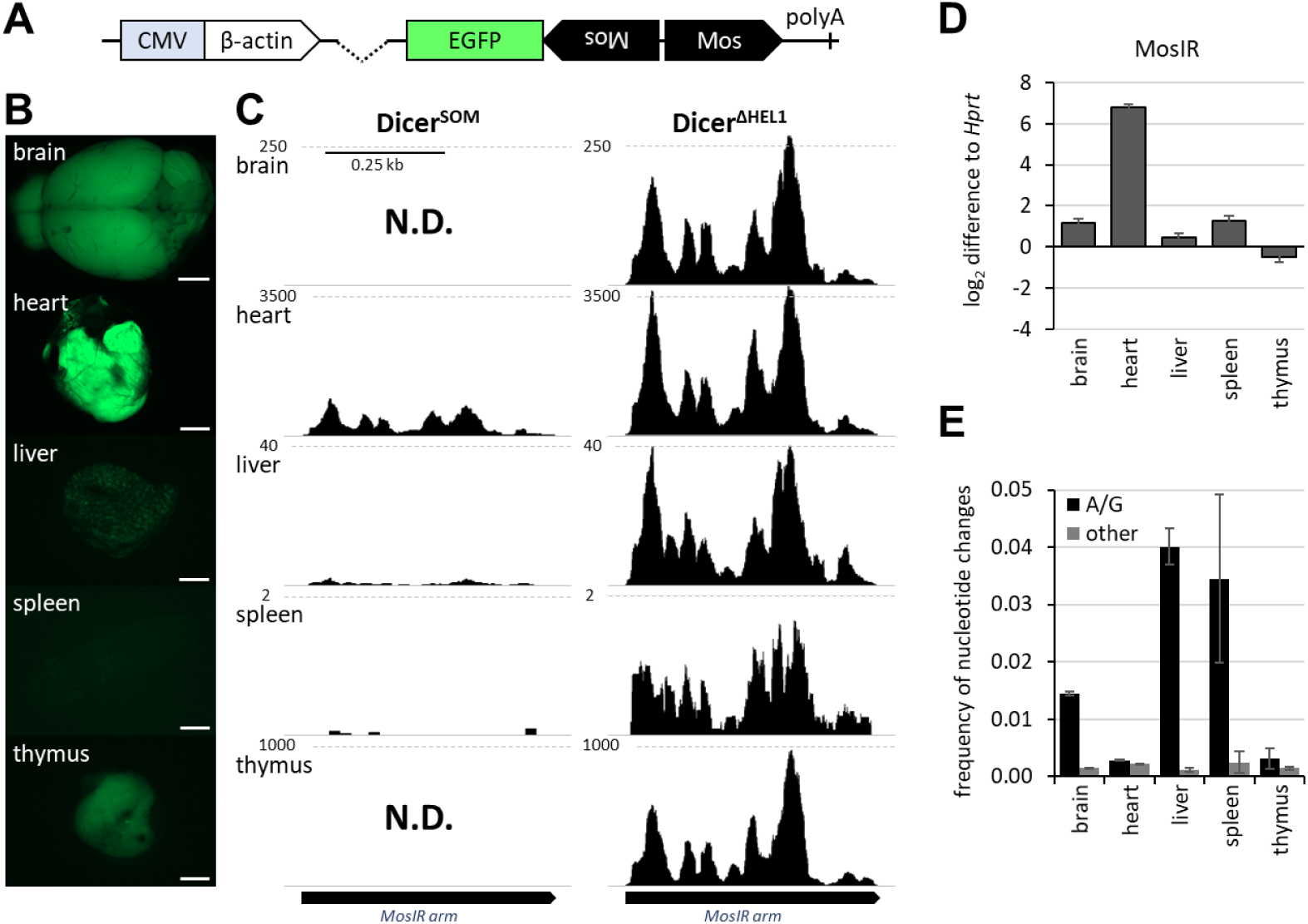
MosIR-derived small RNA production in SOM and ΔHEL1 heterozygotes. (A) Schematic depiction of the CAG-EGFP-MosIR transgene, a mouse transgenic line carrying this transgene has been established and analyzed previously (28). (B) EGFP fluorescence correlates with 21-23 nt MosIR small RNA abundance. Size bar = 2mm. (C) Abundance and distribution of 21-23 nt RNAs derived from MosIR and mapped onto endogenous Mos sequence. (D) qPCR analysis of CAG-EGFP-MosIR transgene expression in *Dicer^SOM/wt^ Pkr^−/−^* organs. Expression is presented as log_2_ relative difference to *Hprt* expression. (E) Analysis of mismatches in 21-23 nt small RNAs derived from MosIR in different organs. The black column represents frequencies of A-to-G change for all 21-23 nt small RNAs derived from MosIR in different organs. The grey column represents frequency of any other nucleotide changes in the same small RNA populations. All error bars = SD.

Notably, qPCR analysis of MosIR expression does not fully correlate with EGFP fluorescence and siRNA abundance (Fig. 4B and 4C vs. 4D). This could be in part explained by variability of *Hprt* expression used for normalization. In particular, RNA-seq data (37) suggest that *Hprt* transcript in spleen transcriptome is relatively less abundant than in brain, liver or thymus and this difference could cause that relative MosIR transcript levels would appear higher in spleen when normalized to *Hprt*. However, this does not explain an order of magnitude lower siRNA levels in spleen when compared to liver. Furthermore, MosIR in thymus appears to produce much higher fluorescence and MosIR siRNA levels than expected from qPCR analysis (Fig. 4B-D). In this case, RNA-seq data (37) suggest that *Hprt* transcript level in brain, liver and thymus transcriptomes is comparable and does not explain this discrepancy. Clearly, MosIR transcript levels, EGFP fluorescence, and MosIR siRNA abundance are influenced by different factors, which can cause observed discrepancies. For example, MosIR transcript level is influenced by CAG promoter activity and stability of the transcript, which can be affected by dsRNA response pathways such as RNA editing or cleavage by Dicer. EGFP fluorescence is influenced by the fraction of MosIR transcripts, which can be efficiently translated. MosIR siRNA abundance depends on the amount of the accessible transcript with the 3’ UTR folded into dsRNA.

One of the factors, which could influence siRNA biogenesis and can be partially investigated in RNA-seq data is adenosine deamination by adenosine deaminases acting on RNA (ADARs, reviewed in (38)). These enzymes convert adenosines in dsRNA into inosines, which appear in RNA sequencing data as A-to-G conversion. Hyper-editing was shown to antagonize RNAi (39). We thus analyzed mismatches of 21-23 nt long RNAs mapped to MosIR sequence with up to 10% mismatches. Under these conditions, ∼ 4% of all adenosines were changed to guanosines in MosIR sequences in liver and spleen while heart and thymus showed at least an order of magnitude lower A-to-G conversion (Fig. 4E). These data document that MosIR editing differs among analyzed organs and could affect siRNA biogenesis and activity.

Similarly to *Optn* endo-siRNAs, we did not observe any significant increase in *Mos* siRNAs in the absence of PKR in any of the analyzed organs (Fig. 5A). These data confirm that the inhibitory effect of PKR on siRNA biogenesis concerns transient transfections of MosIR into cells expressing Dicer^ΔHEL1^. Absence of increased siRNA levels in organs from *Dicer^ΔHEL1/wt^ Pkr^−/−^* Tg(CAG-EGFP-MosIR) animals argues against interference of PKR with dsRNA recognition and its cleavage by Dicer *in vivo*.

**Figure 5.**
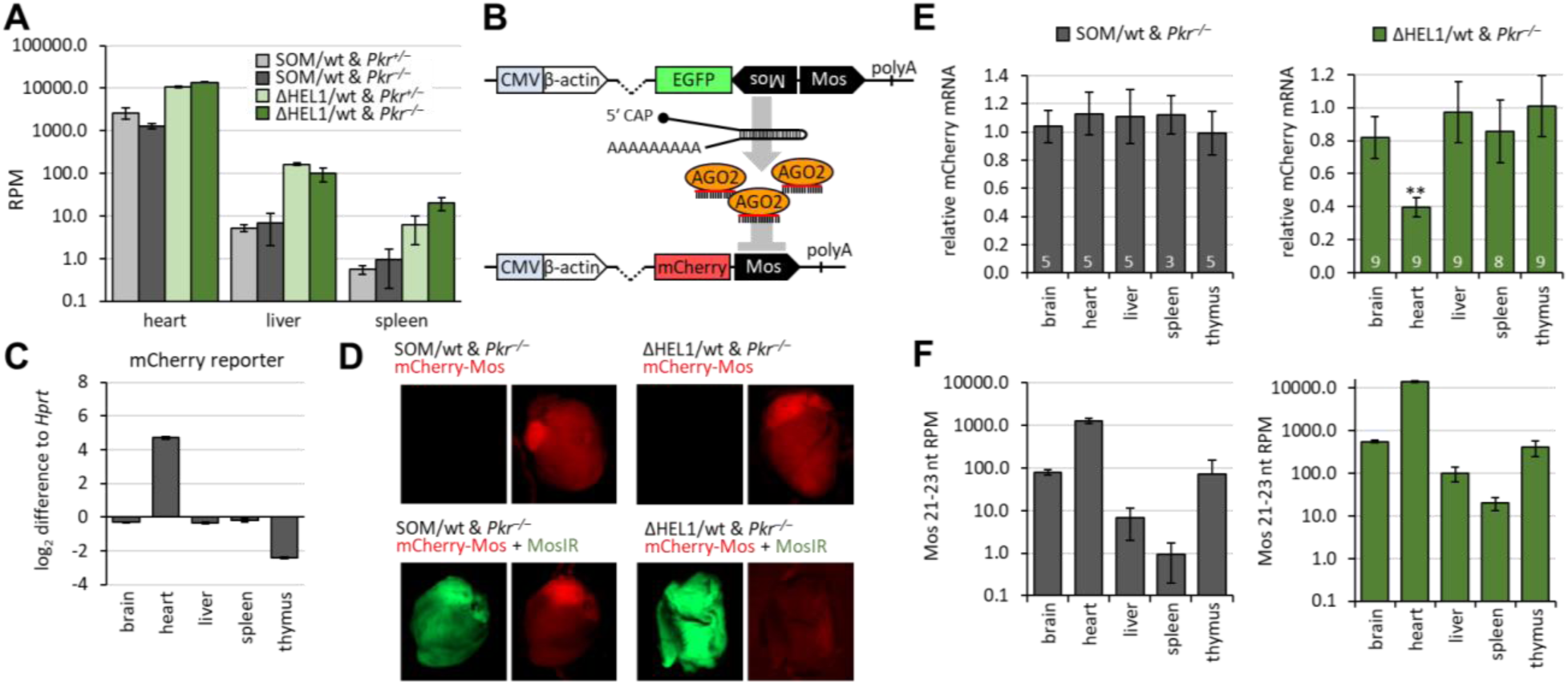
RNAi induction in SOM and ΔHEL1 heterozygotes. (A) Loss of PKR does not significantly enhance production of 21-23 nt RNAs from MosIR. (B) A scheme of the transgenic RNAi system using the transgene Tg(CAG-EGFP-MosIR) as a source of dsRNA and the transgene Tg(CAG-mCherry-Mos) as a sequence-specific target. (C) qPCR analysis of CAG-mCherry-Mos transgene expression in *Dicer^SOM/wt^ Pkr^−/−^* organs of a single animal in a triplicate technical replicate. Expression is presented as log_2_ relative difference to *Hprt* expression. (D) Fluorescence stereomicroscopy images of hearts in different genotypes showing GFP and RFP fluorescence. (E) qPCR analysis of mCherry-Mos levels relative to samples without the MosIR transgene. White numbers indicate the number of organ pairs used for comparison. Error bars = SEM, ** indicates statistically significant reduction of mCherry reporter mRNA level in ΔHEL1 heterozygotes relative to SOM heterozygotes (two-tailed t-test p-value = 0.0002). (F) MosIR endo-siRNA abundance – shown is 21-23 nt RNA abundance in reads per million (RPM) of all 18-32 nt mapped small RNA reads. Except of (E), all error bars = SD.

### Dicer^ΔHEL1^ can generate high siRNA levels needed for functional RNAi *in vivo*

To test whether *Dicer^ΔHEL1^* can support endogenous RNAi, we used MosIR as a source of dsRNA and, since *Mos* is not expressed in somatic tissues, we also produced a transgenic mouse line carrying an mCherry-Mos reporter whose expression is also controlled by the CAG promoter Tg(CAG-mCherry-Mos). This reporter should be sensitive to targeting by MosIR-derived endo-siRNAs (Fig 5B). The ratio of dsRNA to its target should be similar in different organs because MosIR and mCherry-Mos reporter rely on the same CAG promoter and their transcripts have comparable expression pattern across organs (Fig. 5C). To investigate RNAi effects, we crossed mice to combine MosIR and mCherry-Mos transgenes with *Dicer^ΔHEL1^* or *Dicer^SOM^* alleles and analyzed levels of the mCherry-Mos reporter in *Dicer^ΔHEL1/wt^* or *Dicer^SOM/wt^* mice in the presence or absence of MosIR. The analysis was done in the *Pkr^−/−^* background to avoid any possible interference with RNAi albeit we have ruled out that PKR significantly affects siRNA abundance.

Remarkably, mCherry fluorescence was reduced in hearts of *Dicer^ΔHEL1/wt^*mice in the presence of MosIR (Fig. 5D). Because fluorescence signal from the mCherry-Mos reporter in organs was not suitable for quantifying effects of MosIR, qPCR was used to estimate mCherry-Mos reporter targeting by RNAi in brain, heart, liver, spleen, and thymus. We have found that MosIR-induced mCherry-Mos reporter knock-down was present only in the heart of *Dicer^ΔHEL1/wt^*animals (Fig. 5E). RNAi effect thus occurred in the organ where MosIR siRNA abundance reached >10^4^ RPM (Fig. 5F). mCherry-Mos reporter mRNA was knocked down by 53%, but it should be noted that the target is overexpressed relative to *Hprt* (Fig. 5C). In the context of mammalian small RNA biology, >10^4^ RPM in our RNA-seq results is the level of expression of highly abundant miRNAs. Notably, MosIR siRNA abundance around 500 RPM observed in the brain and thymus of *Dicer^ΔHEL/wt^* mice (Fig. 5F) was insufficient to cause a detectable knockdown when the reporter was expressed similarly to (brain) or even several fold less (thymus) than *Hprt* (Fig. 5C). Particularly absence of RNAi in thymus where the reporter was much less expressed than in brain is informative regarding the siRNA abundance threshold for achieving functional RNAi *in vivo*.

Importantly, siRNA concentrations used in RNAi experiments in cultured cells (5-50 nM) would correspond to 10^4^-10^5^ siRNA molecules in 3 pl, which is approximately volume of a single fibroblast cell (40). Abundant miRNAs appear to fall into a similar range – it was estimated that a HeLa cell contains ∼ 50,000 let-7 miRNA molecules (41). This suggests that results from our RNAi experiments *in vivo* are rather in line with experiments in cell culture and that requirement for high abundance of siRNAs is an intrinsic feature of the mammalian RNAi, presumably because it is exploiting molecular mechanism primarily serving the miRNA pathway. In this context, our results provide an important framework for future research on mammalian canonical RNAi. We demonstrate that induction of functional canonical RNAi *in vivo* in mouse soma is possible because mice can tolerate both, increased Dicer activity and high expression of long dsRNA substrate. While these two conditions represent the two main constraints for activation of canonical RNAi *in vivo*, our results suggest that the main limiting factor in canonical RNAi is abundance of dsRNA, because a single allele of Dicer^ΔHEL1^ expressed from Dicer’s endogenous promoter was sufficient to support RNAi in heart highly expressing MosIR.

Since availability of dsRNA so strongly limits RNAi in mammalian cells, it has good predictive value when considering endogenous canonical RNAi in mammals. Genomic loci giving rise to perfectly complementary long dsRNA in somatic cells are typically rare and yield low-abundant siRNAs in somatic organs because the RNAi molecular machinery still recognizes dsRNA and makes AGO2-loaded siRNAs but there is no efficient target repression. dsRNA availability for RNAi is also affected by other molecular mechanisms recognizing dsRNA mammalian cell, such as the above-mentioned RNA editing, or interferon pathway. To efficiently function in mammalian cells, canonical RNAi thus requires adaptations bringing together necessary amounts of Dicer activity and dsRNA. Adaptations identified in mouse oocytes are a notable example of this scenario. They express Dicer^O^, have reduced RNA turnover (42), and suppressed interferon pathway (43). Under these conditions, stable lncRNAs carrying antisense pseudogenes base pair with mRNAs from parental genes and resulting dsRNA is efficiently processed by Dicer into siRNAs, which are able to significantly reduce specific transcript levels (33,34).

Apart from Dicer activity and dsRNA availability, additional factors could influence/restrict RNAi activity. One of the underexplored ones is composition of the AGO2 effector complex. As mammalian RNAi uses the molecular mechanism primarily dedicated to the miRNA pathway, its efficiency may be undermined in additional ways. First, siRNAs also act as miRNAs (44), i.e. they are loaded onto the miRNA-effector complex (miRISC), which is able to bind and repress partially complementary mRNAs through translational repression and destabilization. This is the molecular basis of the so-called siRNA off-targeting (reviewed in (45)). In siRNA transfections, off-targeting is a source of artifacts coming from sequence-specific suppression of partially complementary undesirable targets. In case of a mixture of siRNAs made from long dsRNA, the off-targeting effect is diluted but the transcriptome would function as a sponge for all partially complementary siRNAs and reduce probability of targeting perfectly complementary targets. As RNAi requires only small RNA bound to AGO2 (holoRISC), its efficiency could be increased by increasing holoRISC abundance by, for example, preventing assembly of the miRISC by disrupting interaction of AGO2 with the adaptor protein TNRC6, which provides a landing platform for additional miRISC components (46). Second, highly complementary targets may cause target-mediated miRNA degradation (TDMD) (47). This mechanism would affect efficiency of RNAi because it would reduce multiple turnover activity of siRNA-loaded AGO2. As TDMD operates through destabilization of AGO (48,49), it may constrain RNAi efficiency also at the level of AGO2 available for siRNA loading. TDMD thus could represent another limiting factor worth of investigation when researching potential of mammalian canonical RNAi.

Taken together, despite Dicer function became adapted to miRNA biogenesis early in vertebrate evolution (23,50), limited ability to induce RNAi remains preserved in mammals and can be boosted by a simple N-terminal truncation of Dicer or even full-length Dicer overexpression (15). However, we show *in vivo* using mouse genetic models that efficient RNAi in soma requires achieving considerably high abundance of siRNAs. This is consistent with the idea that RNAi employs molecular machinery, which evolved to primarily serve the miRNA pathway and constrains RNAi in several ways. Consequently, functional mammalian canonical RNAi likely occurs under distinct conditions, which overcome the aforementioned constrains. A distinct case may be participation of RNAi in mammalian antiviral immunity, which has been reported (51,52) but remains unclear. We would like to point out that antiviral RNAi would have two additional features, which experiments presented here lack. First, Dicer-mediated processing of dsRNA of replicating viruses could have an antiviral effect on its own. Second, we analyzed RNAi here using a system of steady-state expression of the trigger and target. This is a different scenario than targeting a replicating system where even a few percent reduction of its replication efficiency may have well-detectable effect after several cycles.

## Methods

### Animals

Tg(CAG-EGFP-MosIR), *Dicer^SOM^* and *Dicer^ΔHEL1^*mice were produced as previously described (23,24,28). The original mutation was introduced into a 129 strain ESC line, which gave rise to a chimeric male mice, which was mated with ICR females to obtain germline transmission. Once the line was established, breeding was maintained on ICR background as well mice were bred onto the C57Bl/6NCrl background for at least six generations. Tg(CAG-mCherry-Mos) and Pkr (Eif2ak2) mutant mice were produced for this study and their production is described further below. Animal experiments were carried out in accordance with the Czech law and were approved by the Institutional Animal Use and Care Committee (approval no. 34-2014).

### Genotyping

For genotyping, tail biopsies were lysed in DEP-25 DNA Extraction buffer (Top-Bio) according to the manufacturer’s instructions. An 1 μl aliquot was used with HighQu DNA polymerase master mix for PCR. Genotyping primers are provided in Table S1.

### Embryo and organ harvest

Mice were mated overnight, and the presence of a vaginal plug indicated embryonic day (E) 0.5. Mice were sacrificed by cervical dislocation. The embryos were collected at E15.5, washed in PBS and used for RNA sequencing. Organs collected from sacrificed animals were either directly used for analysis or stored in −80° C for later use.

### Tg(CAG-mCherry-Mos) mouse line

To generate the Tg(CAG-mCherry-Mos) mouse line, a PCR-amplified 750 bp fragment of the N-terminal part of the Mos transcript (corresponding to nucleotides 292–1025 of the Mos cDNA sequence ENSMUST00000105158.2) carrying the upstream EcoRI-NotI and downstream BglII sequence was inserted into the pCAG-EGFP plasmid (53) cleaved by EcoRI and BglII. The resulting pCAG-Mos plasmid was then digested by EcoRI and NotI, and the mCherry sequence carrying the upstream EcoRI and downstream NotI sequence was inserted to create an in-frame mCherry-Mos fusion. The final pCAG-mCherry-Mos plasmid was verified by restriction digestion and sequencing. SalI and HindIII were used to release the transgene cassette. The 3636 bp construct was gel purified using the QIAGEN gel extraction kit according to the manufacturer’s instructions, repurified using the QIAGEN PCR kit according to the manufacturer’s instructions, and diluted to 10 ng/µl in embryo-certified water. Transgenic mice were generated in the transgenic facility of the Czech Phenogenomics Center of the Institute of Molecular Genetics by injecting linearized DNA into the male pronuclei of C57BL/6 1-cell embryos. Transgene-positive mice were identified by PCR (primer sequences are shown in Table S1). Two founder animals were obtained, the lineage with better detectable mCherry fluorescence was expanded and used for experiments in this work.

### Pkr (Eif2ak2) mutant mice

The *Eif2ak2* (commonly known as *Pkr*) mutant model was produced in the Czech Centre for Phenogenomics at the Institute of Molecular Genetics ASCR using Cas9-mediated deletion of *Eif2ak2* exons 2 (containing the first AUG) to 5 coding for dsRNA-binding domains (amino acids: 1-165). Sequences of guide RNAs were Pi1B: 5’-GTGTTTCCAACCCACCAC<u>AGG</u> in the intron 1 and Pi5B: 5’-GGATCATTGTTGGTACAC<u>AGG</u> in the intron 5 (yielding 7,338-nt deletion). To produce guide RNAs, synthetic 128 nt guide DNA templates including T7 promoter, 18nt sgRNA and tracrRNA sequences were *in vitro* transcribed using the mMESSAGE mMACHINE T7 Transcription Kit (Ambion) and purified using the mirPremier™ microRNA Isolation Kit (Sigma). The Cas9 mRNA was *in vitro* transcribed from pSpCas9 plasmid (PX165; Addgene plasmid #48137) using Ambion mMESSAGE mMACHINE T7 Transcription Kit, and purified using the RNeasy Mini kit (Qiagen). A sample for microinjection was prepared by mixing two guide RNAs in water (25 ng/μl for each) together with Cas9 mRNA (100 ng/μl). Five picoliters of the mixture were microinjected into male pronuclei of C57Bl/6 zygotes and transferred into pseudo-pregnant recipient mice. PCR genotyping was performed on tail biopsies from four-weeks-old animals. A positive founder which transmitted the mutant allele to F_1_ was back-crossed with C57Bl/6NCrl animals for at least five generations before using in experiments.

Knock-out allele was detected using mPKR_i1_Fwd: 5′-GCCTTGTTTTGACCATAAATGCCG and mPKR_E6_Rev: 5′-GTGACAACGCTAGAGGATGTTCCG primers giving a 552 bp product (wild-type allele is too long to be amplified). Wild-type allele was detected using mPKR_i1_Fwd and mPKR_E2_gen_Rev: 5′-TGGCTACTCCGTGCATCTGG primers yielding a 404 bp product.

### Dicer^ΔHEL1/wt^ phenotype analysis

The phenotype analysis of animals with the ICR outbred background at the Czech Centre for Phenogenomics utilized the established phenotyping pipeline (https://mousephenotype.org/impress/index). The full report including description of the tests is provided in the Supplementary File S1.

### Plasmids for RNAi assay

Plasmids for RNAi assay were introduced in detail in our previous work (15). Briefly, three dsRNA-expressing plasmids (MosIR, Lin28IR, and Elavl2IR) use inverted repeats, which efficiently induced RNAi in oocytes of transgenic mice (54–56). In each transfection, one of these plasmids is combined with a non-targeted firefly luciferase reporter pGL4-SV40 (Promega; for simplicity referred to as FL) and a *Renilla* luciferase reporter carrying a Mos cognate sequence complementary to MosIR-expressed dsRNA. All non-commercial plasmids are available from Adgene with details about their construction and sequence. Plasmid sequences (annotated Genbank format) were provided previously (15).

### Cell culture and transfection

Mouse ESCs were cultured in 2i-LIF media: KO-DMEM (Gibco) supplemented with 15% fetal calf serum (Sigma), 1x L-Glutamine (Thermo Fisher Scientific), 1x non-essential amino acids (Thermo Fisher Scientific), 50 µM β-Mercaptoethanol (Gibco), 1000 U/mL LIF (Isokine), 1 µM PD0325901, 3 µM CHIR99021 (Selleck Chemicals), penicillin (100 U/mL), and streptomycin (100 µg/mL). For transfection, cells were plated on a 24-well plate, grown to 50 % density and transfected using Lipofectamine 3000 (Thermo Fisher Scientific) according to the manufacturer’s protocol. Cells were co-transfected with 100 ng of each FL and RL reporter plasmids, 400 ng of a dsRNA-expressing plasmid and, eventually, 400 ng of a plasmid expressing a tested factor per well. The total amount of transfected DNA was kept constant (1 μg/well) using pBluescript stuffer. Cells were collected for analysis 48 hours post-transfection.

### Luciferase assay

Dual luciferase activity was measured according to Hampf M. & Gossen M. (57) with some modifications. Briefly, cells were washed with PBS and lysed in PPTB lysis buffer (0.2% v/v Triton X-100 in 100 mM potassium phosphate buffer, pH 7.8). A 3-5 µl aliquots were used for measurement in 96-well plates using Modulus Microplate Multimode Reader (Turner Biosystems). First, firefly luciferase activity was measured by adding 50 µl substrate (20 mM Tricine, 1.07 mM (MgCO_3_)_4_·Mg(OH)_2_·5H_2_O, 2.67 mM MgSO_4_, 0.1 mM EDTA, 33.3 mM DTT, 0.27 mM Coenzyme A, 0.53 mM ATP, 0.47 mM D-Luciferin, pH 7.8) and signal was integrated for 10 sec after 2 sec delay. Signal was quenched by adding 50 µl Renilla substrate (25 mM Na_4_PP_i_, 10 mM Na-Acetate, 15 mM EDTA, 500 mM Na_2_SO_4_, 500 mM NaCl, 1.3 mM NaN_3_, 4 µM Coelenterazine, pH 5.0) and *Renilla* luciferase activity was measured for 10 sec after 2 sec delay.

### Microscopy

Mice organs were harvested, rinsed with ice-cold PBS and placed on ice. Organ images were acquired on a Zeiss Axio Zoom.V16 stereo microscope equipped with a Zeiss Axiocam 512 mono camera and analyzed with the Zeiss Zen 2.5 lite software. 1x 0.25 NA objective, the fluorescence filter sets GFP (excitation wavelength: 488 / emission wavelength: 509) + RFP (excitation wavelength: 590 / emission wavelength: 612) and the Bright Field optics were used for image acquisition. Background fluorescence levels for each organ were determined using WT littermates.

### qPCR

For qPCR analysis, organs from adult mice (9-13 weeks old) were harvested and homogenized in Qiazol lysis reagent (Qiagen) and total RNA was isolated by phenol–chloroform extraction according to the manufacturer’s protocol. 1 µg of total RNA was reverse transcribed using LunaScript RT SuperMix Kit (New England Biolabs) according to the manufacturer’s instructions and 1 µl cDNA was used as a template for a 10 µl qPCR reaction. qPCR was performed on LightCycler 480 (Roche) and the Maxima SYBR Green qPCR master mix (Thermo Fisher Scientific) was used for the qPCR reaction. qPCR was performed in technical triplicates for each biological sample. Average Ct values of the technical replicates were normalized to the housekeeping genes *Hprt*, *B2m* and *Alas1* using the ΔΔCt method. A list of the primers used for qPCR is provided in Supplementary Table S1.

### RNA sequencing

#### ESC small RNA sequencing (RNA-seq)

Cells were plated on 6-well plates and grown to 80 % density. Cells were transfected with 2 μg/well of pCAG-EGFP-MosIR plasmid and cultured for 48 hours. Cells were washed with PBS, homogenized in Qiazol lysis reagent (Qiagen) and total RNA was isolated by Qiazol-chloroform extraction and ethanol precipitation method (58). RNA quality was verified by Agilent 2100 Bioanalyzer. Small RNA libraries were constructed using NEBNext Multiplex Small RNA Library Prep Set for Illumina (New England Biolabs) according to the manufacturer’s protocol. Small RNA libraries were size selected on 6% PAGE gel, a band of 140 - 150 bp was cut from the gel and RNA was extracted using Monarch^®^ Genomic DNA Purification Kit. Quality of the libraries was assessed by Agilent 2100 bioanalyzer. Libraries were sequenced on the Illumina HiSeq2000 platform at the Genomics Core Facility at EMBL.

#### E15.5 small RNA-seq

E15.5 embryos were removed from the uterus and washed in PBS. The yolk sac was taken for genotyping and embryos were transferred into RNAlater (Thermo Fisher Scientific). Embryos were homogenized in Qiazol lysis reagent (Qiagen) and total RNA was isolated by Qiazol-chloroform extraction and ethanol precipitation method (58). Small RNA libraries were constructed using Nextflex Small RNA-seq kit v3 for Illumina (Perkin Elmer) according to the manufacturer’s protocol; 3′ adapter ligation was performed overnight at 20 °C, 15 cycles were used for PCR amplification and NextFlex beads were used for size selection. Final libraries were sequenced by 75-nucleotide single-end reading using the Illumina NextSeq500/550 platform at the core genomics facility of IMG.

#### Adult organ small RNA-seq

For small-RNA-seq analysis, organs from adult mice (9-13 weeks old) were harvested and homogenized in Qiazol lysis reagent (Qiagen) and total RNA was isolated by phenol–chloroform extraction according to the manufacturer’s protocol. Small-RNA libraries were prepared using the NextFlex Small-RNA-seq v3 kit (Amplicon) according to the manufacturer’s protocol; 3′ adapter ligation was performed overnight at 20 °C, 15-18 cycles were used for PCR amplification and gel purification was performed for size selection. For gel purification, libraries were separated on a 2.5% agarose gel using 1× lithium borate buffer and visualized with ethidium bromide. The 140-160 bp fraction was cut off the gel and DNA was isolated using the MinElute Gel Extraction Kit (Qiagen). Final libraries were sequenced by 75-nucleotide single-end reading using the Illumina NextSeq500/550 platform at the core genomics facility of IMG.

The list of small RNA-seq libraries from adult brain, heart, liver, spleen and thymus is in the Table S2. Next to age-matched wild type controls (C57Bl/6NCrl background), we sequenced four distinct genotypes on the C57Bl/6NCrl background, which included a transgene Tg(CAG-EGFP-MosIR) and one of the four genotypes: *Dicer^SOM/wt^ Pkr^+/–^, Dicer^SOM/wt^ Pkr^−/−^*, *Dicer^ΔHEL1/wt^ Pkr^+/–^, Dicer ^ΔHEL1/wt^ Pkr^−/−^*. *Dicer^SOM/wt^* animals provided another control for phenotype analyses because the *Dicer^SOM^* allele has the same sequence as the *Dicer^ΔHEL1^* allele except of the HEL1 sequence.

### Bioinformatic analyses

RNA-seq data (Table S2) were deposited in the Gene Expression Omnibus database under accession numbers GSE243016 (reviewer access token: svgpsgamxvcjrad) and GSE242871 (reviewer access token: gjuhyqeglfsnvgh)

### Mapping of small RNA-seq data

Small RNA-seq reads were trimmed in two rounds using fastx-toolkit version 0.0.14 (http://hannonlab.cshl.edu/fastx_toolkit) and cutadapt version 1.8.3 (59). First, 4 random bases were trimmed from left side:

`fastx_trimmer -f 5 -i {INP}.fastq -o {TMP}.fastq`

Next, NEXTflex adapters were trimmed. Additionally, the N-nucleotides on ends of reads were trimmed and reads containing more than 10% of the N-nucleotides were discarded:

cutadapt --format=“fastq” --front=”GTTCAGAGTTCTACAGTCCGACGATCNNNN” -- adapter=”NNNNTGGAATTCTCGGGTGCCAAGG” --error-rate=0.075 --times=2 -- overlap=14 --minimum-length=12 --max-n=0.1 --output=”${TRIMMED}.fastq” -- trim-n --match-read-wildcards ${TMP}.fastq

Trimmed reads were mapped to the mouse (mm10) genome with following parameters:

STAR --readFilesIn ${TRIMMED}.fastq.gz --runThreadN 4 --genomeDir ${GENOME_INDEX} --genomeLoad LoadAndRemove --readFilesCommand unpigz -c -- readStrand Unstranded --limitBAMsortRAM 20000000000 --outFileNamePrefix ${FILENAME} --outReadsUnmapped Fastx --outSAMtype BAM SortedByCoordinate -- outFilterMultimapNmax 99999 --outFilterMismatchNoverLmax 0.1 -- outFilterMatchNminOverLread 0.66 --alignSJoverhangMin 999 -- alignSJDBoverhangMin 999

### miRNA expression analyses

Mapped reads were counted using program featureCounts (60). Only reads with lengths 19-25nt were selected from the small RNA-seq data:

`featureCounts -a ${ANNOTATION_FILE} -F ${FILE} -minOverlap 15 -fracOverlap 0.00-s 1 -M -O -fraction -T 8 ${FILE}.bam`

The GENCODE gene set (37) was used for the annotation of long RNA-seq data. For small RNA-seq data analysis, the miRBase 22.1. (61) miRNA annotation was combined with published mirtron annotation (25). Small RNA expression was analyzed as described previously (23). Statistical significance and fold changes in gene expression were computed in R using the DESeq2 package (62). Genes were considered to be significantly up- or down-regulated if their corresponding p-adjusted values were smaller than 0.05. The DESeq2 baseMean and fold changes were plotted and visualized by home-made R scripts as described previously (23).

### Small RNA editing and tailing analysis

Reads were mapped to MosIR with up to 10% of mismatches. Reads with mismatches were then extracted and classified by a homemade script.

## Acknowledgements

We thank Vladimir Benes and EMBL sequencing facility for help with RNA-seq experiments, and Kristian Vlahovicek for providing hardware support for bioinformatics analyses. Development of *Dicer^SOM^* and *Dicer^ΔHEL1^*mouse models was funded from the European Research Council under the European Union’s Horizon 2020 research and innovation programme (grant agreement No 647403, D-FENS). Main funding was provided by the Czech Science Foundation EXPRO grant 20-03950X. Additional support was provided by the Ministry of Education, Youth, and Sports (MEYS) project NPU1 LO1419. Financial support of V.B., F.H., and M.K. was in part provided by the Charles University in a form of a PhD student fellowship; this work will be in part used to fulfil requirements for a PhD degree and hence can be considered “school work”. The authors used services of the Czech Centre for Phenogenomics at the Institute of Molecular Genetics supported by the Czech Academy of Sciences RVO 68378050 and by the project LM2018126 and LM2023036 Czech Centre for Phenogenomics provided by Ministry of Education, Youth and Sports (MEYS)of the Czech Republic. We also acknowledge services of the Light Microscopy Core Facility, IMG, Prague, Czech Republic, supported by MEYS – LM2023050 and RVO – 68378050-KAV-NPUI. Computational resources were provided by the e-INFRA CZ project (ID:90254), supported by the Ministry of Education, Youth and Sports of the Czech Republic and by the ELIXIR-CZ project (ID:90255), part of the international ELIXIR infrastructure.

## Author contributions

Conceptualization: PS, RM

Data curation: VB, ES, JPa, FH, JPr, RM, PS

Formal analysis: VB, JPa, RM, JPr, PS Funding acquisition: RS, PS

Investigation: VB, JPa, RM, ZL, ET, FH, MIRK, IJ, JPr

Project administration: RM, JPr, RS, PS

Resources: JPr, RS Software: JPa, FH

Supervision: RM, JPr, PS, RS

Visualization: VB, JPa, JPr

Validation: PS, RM, JPr Writing – original draft: PS

Writing – review & editing: VB, JPa, RM, ZL, ET, FH, MIRK, IJ, JPr, RS, PS

## Data Availability

RNA sequencing data were deposited to Gene Expression Omnibus (GEO) with the following accession numbers GSE243016 and GSE242871. The full-list of samples is provided in the Table S2. Original codes were deposited at https://github.com/fhorvat/2023.RNAi_activation. Any additional information required to reanalyze the data reported in this paper is available from the lead contact upon request.

## SUPPLEMENT

**Figure S1.**
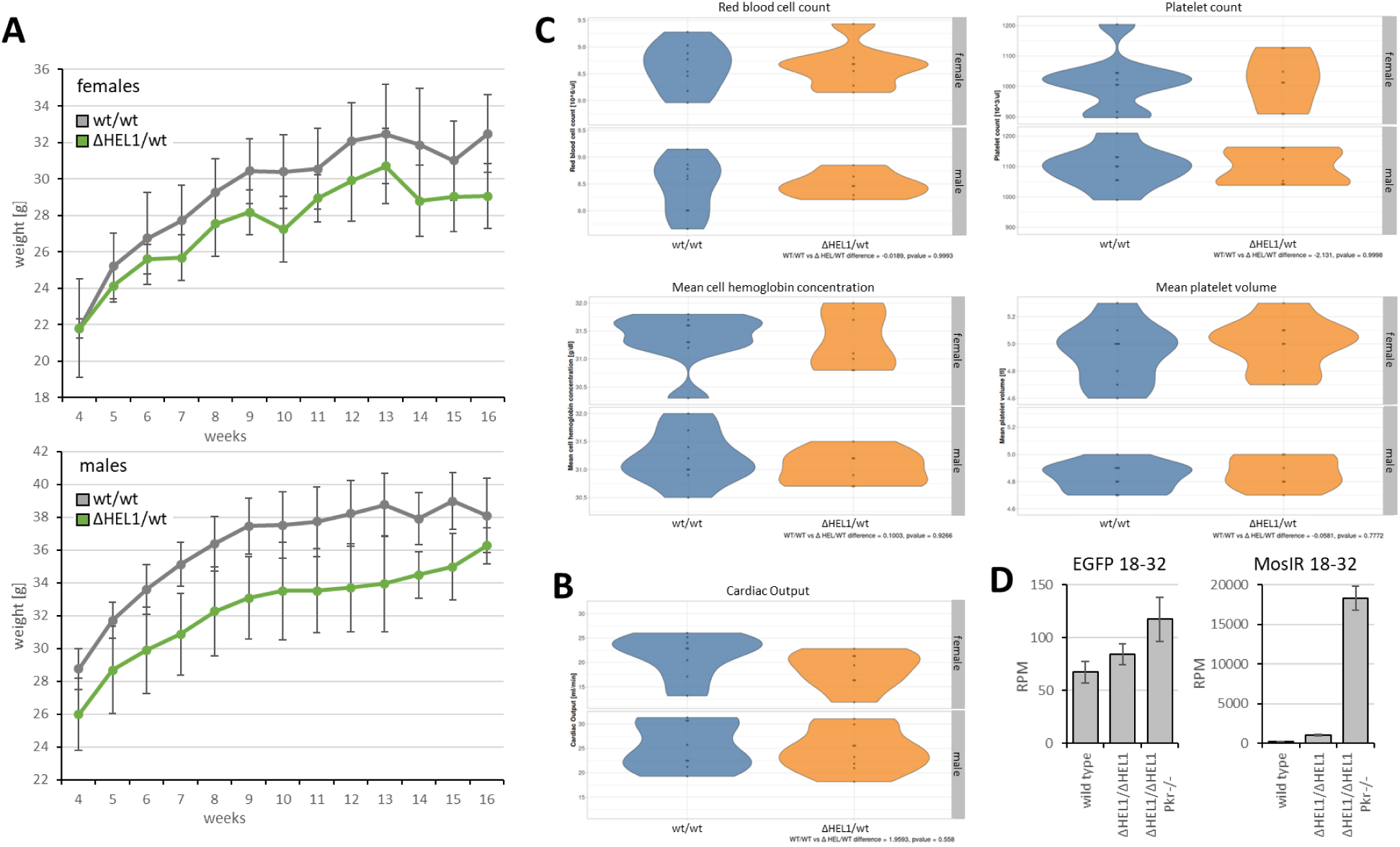
Selected phenotype features. (A) Growth curves of males and females on the ICR background. (B) *Dicer^ΔHEL1/wt^*animals have normal cardiac output. (C) Blood parameters disrupted in *Dicer^ΔHEL1/ΔHEL1^* (23) are normal in *Dicer^ΔHEL1/wt^* mice. (D) Analysis of abundance of EGFP and MosIR 18-32 nt RNA fragments in different ESCs lines.

**Figure S2.**
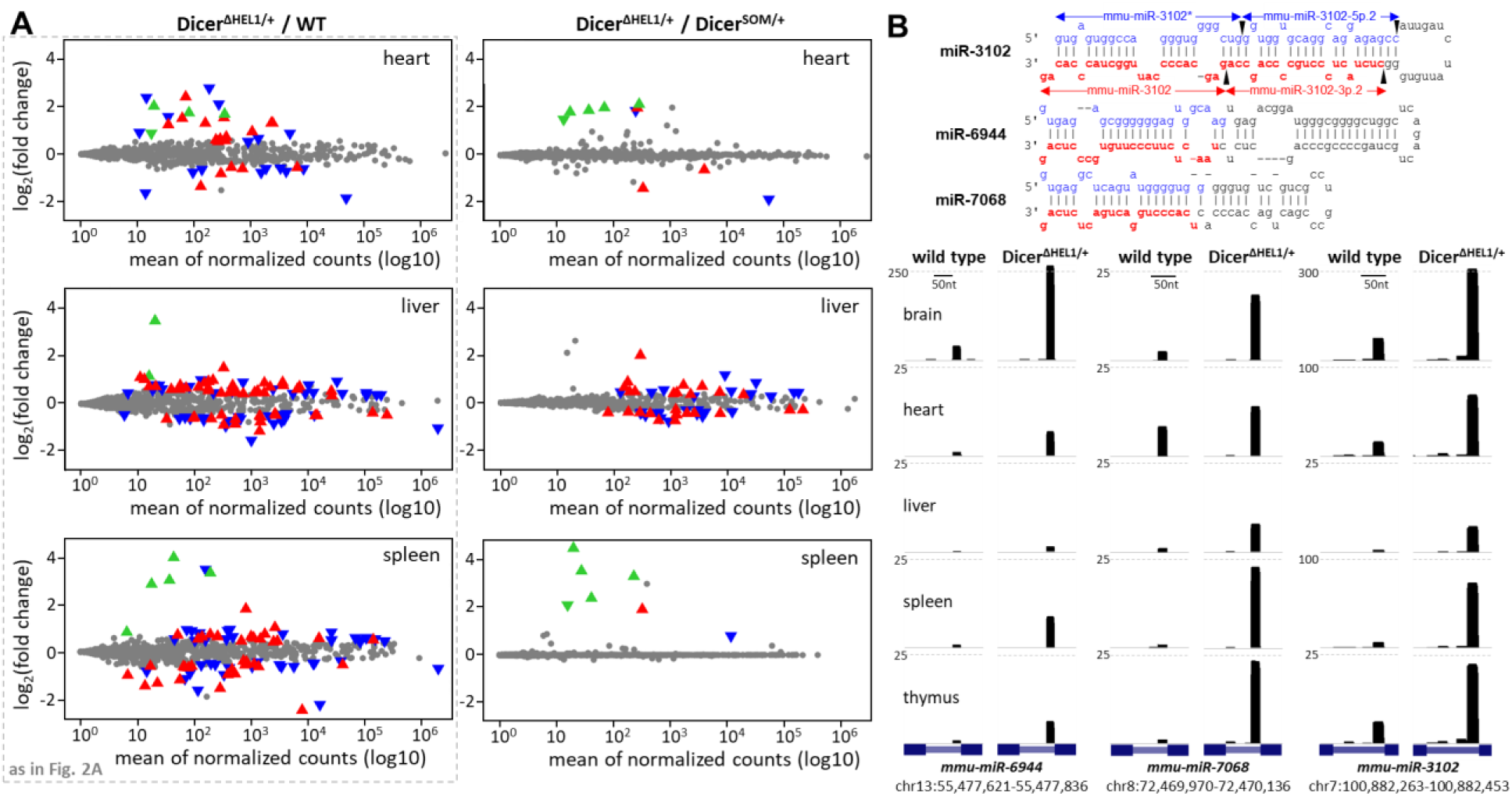
Supplementary data for miRNome changes. (A) MA plots depicting changes in levels of annotated murine miRNAs (miRBase 22.1 (23)) in *Dicer^ΔHEL1^*^/wt^ *Pkr*^+/–^ Tg(CAG-EGFP-MosIR) animals relatively to age-matched *Dicer^SOM^*^/wt^ *Pkr*^+/–^ Tg(CAG-EGFP-MosIR). Significantly dysregulated 5p and 3p miRNAs (DESeq p-value 0.05) are shown as oriented blue ▾and red ▴ triangles, respectively. Significantly dysregulated mirtrons are represented by green triangles whose orientation indicates 5p and 3p miRNA strands. (B) Most upregulated mirtrons. Mirtron precursor schemes were adapted from structures presented in miRBase (61). Below are UCSC browser snapshots showing abundance of 21-23 nt reads in mirtron loci in normal and ΔHEL1 mice. The vertical scale is in counts per million (CPM) of 18-32 nt reads.

**Figure S3.**
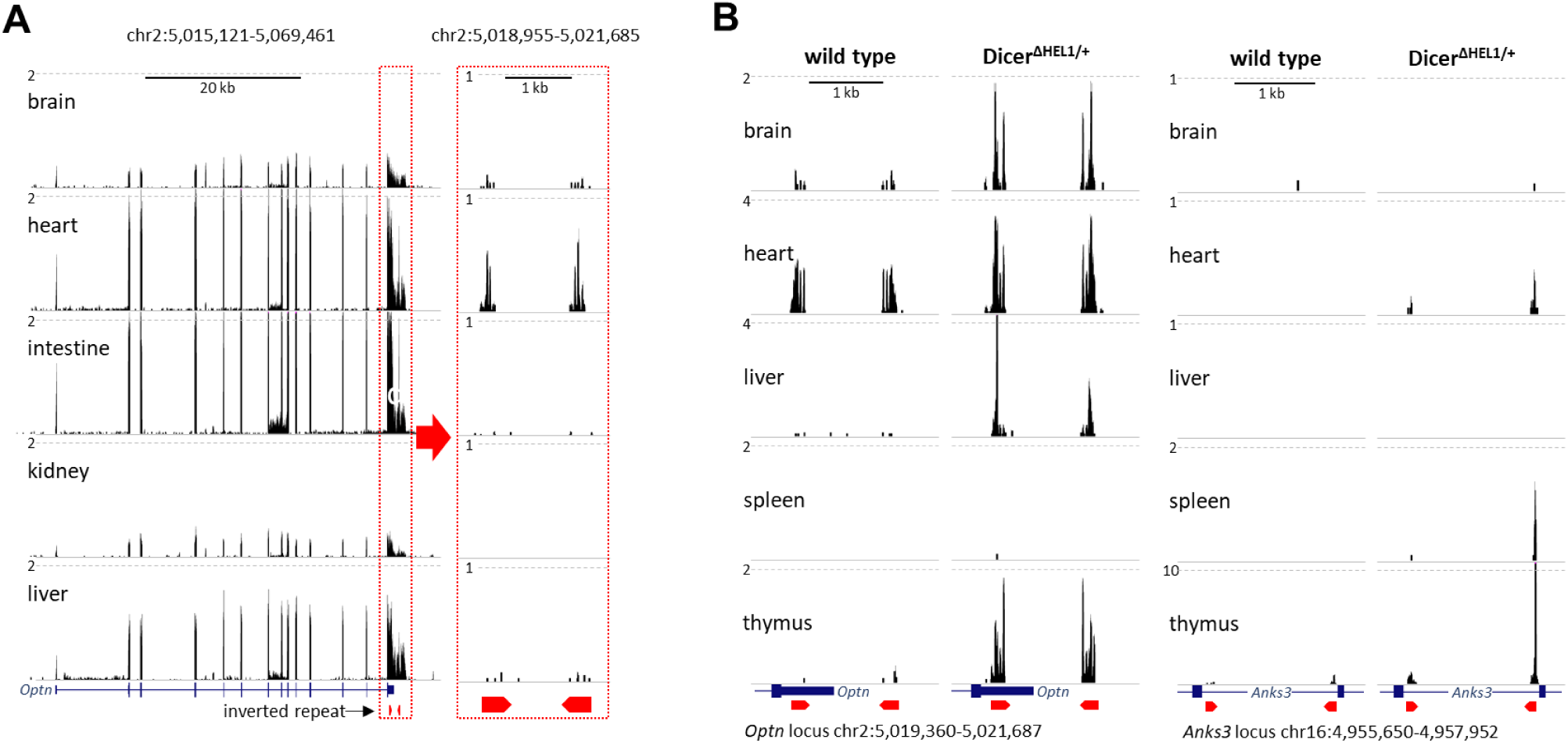
Supplementary data for endo-siRNA analysis. (A) *Optn* expression and siRNA production in different organs in wild type mice. On the left side is a UCSC browser snapshot of *Optn* transcript expression in different organs using publicly available RNA-seq libraries from different organs (63). The vertical scale is counts per million of reads (CPM). Next to it are 21-23 nt RNAs from small RNA sequencing data from the same organs (64) mapped into the *Optn* inverted repeat region (red pentagons). The vertical scale is in counts per million of 19-32 nt reads. (B) UCSC browser snapshots showing 21-23 nt reads in *Optn* and *Anks3* loci in normal and ΔHEL1 samples (*Dicer^ΔHEL1^*^/wt^ *Pkr*^+/–^ Tg(CAG-EGFP-MosIR)).

**Table S1.**
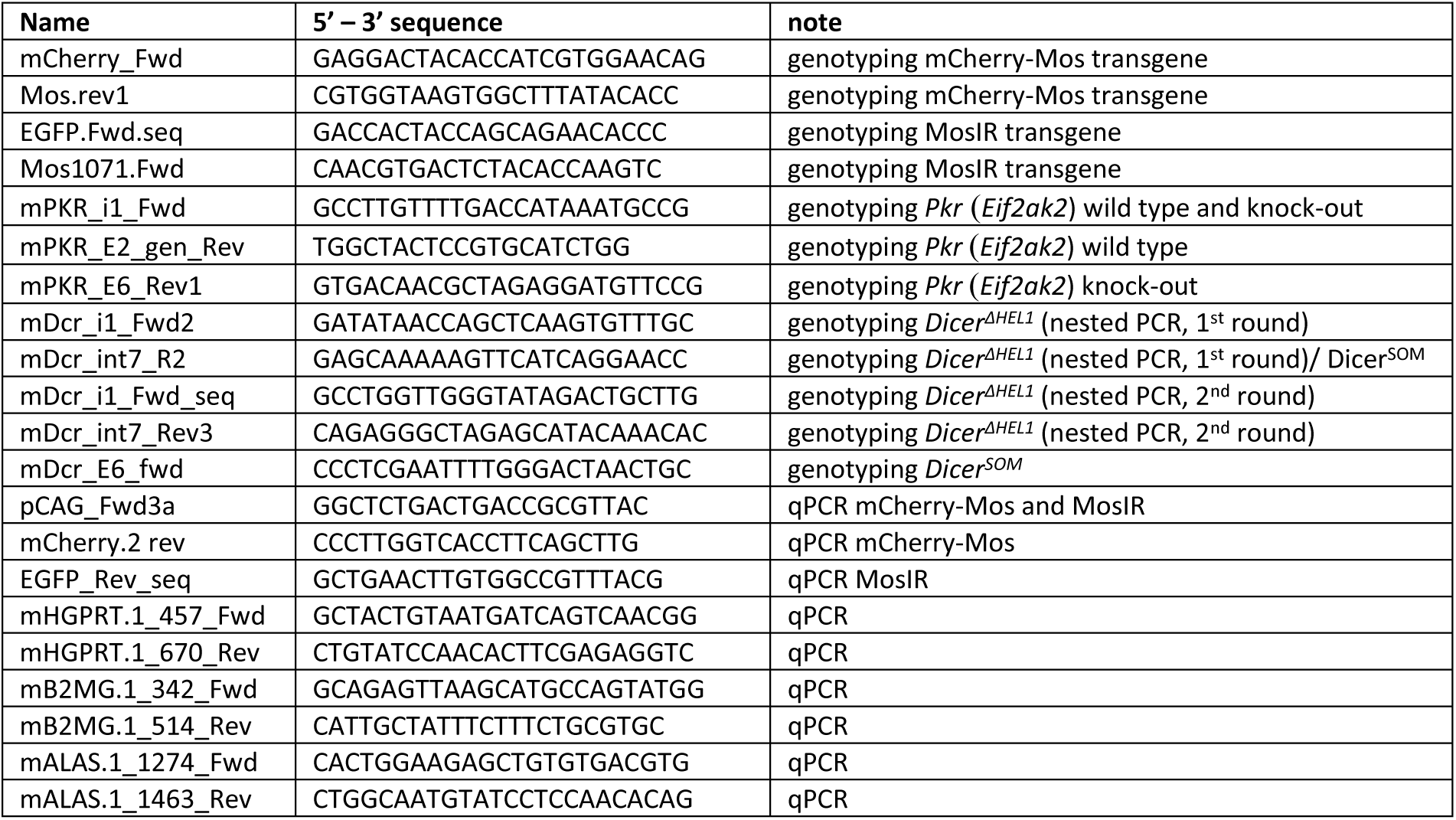
Primers.

Table S2 RNA-seq samples used in the study

Supplementary File S1 A complete *Dicer^ΔHEL1^*^/wt^ phenotyping report (html file)

