## Supplementary File 1 for "Genetic activation of canonical RNA interference in mice": 2023 Buccheri Supplementary File S1-2.htm

\Delta Hel1


### \(\Delta\) Hel1

The Czech Centre for Phenogenomics (CCP) is a large research
infrastructure dedicated to providing; first-class expertise, tools, and
services to reveal gene functions in human diseases, identifying new
therapeutic targets for treating human diseases and enhancing the
understanding of the genetic bases for human diseases. CCP was
established in 2015 as part of the wider Biotechnology and Biomedicine
Center of the Academy of Sciences and Charles University in Vestec
(BIOCEV) project and is hosted by the Institute of Molecular Genetics of
the ASCR, v. v. i. Located in the municipality of Vestec, Czech
Republic, CCP provides open access, state of the art services for both
the national and international research community. As a member of two
large consortia (IMPC and INFRAFRONTIER), CCP is committed to
discovering and ascribing biological function to each gene through the
generation of mutant mouse lines and use of the broad based primary
phenotyping pipeline which includes all the major adult organ
systems.

**Director** : Radislav Sedlacek, Assoc. Prof., PhD

**Phenotyping Module Head**: Jan Prochazka, PhD

Czech Centre for Phenogenomics, BIOCEV (IMG), Prumyslova 595, 252 50
Vestec, Czech Republic

### 1 Primary Screening at CCP

#### 1.1 Staff Members

##### Biochemistry and Hematology

**Head:** Roldan Medina De Guia

**Technician(s):** Mariya Glushchenko, Eva Štefancová,
Yu-chieh Wu

##### Bioimaging and Embryology

**Head:** Jan Prochazka

**Specialist:** Frantisek Spoutil

**Technician(s):** Michaela Prochazkova, Sarah Clewel,
Ivana Uramova, Tereza Michalcikova and Marie Kleisnerova

##### Cardiovascular

**Head:** Jiri Lindovsky

**Technician(s):** Petr Macek, Sarka Karbanova

##### Hearing

**Head:** Jiri Lindovsky

**Technician(s):** Miles J. Raishbrook, Jan Majernik

##### Histopathology

**Head:** Dagmar Zudova,

**Specialist(s):** Barbora Pavlu

**Technician(s):** Linda Kutlíková, Christine Sophia
Carl Cañada, Aneta Cestrová,Thuy Lien Duongová, Attila Juhász

##### Immunology

**Head:** Jana Balounova

**Specialist(s):** Kristína Vičíková

**Technician(s):** Kristyna Kunclova, Michaela
Simova

##### Lung function

**Head:**

**Technician(s):** Vaclav Zatecka

##### Metabolism

**Head:** David Pajuelo Reguera

**Technician(s):** Rajasree Sain

##### Neurobiology and Behaviour

**Head:** Agnieszka Kubik-Zahorodna

**Technician(s):** Pavel Jina, Rozalie Novakova,
Katarina Kanasova, Pavlina Kucerova

##### Vision

**Head:** Marcela Palkova

**Specialist:** Jiri Lindovsky

**Technician(s):** Viktoriia Symkina

##### Bioinformatics

**Head:** Vendula Novosadova

**Specialist:** Julia Potip, Frantisek Malinka and
Carlos Trufen

#### 1.2 Animal Care

At CCP, mice are housed in GM500 cages in individually and digital
ventilated caging (IVC and DVC) systems (IVC and DVC System Green Line,
Tecniplast, Italy) on aspen bedding (Tapvei, Estonia). The IVCs operate
with positive pressure. Mice are transferred to new cages at 2 week
intervals in Laminar Flow Class II changing stations (Tecniplast,
Italy). Mice are fed with autoclaved standard rodent high energy
breeding diet (Altromin 1314 Forti) and given reverse osmosis filtered
and chlorinated water ad libitum. Light is adjusted to a 12h/12h
light/dark cycle; temperature and relative humidity are maintained at
22±1 °C and 55±5 %, respectively. In specified modules husbandry
conditions are adjusted according to the experiment requirements. All
materials entering the breeding and experimental area are decontaminated
by autoclaves or by Vaporized hydrogen peroxide (VHP). All persons
entering the facility change their clothes (autoclaved jackets, trousers
and socks) and shoes, wear caps, masks and gloves before entering the
CCP facility. Entry into the facility is by wet or air shower depending
on the barrier status of the designated room.

Outbred 6-week-old SPF CD 1 mice are used as sentinels and kept on a
mixture of soiled bedding from all cages of the IVC racks connected to
one IVC blower. Health monitoring is carried out by outsourced service
of the sentinel mice by certified laboratories according to FELASA
recommendations http://www.felasa.org.

All animals are kept according to Directive 2010/63/EU and Czech
legislation.

#### 1.3 Quality Assurance

All experiments and investigations carried out at the Czech Centre
for Phenogenomics (CCP) are conducted according to standard operating
procedures (SOPs) and with approval of the IMG ethics committee.

All data generated is validated by CCP’s expert scientists who check
the biological and mathematical relevance of the data.

The constant development of staff is a vital part of CCP’s quality
assurance. To this end, CCP staff members are cross trained in multiple
techniques and regularly attend workshops, seminars and conferences to
further develop their skills and knowledge.

#### 1.4 Workflow

The Czech Centre for Phenogenomics standard primary workflow follows
the international mouse phenotyping consortium (IMPC) pipeline.

#### 1.5 Statistical Analysis

All numerical univariate data are analyzed using generalised mixed
model. The fixed variables are always Genotype, Gender, in the case that
data are dependent on weight the other fixed variable is weight, for
time series data the last fixed variable is time. The random variables
are Date and Experimenter. Categorical data are analyzed by Fisher test
with grouping WildType Female, WildType Male, Knockout Female, Knockout
Male.

### 2 Standard Screening Procedures

#### 2.1 Viability and Fertility

Purpose of these tests is to assess the fertility of homozygous
knockout mice and the postnatal viability, sub-viability, and lethality
of homozygous mice during cohort production.

For fertility screen homozygous mice (minimum age of 8 weeks, maximum
age of 14 weeks) are mated for 4-6 weeks. Strains that produce no
progeny or pregnant dams after 4 to 6 weeks progress to secondary
screening.

For viability screen mice are monitored for their genotypes after Het
X Het breeding. At least 28 pups are genotyped. Then strains which
produce no homo/hemizygotes are considered as **lethal**.
Strains that produce less than normal numbers of homozygous/hemizygous
male or female pups are considered as **subviable**.

#### 2.2 Body Weight

The body weight of each mouse is to be measured weekly, from the 4th
until the 16th week. Optional additional weighs may also be
recorded.

#### 2.3 Neurobiology and Behaviour

Neurobehavioral tests using transgenic animal models make it possible
to understand genetic mechanisms underlying neurological and psychiatric
disorders including, but not limited to, anxiety, schizophrenia, mood
disorders, and Parkinson’s disease. To understand the mechanisms
underlying these disorders we employ a number of tests which analyse
motor abilities, cognitive functions, emotion, sensory processing as
well as neurological, gait, auditory, and vision screens of rodents.

All animals are adapted to the testing room for 1hr prior to testing.
Before placement of an animal in any test, the equipment is cleaned with
alcohol to remove olfactory traces.

##### 2.3.1 Animal emotionality and affect

To study animal emotionality and affect we put into service tests
like Open Field, Elevated Plus Maze, Light/Dark box, Tail Suspension
Test, and Forced Swim Test.

**Open Field**

The open Field test is conducted according to IMPC
protocol and evaluates animal motility triggered by exploratory
drive in a new environment (Welker 1959). It is also used as an initial
screen for general anxiety elicited in a well-lit, open unprotected
space (Whimbey & Denenberg 1967). Detailed and fully automated
analysis of animal behaviour in open space is based on video tracking
system (Viewer, Biobserve GmbH). The software distinguishes an animal as
an object contrasting with the background (Fig.1). To obtain the
sufficient contrast of diversely pigmented animals with simultaneous
preservation of experimental conditions we are using infrared light
built into the base, on which all IR translucent mazes are placed.
Testing apparatus is uniformly illuminated with the light intensity of
200 lux in the centre of the field. Simultaneously four animals are
tested in four separated open fields for 20 min. Testing arena is
virtually divided to periphery and centre zones, where centre zone
constitutes 38% of whole arena. Presence time, distance travelled, no.
of visits, latency to enter centre zone, average speed, resting and
activity time, velocity classes, and many others indices are
automatically computed for periphery and centre zone as well as for each
5 min. interval.

**Fig. 1:** Tracking the
animals in open fields

**Light-Dark Box**

LDB is divided into two compartments; the light part and enclosed
dark one. The dark part constitute of one third of total apparatus area.
The light part is uniformly illuminated with the light intensity of 430
lux. Each animal is allowed to freely explore the EPM for 10 minutes.
Presence time, distance travelled, no. of visits, latency to enter,
average speed, resting and activity time, velocity classes, and many
others indices are automatically computed for both box’s
compartments.

##### 2.3.2 Neuromotor Abilities

Animal sense of balance and motor coordination can be evaluated in
Beam Walk test and Rotarod, which also allows assessing motor learning
abilities. Other neuromotor functions are assessed by grip strength
measurements and detailed automated gait analysis. Grip Strength
measures maximal muscle strength of forelimbs and hind limbs on an
automated grip strength meter. The test can indicate neuromuscular
abnormalities. Gait Analysis is based on a fully automated analysis of
video records of animal foot prints. Gait analysis provides not only
information about motor coordination but also detailed kinematic
description of animal gait. The test can be used to studding models of
Amyotrophic Lateral Sclerosis (ALS), pain/arthritis, Parkinson’s
disease, muscle injury model or spinal cord injury.

**SHIRPA**

The purpose of SHIRPA assessments is to examine mice for obvious
physical characteristics, behaviours, and morphological
abnormalities.

After one hour of acclimatisation to the experimental room conditions
the mouse is placed into the transparent jar and basic observations of
animal activity and presence are recorded. Subsequently transfer
arousal, animal appearance, basic reflexes, and gait in arena are
estimated. Above the arena the mouse is examined for trunk curl, limb
grasping, and positional passivity. At the end of the evaluation the
animal is placed into the transparent tube. The tube is turned to
position mouse upside down for observing its ability of righting.

**Grip Strength**

Testing is performed in the room with light intensity of 110 lux.
Forelimbs and combined forelimbs and hindlimbs grip-strengths are
determined using an automated meter (Bioseb, Vitrolles, France). To
assess neuromuscular function, the mouse is pulled over a grid; test is
recorded in three trials. To account for potential effects of body mass
all grip strength measurements are normalized to body mass.

##### 2.3.3 Sensorimotor Gating

**Acoustic Startle**

Animal sensorimotor gating is tested in Startle Reflex system, which
allows to determine also Prepulse Inhibition (PPI). The acoustic Startle
Reflex test (SR) is an automated analysis of startle reflex in response
to acoustic stimuli. The test assesses sensorimotor processing by
measuring both afferent sensory information transmission and efferent
motor response (Yeomans and Frankland, 1995). The test can also serve as
a primary screen for hearing impairment. The lack of sufficient sensory
gating mechanism is thought to lead to an overflow of the sensory
stimulation and disintegration of the cognitive functions (Grillon et
al., 1994). SR paradigm is therefore largely used to assess the effects
of putative anti-psychotics and to explore possible genetic and
neurobiological mechanisms of psychosis-related behaviour. PPI is an
attenuation of startle response magnitude by pre-exposure to
non-startling stimulus. PPI provides operational measurement of
sensorimotor gating reflecting the ability of an animal to successfully
integrate relevant and inhibit irrelevant sensory information (Norris
and Blumenthal, 1996). Impaired PPI is observed in schizophrenia as well
as in other neuropsychiatric disorders (Swerdlow and Geyer, 1998).
Testing is automated and performed in three sound proofed cabinets;
protocol execution is controlled by software (Med Associates Inc., USA).
Animals are habituated to the holder for 10 minutes, and then test is
initiated. Four different pre-pulse magnitudes are used for pre-pulse
alone and with startle stimulus. Ten repetitions of startle, pre-pulse
and combination of pre-pulse with startle stimuli are conducted in
pseudorandom order. The amplitude of animal response to all stimuli is
automatically recorded and % of startle response decrease in PPI is
computed manually.

##### 2.3.4 Cognitive functions

Cued and Contextual Conditioning, Context Discrimination, Barnes
Maze, Novel Object Recognition (NOR), and IntelliCage are tests used in
our facility to evaluate animal learning and memory

**Cued and Contextual Conditioning**

Cued and Contextual Conditioning are based on classical Pavlovian
fear conditioning, an associative learning test routinely used to study
biological basis of fear, learning, and memory. Contextual, but not
cued, fear conditioning is regarded as hippocampus-dependent. Although
this statement is true in most of experimental designs, animals can
compensate for hippocampal damage (Frankland et al, 1998). To overcome
this limitation, we also employed Context Discrimination test where
intact hippocampus is critical (Bidenkapp and Rudy, 2007).

Fear Conditioning (FC) is one of the forms of passive learning that
can be used in many strains of mice and rats, even when more pronounced
motor deficits are problematic in other learning assays. The dependent
measure used in contextual and cued FC is a freezing, defined as
“absence of movement except for respiration”, which takes place
following pairing of an unconditioned stimulus (US) with a conditioned
stimulus (CS). Aversive stimulus, foot shock, is used for US, and
neutral cue, tone and a particular context, are used for CS. The
associative learning take place in a cage enclosed within a soundproof
cabinet (Ugo Basile, Gemonio, Italy). The cage is equipped with a
stainless steel rod floor for shock delivery (Fig.2). Testing is
performed in the room with light intensity of 110 lux. The acquisition
trial starts with a 3 min of adaptation period, after which mice are
presented with two pairings of the CS (20 sec of 4 kHz pure tone at 77
dB) and the US (a 1 sec, 0.3 mA constant current to the cage floor). The
US is presented with the termination of the CS. The intra-trial interval
(ITI) is 2 min. Mice remains in the chamber for 1 min after the last
shock delivery. Animals are tested for contextual memory 24 hours after
training. They are reintroduced to the training context and behaviour is
recorded for 3 min with no CS or US presentation. Delayed cue memory is
tested three hours later. Animals are placed in a novel context, with
different cage wall pattern and a smooth floor texture. The freezing
response to the CS is monitored for 2 min. for 2 min.

**Fig. 2:** Freezing
detection of the animal in fear conditioning box

#### 2.4 Metabolism

The metabolism unit uses a multi-technique approach to build a
complex picture of the overall energy storage and expenditure in the
whole animal model. In humans, disturbances in energy balance result in
the development of obesity or anorexia nervosa which presents as weight
gain or weight loss respectively. Cancer cachexia describes a syndrome
of progressive weight loss, anorexia, and persistent erosion of host
body cell mass in response to a malignant growth. A decline in food
intake relative to energy expenditure (which may be increased, normal,
or decreased) is the fundamental physiologic derangement leading to
cancer-associated weight loss. The primary metabolic screen uses
indirect calorimetry, intraperitoneal glucose tolerance test (IPGTT) and
body composition analysis to analyse the overall energy expenditure,
glucose metabolism and energy storage of the model. This data can be
used to guide more in depth analysis at the molecular level to further
understand the complex mechanism pathways controlling and affecting
energy homeostasis.

##### 2.4.1 Indirect Calorimetry

The oxygen consumption, carbon dioxide production and heat production
are measured using the PhenoMaster system from TSE systems. Animals are
individually housed in their home cages (Sealsafe Plus/Green line;
Techniplast) with food and water available ad libitum. Food and water
intake are measured for the duration of the study as well as locomotor
activity (measured by infrared beam cuts). Animals are weighed &
placed in individual cages. The animals are allowed to acclimatise in
the system for 24 hrs. During this time,measurements are recorded,
however the data obtained during this time is not used in the final
analysis (Fig. 1). At the end of the experiment, animals are weighed and
returned to their original home cage.

**Fig. 1:** Calorimetry
workflow

##### 2.4.2 Intraperitoneal glucose tolerance test (IPGTT)

The glucose tolerance test measures the clearance of an
intraperitoneally injected glucose load from the body. It is used to
detect disturbances in glucose metabolism that can be linked to human
conditions such as diabetes or metabolic syndrome.

Animals are transferred to clean cages and fasted overnight
(16-18hrs) with water available ad libitum. Animals are then weighed and
the fasted blood glucose levels is recorded (T=0) from whole blood taken
from the tail vein using a standard glucometer. 20% glucose solution (2g
of glucose/kg body mass) in PBS is by intraperitoneal (IP) injection.
The blood glucose level is measured 15, 30, 60 and 120 mins after the
administration of glucose. After the final measurement, animals are
returned to the original home cage and food and water available ad
libitum.

##### 2.4.3 Body composition

*In vivo* body composition analysis is an important and
effective tool both in early and late adult phenotyping in IMPC to
detect genes involved in energy balance regulation and the development
of lean or obese phenotypes when fed a chow diet. We use an equipment
based on quantitative Nuclear Magnetic Resonance (qNMR), the Minispec
LF90ii (Bruker). This equipment has been specially designed to measure
the body composition of rats and mice, using a magnetic field strength
of 0.47 Teslas. qNMR allows measuring body composition quickly,
non-invasively, and without anesthesia. Implementing qNMR significantly
reduces experimental burdens improving animal welfare. qNMR is a
high-throughput method and suitable for longitudinal studies.

Measurements are performed on each mouse separately. Before the
measurement, we weigh the animal’s body weight. The mice are placed in a
red plastic restrainer or holder, introduced into the magnet tunnel on a
horizontally oriented axis, and the equipment measure the body
composition in less than 4 minutes at room conditions. After the
measurement, the animals are returned to their original cages. With this
procedure, we measure lean mass, fat mass and free fluid mass in a
quick, easy and less stressful way for the animal.

**Fig. 1:** Calorimetry
workflow

#### 2.5 Cardiovascular function

Echocardiography and electrocardiography (ECG) are the most
frequently used techniques for diagnosing cardiovascular diseases. These
methods are non-invasive and the combination of both techniques allows a
first characterization of cardiovascular physiology and function of
genetically modified mice.

##### 2.5.1 Echocardiography

Murine echocardiography is a “gold standard” method to assess
cardiovascular structure and function (systolic and diastolic) in
anaesthetized mice that has been adopted from human system (Ram et al,
2011). In order to visualize the small mouse heart and record the rapid
heart rates, the cardiac ultrasound imaging is acquired using the Vevo
2100 Imaging System (VisualSonics, Inc.) with a 30 MHz transducer
(MS400) operating at a frequency that provides highly reliable and
reproducible image quality. Echocardiography is performed on
anaesthetised (isoflurane) mice and during imaging, the concentration of
anaesthesia is controlled to maintain a heart rate of 450-500 beats/min.
The primary echocardiographic screen includes two-dimensional (2D)
B-mode (“brightness” mode) imaging view of left ventricle (LV) along the
parasternal long axis (PLAX) and M-mode (“motion” mode) image of the
heart in short-axis view (SAX) for accurate linear measurements of left
ventricular internal dimensions (Collins et al., 2009, Moreth et al.,
2014, Stypmann et al, 2009).

**Fig. 1:** Equipment of the
Cardiovascular Phenotyping Unit. Upper panel – Vevo2100 Ultrasound
Imaging System (VisualSonics, Inc.); bottom panel – ECGenie system
(Mouse Specifics, Inc.)

A proper cardiac B-mode view allows visualization of the left atrium
and ventricle, a small portion of the right ventricular wall and the
output of aorta. The apex and the beginning of ascending aorta of the
heart lay on the same horizontal line. M-mode echocardiography is
performed at the ventricular level at the papillary muscle level and
leads to a 1D high-resolution temporal course of the diameter changes of
the LV and the wall thickness of the anterior and posterior LV wall in
systole as well in diastole. All M-mode echocardiography measurements
follow the guidelines of the American Society of Echocardiography (Sahn
et al, 1978). The SAX M-mode images are used to measure left ventricular
end-diastolic internal diameter (LVEDD), left ventricular end-systolic
internal diameter (LVESD), and diastolic and systolic posterior wall
thickness (LVPW) in three consecutive beats. Fractional shortening (FS)
is calculated as FS%=[(LVEDD-LVESD)/LVEDD]x100. Ejection fraction (EF)
is calculated as EF%=100 x ((LVvolD-LVvolS)LVvolD) with
LVvol=((7.0/(2.4+LVID) x LVID3). The corrected left ventricular mass (LV
Mass Cor) is calculated as LV Mass Cor=0.8 (1.053 x ((LVIDD + LVPWD +
IVSD)3 - LVIDD3)). The Stroke volume (SV) is the volume of blood pumped
from one ventricle of the heart with each beat. The Stroke volume of the
LV is obtained by subtracting end-systolic volume (ESV) from
end-diastolic volume (EDV). The cardiac output (CO) is the volume of
blood pumped by the heart per minute. In addition, heart rate and
respiratory rate are calculated by measuring three systolic intervals,
respectively three respiratory intervals.

**Fig. 2:** Representative
B-mode echocardiographic image of an adult mouse heart, PLAX view (a)
and SAX view (b). M-mode image of the left ventricle at level of
papillary muscles (c)

##### 2.5.2 Electrocardiography

Genetically modified mice with a cardiovascular phenotype can be
characterized by recording changes of the rhythm of the heart, these
include slowing (bradycardia) or quickening (tachycardia) of the heart
rate, heart rate variability and/or alterations in the length of the
interval of sinus waves that are reflected in electrocardiogram (ECG)
traces. ECG reflects the electrical activity of the heart and is
comprised of P, Q, R, S, and T waves. The P wave is produced by atrial
depolarization, the QRS pulses represent ventricular depolarization, and
the T wave represents ventricular repolarization (Goldbarg et al.,
1968). Many studies have been conducted to examine the patterns of
activation and repolarization in the mouse heart and their relation to
the electrical cardiac activity (for a review see Boukens et al., 2014).
In our facility, the murine ECG is recorded in conscious mice through
the animal`s paws using the ECGenie system (Mouse Specifics, Inc.). The
size and spacing of disposable footplate electrodes facilitate contact
between the electrodes and the paws to provide Einthoven lead II ECG
(Boukens et al., 2014). For each animal, heart depolarization and
repolarization intervals and amplitudes are evaluated from continuous
ECG recording after 5-min acclimation period. Only runs where at least
15 ECG beats could be included in the analysis are chosen.

Data are analysed using standard protocols for ECG signal analysis by
e-MouseTM (Mouse Specifics, Inc.). The software uses a peak detection
algorithm to find the peak of the R-waves and to calculate heart rate
(HR). The software plots its interpretation of P, Q, R, S, and T for
each beat so that unfiltered noise or motion artefacts are rejected.
This is followed by calculations of the mean of the ECG time intervals
for each set of waveforms. The corrected QT interval (QTc) is calculated
by dividing the QT interval by the square root of the preceding RR
interval. QT dispersion is measured as inter-lead variability of QT
intervals. The QTc dispersion was calculated as the rate corrected QT
dispersion.

**Fig. 3:**
Schematic representation of the murine ECG

#### 2.6 Lung function

For an optimal gas-exchange in the lungs the air should be able to
easily flow in and out of the lungs. Key parameters for this are the
resistance the air encounters and the elasticity of the lung tissue
which allows the lungs to empty themselves efficiently during the
exhalation phase.

Both these parameters can be measured by the ‘Forced Oscillation’
technique in which pressure waves, called perturbations, are applied to
the lungs and the response to these pressure waves is recorded and
analyzed. To perform these perturbation mice are first anaesthetized
with ketamine and xylazine and intubated after which they are attached
to a computer controlled piston ventilator, the Flexivent FX from
Scireq. Mice are ventilated at 190 breaths per minute, which allows for
an optimal oxygenation and fast recovery of the oxygen saturation after
perturbations.

For the analysis of the response curves of the perturbations, a
mathematical model of the lungs is used. There are mainly 2 models and
also 2 types of perturbation.

In the first model, called the single compartment model, the lung is
modelled as a balloon, which confers the elasticity and represents the
lung tissue, attached to a tube, which confers the resistance to the
airflow and represents the airways. This model is simple but very robust
and the perturbation applied is relatively simple, it just consists of a
sinusoidal wave at a frequency of 150 cycles per minute. This
perturbation is called the SnapShot-150, which is abbreviated SN in
figures. From this 2 variables are calculated, the airway resistance (R)
and the elastance (E) of the lungs. Sometimes the elasticity parameter
is expressed as compliance (C), which is the reciprocal of the elastance
(C = 1/E).

The second model, the complex phase model, is more complex and it
models the airway resistance (R) and the resistance in the lung tissue,
called tissue damping (G), separately. This is possible due to a
difference in frequency response of these 2 structures. For the
elasticity parameters, the component of the airways is negligible and
thus only 1 elasticity parameter is calculated, the tissue elastance
(H). As the difference between airways and lung-tissue is the frequency
response of these tissues, the perturbation is a multi-frequency
perturbation. We use 2 different ones in our primary screen. The first
one takes 3 seconds, which gives a less precise estimate, but which can
be performed in mice with severely compromised lung-functions as the
period without fresh air is only 3 seconds. This manoeuver is called
Quick-prime 3, abbreviated as QP3 in figures. The second manoeuver, the
Prime-8, abbreviated P8, last as its name suggests, 8 seconds. For a
mouse breathing normally at about 180 to 200 breaths per minute this is
relatively long and could thus lead to oxygen desaturation in animals
with severely compromised lung function. In this case the animal will
start spontaneous breathing movements which will compromise the
measurement. This estimation of the lung function parameters is more
accurate as more data points are available for model fitting.

There is an additional manoeuver which is performed which is called
‘Deep inflation’ and is a forced deep inhalation. This serves 2
purposes, firstly, to measure the lung capacity of the animal and
secondly to open up the lungs completely so that all parts of the lung
are recruited in the consecutive manoeuvers.

All manoeuvers are executed in an automated script 5 times. If some
measurements are invalid during script execution, they are repeated
until 5 valid measures are obtained, with the exception of the prime-8
measurements. These measurements are not reattempted after the script if
obvious oxygen desaturation occurs, as it would be harmful for the
animal and would not lead to good quality data.

#### 2.7 Bioimaging

The bioimaging unit is dedicated to comprehensive morphological and
functional characterization of animal models by whole-body imaging
systems in vivo and ex vivo. The primary tests cover morphological
analysis of the entire skeleton and quantitative analysis of mineral
content. Data for 3D body composition analysis is also acquired during
the same scan (Judex et al., 2010). Secondary tests and supplemental
approaches are established for more detailed and comprehensive
characterization of skeleton related phenotypes:

- detail ex-vivo bone structure imaging with quantification of bone
  porosity and mineral content (osteoporosis or arthritis models), or
  in-vivo of selected skeleton area (e.g. femur or autopodium)
- segmentation and volumetric analysis of individual bones – for
  craniofacial phenotypes)
- enamel and dentin mineralization imaging (Wald et al., 2017)
- customized optical imaging including fluorescence protein detection,
  fluorescence probes for inflammation, tumour development or bone
  remodeling
- skeletal preparation and alizarin/alcian staining for bone/cartilage
  visualization.

Animals are anaesthetised using 20% zoletil solution. Once
anaesthetized, animals are weighed and body length is recorded with dial
caliper. Animals are placed into the animal restrainer and the scan is
initiated using the SkyScan software.

**microCT (Skyscan 1176, Bruker)** *in vivo*
scanner provides sensitive and high resolution 3D imaging based on X-ray
projections with voxel size 9 – 35μm. Our imaging set up is suitable
either for *in vivo* or *ex vivo* imaging of large
specimen up to size of rat. *In vivo* scans can provide
comprehensive 3D visualization of bones and other mineralized tissues
like teeth, but also quantification of total body fat mass and lean mass
(alternative to DEXA analysis) and their changes in time.

*Ex vivo* scanning mode can be used for higher resolution
imaging of large body parts or use of contrast agents for imaging soft
tissues (liver, kidney, heart, neuronal tissue) or imaging fixed embryos
(E9.5 – E18.5) with use of appropriate contrast agents (Iodine, PTA).
For complete microCT analysis post-processing of projection data is
necessary, especially 3D reconstruction and further segmentation for
visualization of morphological phenotypes and quantification of selected
organs.

The system calibrates automatically after turning on after 3 or more
days without usage, plus flat field correction is processed whenever the
scanning parameters are changed.

**Optical whole body imaging system (Xtreme, Bruker)**
includes highly sensitive camera for detection of emitted photons based
on enzymatic activity or fluorescence. This approach is suitable for in
vivo imaging with emphasis on noninvasive imaging setup for longitudinal
monitoring of studied process. This imaging modality is established for
inflammation, tumour progression, metastasis or cell homing experiments.
We have successfully published our imaging protocol in the study of
genetic regulation of DSS colitis (Brauer et al., 2015).

**Software tool box:** NRecon, DataViewer, ITK-snap,
3D-slicer, CTvox, CTanalyser, CTvolume,Imaris, microCT (Skyscan 1176,
Bruker), Optical whole body imaging system (Xtreme, Bruker)

#### 2.8 Auditory Brain Stem Response (Hearing)

Auditory Brainstem Response (ABR) is an acoustically evoked
electrical potential recorded by needle electrodes placed subdermally on
the top of animal’s head. It represents a sum of activity of brainstem
neurons responsible for sound processing. Typical ABR evoked by a short
sound of supra-threshold intensity is a waveform composed of several
successive peaks that appear between 2 – 6 ms after the sound onset.
Their amplitudes vary in the range of hundreds of nanovolts and
correlate with the stimulus intensity. The latencies of individual ABR
peaks reflect successive activation of the auditory nerve, the cochlear
nucleus, the superior olivary complex and the inferior colliculus, which
comprise the brainstem portion of the auditory pathway (for review see
Biacabe et al., 2001; Møller, 2006).

The ABR recording protocol follows guidelines as described by Ingham
et al., 2011. The mouse is anaesthetized by i.p. injection of Zoletil
(mixture of tiletamine and zolazepam) and placed on a heating pad inside
an 6 m^3^ anechoic soundproof chamber. Three subdermal needle
electrodes are positioned as follows: active electrode on the vertex,
reference electrode behind the left ear and ground electrode behind the
right ear, see Fig. 1. The system for sound production and biosignal
recording is based on TDT workstation RZ6 (Tucker-Davies Technologies,
USA). Acoustic stimuli are delivered by a high quality ribbon tweeter
RAAL 140-15D (RAAL Advanced Loudspeakers, Serbia) placed approximately
40 cm in front of the mouse. Each stimulus is played repeatedly and the
averaged ABR signal is recorded. Initially, the ABRs are evoked by
broadband stimuli (“clicks”, i.e. 100 microseconds lasting rectangular
pulses) followed by bursts of pure tones (5 ms long, 1 ms rise/fall
times) of 306, 2412, 1818, 12 24 and 630 kHz frequency respectively.
Stimulation always starts with the lowest sound intensity of 0 dB SPL
(sound pressure level) then the intensity is increased by 5 dB steps
until 85 dB SPL is reached.

**Fig. 1:** mouse
under general anesthesia with subcutaneous electrodes for ABR recording.
Black wire - reference electrode, behind the left ear, green wire -
ground electrode behind the right ear, red wire - active electrode on
the vertex.

The primary output of ABR analysis is detection of hearing thresholds
for the broadband stimulus (click) and for each frequency used. The
threshold is defined as the lowest intensity of sound which evoked
distinguishable response related to (time-locked to) the stimulus.
Statistical significance of the difference in thresholds between two
groups of animals is tested by two-way ANOVA with replication combined
with post-hoc tests. Other parameters such as latency, amplitude and
waveform shape may be examined in addition if it is of interest.

**Fig. 2:**
Auditory Brainstem Response. The animal was stimulated by a pure tone of
24 kHz frequency, 5 ms duration and intensities ranging between 0 - 85
dB SPL, the stimulus onset is marked with a vertical arrow (Stim.). Each
curve represents an average of 600 repetitions. The lowest sound
intensity that evoked a response is referred to as the hearing threshold
for the respective frequency, marked with a horizontal arrow (Thr.).
Five typical peaks of ABR signal are labelled with Roman numerals I to
V. Note that with increasing intensity not only the peaks are higher but
they have shorter latency, too.

#### 2.9 Eye morphology

Eyes are a very important sense organ for human beings.
Unfortunately, many people suffer from serious ocular diseases like
cataracts, glaucoma or retinal disorders and their life quality is
strongly affected. Many genes are involved in a variety of ocular
disorders. As the genome of the mouse is well known, the mouse is a
suitable model for understanding of the biochemical, genetic and
physiological mechanisms of many hereditary degenerative ocular
disorders.

The vision screen uses non-invasive imaging devices to examine the
anterior segment (Pentacam) and retina of the eyes (OCT). The animals
are anaesthetised using 20% Zoletil and the pupils are dilated using eye
drops containing 0.5% Atropin. The mouse is placed on a platform of the
Pentacam and the centre of the pupil is orientated to the light-beam
slit of the Pentacam´s head. This device enables the measurement of many
parameters of the cornea and the lens (e.g. surface, form, opacity,
thickness and density) for each eye. All 25 scanned images from
different angles of the eye are carried out by a rotating Scheimpflug
Camera of the Pentacam they are analysed in order to assess any
abnormalities of the eye.

The optical coherent tomograph (OCT) scans, quantifies the reflection
of a light beam sending from the layers of the retina and composes
cross-sectional images of the retina. Each cross-section image of the
retina will be evaluated and a variety of parameters are measured
e.g. the thickness and the gross morphology of the retina, form and
position of the optical disc, the blood vessels and their pattern.

**Fig. 1:** Representative
example of the retinal fundus with the optic disc and blood vessel
system (left) and a cross-sectional image of retina (right) in the wild
type mouse C57Bl/6NCrl.

#### 2.10 Biochemistry and Hematology

Biochemical and haematology phenotyping is based on robust primary
screening developed under Eumorphia and EMPReSS, a European Mouse
Phenotyping Resource for Standardised Screens. Phenotyping investigation
includes also newest development in INFRAFRONTIER and IMPC consortia.
These tests comprise primary phenotyping pipeline and are carried out
according the standardised protocols as detailed in (IMPReSS). The
Biochemical and haematology unit currently analyses various blood
metabolites, ions, hormones, and enzymes of genetically modified mice
compared with age matched controls. Changes in these data can be linked
to metabolic and functional abnormalities of specific organs such as
liver, kidney, and gastrointestinal tract.

**Blood Withdrawal and Storage**

Blood samples are taken from isoflurane-anesthetized mice by
retro-bulbar sinus puncture with non-heparinized glass capillaries.
Samples are collected in two different types of coated tubes
(Lithium/Heparin (KABE cat # 078028) and EDTA (KABE cat # 078035). After
collection each sample is mixed by gentle inversion and then kept on RT
until centrifugation. Samples are centrifuged within 1 hour of
collection at 5000 X g, for 10 minutes at 8°C. Once separated from the
cells, plasma samples are refrigerated if analysis is delayed. If plasma
cannot be analyzed fresh, samples are stored at -20oC (short-term) or
-70oC (long-term) storage. Freeze/thaw cycles are avoided.

Plasma is analyzed on Beckman AU480 biochemical analyzer. All
calibration and quality control samples are within expiration period and
measured values fulfilled required parameters for each method used in
this report.

**Hematology**

Our hematology test requires 20 μl of whole blood in EDTA coated
tubes per sample. 20 μl of whole blood is diluted to final volume is 200
μl. Complete and differential blood cell counts were measured on
Mindray-BC-5000Vet

#### 2.11 Immunology

As a integral part of the terminal screen, immunophenotyping involves
characterisation of particular immune cell populations in terms of their
cellularity and phenotype using multicolor flow cytometry.

The procedures are based on standard immunophenotyping protocols of
the Adult and Embryonic Phenotype Pipeline that has been agreed by the
research institutions involved MPReSS
-International Mouse Phenotyping Resource of Standardised
Screens.

According to these guidelines, we use mouse spleenocytes for standard
immunophenotyping. However, we are able to analyze peripheral blood,
lymph nodes, thymi, bone marrow or peritoneal lavage samples as
well.

**Fig. 1:** Gating strategy
to discriminate T cell populations using Panel A

Routinelly, we utilize two panels to detect lymphoid and myeloid
cells. In addition, the spleen weight is an important indicator which is
also recorded. The procedure includes several quality control steps
(cell viability upon digestion, number of cells/sample after
acquisition).

For primary tests, we typically use three males and three females
knock out (KO) animals plus three male and three female wild type
controls (WT), plus optionally one or two control mice as internal
controls. Flow cytometry data are acquired using BD LSRFortessaTM SORP,
manualy analyzed in FlowJo software and statistically evaluated.

**Fig. 2:** Gating strategy
to discriminate B cell and myeloid populations using Panel B

#### 2.12 Histopathology

The histopathology screen is dedicated for macroscopic and
microscopic analysis of pathological alterations occurring in mutant
mouse models. The unit is comparing gross morphology and microscopic
differences between wild-type and gene-engineered mouse models. A major
task is to screen for set of pathologies connected to specific genetic
status using the tools and techniques of the classical and molecular
pathology.

##### 2.12.1 Gross Morphology and Tissue Processing

**Full mouse necropsy with organ isolation**

A complete necropsy is performed to detect and record abnormal
macroscopic alterations in internal and external organs, record body,
spleen, liver, kidney (dx., sin.) and heart weights (according to IMPC
Heart Weight SOP) and tibia length.

Samples from IMPC ‘Tissue Collection List’ (minimal number is 2
mutant males and 2 mutant females) are fixed, trimmed, processed and
embedded in paraffin blocks. The femur is also prepared and blood smears
and bone marrow smears are also prepared.

Images of all abnormal findings are documented and saved in database.
All collected tissues are placed in a labelled jar containing a
sufficient volume of fixative (minimum of 10:1 fixative to tissue). All
abnormal findings are recorded according to our Centre-specific database
using the standardized IMPC Gross Pathology ontology. For long-term
storage, the fixed samples are stored in paraffin blocks.

**Organ sampling and trimming**

Individual organs are processed. Unless otherwise specified, organs
are processed according to the *Revised guides for organ sampling and
trimming in rats and mice*, published in 2003 and 2004 by the
Registry of Industrial Toxicology Animal-data (RITA) and the North
American Control Animal Database (NACAD) groups (Exp Toxic Pathol 55:
91-106, Exp Tox Pathol 55: 413-431, and Exp Tox Pathol 55: 433-449).

**Tissue processing and embedding**

By using a state-of-the-art automated vacuum tissue processor (Leica
ASP6025), we process and paraffin-embed specimens with the highest
levels of reproducibility and quality. Where applicable, decalcification
to remove mineral from bone or other calcified tissues is performed
prior to processing to paraffin. Tissues are embedded in paraffin using
standard orientation according to IMPC Tissue Embedding and Block
Banking IMPC\_BLK\_001 SOP.

##### 2.12.2 Sectioning and Staining

**Sectioning**

Standard paraffin sections are cut using a Leica RM2255-FU microtome.
The standard thickness of sections is 2µm.

**H&E staining**

The standard primary staining procedure is the hematoxylin and eosin
(H&E) stain. The use of the Ventana Symphony automated staining
system ensures high throughput processing with consistent and
reproducible results.

##### 2.12.3 Histopathological examination

Histopathological examination is performed using the Carl Zeiss Axio
Imager.Z2 or Leica DM3000 microscope. All abnormal findings are recorded
in our Centre-specific database using the standardized IMPC
Histopatology ontology.

### 3 Results

#### 3.1 Weight Curve - 4 week to 16 week

| STRAIN | GENDER | count |
| --- | --- | --- |
| WT/WT | female | 8 |
| WT/WT | male | 8 |
| Δ HEL/WT | female | 8 |
| Δ HEL/WT | male | 8 |

|  | test | Difference | p\_value | Significance |
| --- | --- | --- | --- | --- |
| c2 | WT/WT vs Δ HEL/WT | 2.9151 | 0.0000 | TRUE |
| c3 | Female vs Male | -5.6632 | 0.0000 | TRUE |
| c4 | Female WT/WT vs Female Δ HEL/WT | 1.8323 | 0.1636 | FALSE |
| c5 | Male WT/WT vs Male Δ HEL/WT | 3.9979 | 0.0000 | TRUE |
| c6 | Female Δ HEL/WT vs Male Δ HEL/WT | -4.5804 | 0.0000 | TRUE |

#### 3.2 Open field

**Number of animals**

| STRAIN | GENDER | count |
| --- | --- | --- |
| WT/WT | female | 8 |
| WT/WT | male | 8 |
| Δ HEL/WT | female | 8 |
| Δ HEL/WT | male | 8 |

##### 3.2.1 Center average speed - total (cm/s)

|  | test | Difference | p\_value | Significance |
| --- | --- | --- | --- | --- |
| c2 | WT/WT vs Δ HEL/WT | 0.550 | 0.9415 | FALSE |
| c3 | Female vs Male | -1.175 | 0.7910 | FALSE |
| c4 | Female WT/WT vs Female Δ HEL/WT | -0.725 | 0.9519 | FALSE |
| c5 | Male WT/WT vs Male Δ HEL/WT | 1.825 | 0.5477 | FALSE |
| c6 | Female Δ HEL/WT vs Male Δ HEL/WT | 0.100 | 0.9999 | FALSE |

##### 3.2.2 Center distance travelled - total (cm)

|  | test | Difference | p\_value | Significance |
| --- | --- | --- | --- | --- |
| c2 | WT/WT vs Δ HEL/WT | -185.0625 | 0.8086 | FALSE |
| c3 | Female vs Male | 48.8875 | 0.9955 | FALSE |
| c4 | Female WT/WT vs Female Δ HEL/WT | -557.5625 | 0.2371 | FALSE |
| c5 | Male WT/WT vs Male Δ HEL/WT | 187.4375 | 0.9188 | FALSE |
| c6 | Female Δ HEL/WT vs Male Δ HEL/WT | 421.3875 | 0.4867 | FALSE |

##### 3.2.3 Center permanence time - total (s)

|  | test | Difference | p\_value | Significance |
| --- | --- | --- | --- | --- |
| c2 | WT/WT vs Δ HEL/WT | 4.2012 | 0.9997 | FALSE |
| c3 | Female vs Male | 35.1363 | 0.9049 | FALSE |
| c4 | Female WT/WT vs Female Δ HEL/WT | -6.0500 | 0.9997 | FALSE |
| c5 | Male WT/WT vs Male Δ HEL/WT | 14.4524 | 0.9959 | FALSE |
| c6 | Female Δ HEL/WT vs Male Δ HEL/WT | 45.3875 | 0.9116 | FALSE |

##### 3.2.4 Center resting time - total (s)

|  | test | Difference | p\_value | Significance |
| --- | --- | --- | --- | --- |
| c2 | WT/WT vs Δ HEL/WT | 10.7177 | 0.9922 | FALSE |
| c3 | Female vs Male | 34.0483 | 0.8754 | FALSE |
| c4 | Female WT/WT vs Female Δ HEL/WT | 16.7891 | 0.9894 | FALSE |
| c5 | Male WT/WT vs Male Δ HEL/WT | 4.6463 | 0.9998 | FALSE |
| c6 | Female Δ HEL/WT vs Male Δ HEL/WT | 27.9769 | 0.9648 | FALSE |

##### 3.2.5 Distance travelled - total (cm)

|  | test | Difference | p\_value | Significance |
| --- | --- | --- | --- | --- |
| c2 | WT/WT vs Δ HEL/WT | -88.2938 | 0.9973 | FALSE |
| c3 | Female vs Male | 73.5188 | 0.9985 | FALSE |
| c4 | Female WT/WT vs Female Δ HEL/WT | -981.8875 | 0.4175 | FALSE |
| c5 | Male WT/WT vs Male Δ HEL/WT | 805.3000 | 0.5873 | FALSE |
| c6 | Female Δ HEL/WT vs Male Δ HEL/WT | 967.1125 | 0.4336 | FALSE |

##### 3.2.6 Number of center entries - total

|  | test | Difference | p\_value | Significance |
| --- | --- | --- | --- | --- |
| c2 | WT/WT vs Δ HEL/WT | -13.5625 | 0.2289 | FALSE |
| c3 | Female vs Male | -0.1875 | 1.0000 | FALSE |
| c4 | Female WT/WT vs Female Δ HEL/WT | -28.5000 | 0.0274 | TRUE |
| c5 | Male WT/WT vs Male Δ HEL/WT | 1.3750 | 0.9991 | FALSE |
| c6 | Female Δ HEL/WT vs Male Δ HEL/WT | 14.7500 | 0.4569 | FALSE |

##### 3.2.7 Periphery average speed - total (cm/s)

|  | test | Difference | p\_value | Significance |
| --- | --- | --- | --- | --- |
| c2 | WT/WT vs Δ HEL/WT | 0.1938 | 0.9064 | FALSE |
| c3 | Female vs Male | 0.0812 | 0.9924 | FALSE |
| c4 | Female WT/WT vs Female Δ HEL/WT | -0.2875 | 0.8937 | FALSE |
| c5 | Male WT/WT vs Male Δ HEL/WT | 0.6750 | 0.3527 | FALSE |
| c6 | Female Δ HEL/WT vs Male Δ HEL/WT | 0.5625 | 0.5219 | FALSE |

##### 3.2.8 Periphery distance travelled - total (cm)

|  | test | Difference | p\_value | Significance |
| --- | --- | --- | --- | --- |
| c2 | WT/WT vs Δ HEL/WT | 159.2750 | 0.9553 | FALSE |
| c3 | Female vs Male | -37.8875 | 0.9993 | FALSE |
| c4 | Female WT/WT vs Female Δ HEL/WT | -299.3250 | 0.9029 | FALSE |
| c5 | Male WT/WT vs Male Δ HEL/WT | 617.8750 | 0.4972 | FALSE |
| c6 | Female Δ HEL/WT vs Male Δ HEL/WT | 420.7125 | 0.7729 | FALSE |

##### 3.2.9 Periphery permanence time - total (s)

|  | test | Difference | p\_value | Significance |
| --- | --- | --- | --- | --- |
| c2 | WT/WT vs Δ HEL/WT | -4.1687 | 0.9997 | FALSE |
| c3 | Female vs Male | -35.2062 | 0.9045 | FALSE |
| c4 | Female WT/WT vs Female Δ HEL/WT | 6.0375 | 0.9997 | FALSE |
| c5 | Male WT/WT vs Male Δ HEL/WT | -14.3750 | 0.9960 | FALSE |
| c6 | Female Δ HEL/WT vs Male Δ HEL/WT | -45.4125 | 0.9116 | FALSE |

##### 3.2.10 Periphery resting time - total (s)

|  | test | Difference | p\_value | Significance |
| --- | --- | --- | --- | --- |
| c2 | WT/WT vs Δ HEL/WT | -5.6434 | 0.9991 | FALSE |
| c3 | Female vs Male | -31.1632 | 0.9145 | FALSE |
| c4 | Female WT/WT vs Female Δ HEL/WT | 22.5107 | 0.9817 | FALSE |
| c5 | Male WT/WT vs Male Δ HEL/WT | -33.7975 | 0.9421 | FALSE |
| c6 | Female Δ HEL/WT vs Male Δ HEL/WT | -59.3173 | 0.7895 | FALSE |

##### 3.2.11 Whole arena average speed - total (cm/s)

|  | test | Difference | p\_value | Significance |
| --- | --- | --- | --- | --- |
| c2 | WT/WT vs Δ HEL/WT | -0.0938 | 0.9945 | FALSE |
| c3 | Female vs Male | 0.0562 | 0.9989 | FALSE |
| c4 | Female WT/WT vs Female Δ HEL/WT | -0.8250 | 0.4159 | FALSE |
| c5 | Male WT/WT vs Male Δ HEL/WT | 0.6375 | 0.6319 | FALSE |
| c6 | Female Δ HEL/WT vs Male Δ HEL/WT | 0.7875 | 0.4643 | FALSE |

##### 3.2.12 Whole arena resting time - total (s)

|  | test | Difference | p\_value | Significance |
| --- | --- | --- | --- | --- |
| c2 | WT/WT vs Δ HEL/WT | 4.50 | 0.9937 | FALSE |
| c3 | Female vs Male | 3.15 | 0.9979 | FALSE |
| c4 | Female WT/WT vs Female Δ HEL/WT | 37.95 | 0.4112 | FALSE |
| c5 | Male WT/WT vs Male Δ HEL/WT | -28.95 | 0.6374 | FALSE |
| c6 | Female Δ HEL/WT vs Male Δ HEL/WT | -30.30 | 0.6059 | FALSE |

##### 3.2.13 Center average speed

|  | test | Difference | p\_value | Significance |
| --- | --- | --- | --- | --- |
| c2 | WT/WT vs Δ HEL/WT | 1.1740 | 0.5632 | FALSE |
| c3 | Female vs Male | -0.8568 | 0.9938 | FALSE |
| c4 | Female WT/WT vs Female Δ HEL/WT | -0.1420 | 0.9995 | FALSE |
| c5 | Male WT/WT vs Male Δ HEL/WT | 2.4900 | 0.2204 | FALSE |
| c6 | Female Δ HEL/WT vs Male Δ HEL/WT | 0.4591 | 0.9991 | FALSE |

##### 3.2.14 Center distance travelled

|  | test | Difference | p\_value | Significance |
| --- | --- | --- | --- | --- |
| c2 | WT/WT vs Δ HEL/WT | -29.8241 | 0.9348 | FALSE |
| c3 | Female vs Male | 23.3947 | 0.9982 | FALSE |
| c4 | Female WT/WT vs Female Δ HEL/WT | -128.7124 | 0.2832 | FALSE |
| c5 | Male WT/WT vs Male Δ HEL/WT | 69.0643 | 0.7694 | FALSE |
| c6 | Female Δ HEL/WT vs Male Δ HEL/WT | 122.2830 | 0.8353 | FALSE |

##### 3.2.15 Center permanence time

|  | test | Difference | p\_value | Significance |
| --- | --- | --- | --- | --- |
| c2 | WT/WT vs Δ HEL/WT | 0.8989 | 0.9997 | FALSE |
| c3 | Female vs Male | 12.6685 | 0.9830 | FALSE |
| c4 | Female WT/WT vs Female Δ HEL/WT | -8.5481 | 0.9188 | FALSE |
| c5 | Male WT/WT vs Male Δ HEL/WT | 10.3459 | 0.8660 | FALSE |
| c6 | Female Δ HEL/WT vs Male Δ HEL/WT | 22.1155 | 0.9262 | FALSE |

##### 3.2.16 Center resting time

|  | test | Difference | p\_value | Significance |
| --- | --- | --- | --- | --- |
| c2 | WT/WT vs Δ HEL/WT | -1.1290 | 0.9985 | FALSE |
| c3 | Female vs Male | 11.0548 | 0.9983 | FALSE |
| c4 | Female WT/WT vs Female Δ HEL/WT | -7.2540 | 0.8842 | FALSE |
| c5 | Male WT/WT vs Male Δ HEL/WT | 4.9959 | 0.9579 | FALSE |
| c6 | Female Δ HEL/WT vs Male Δ HEL/WT | 17.1798 | 0.9938 | FALSE |

##### 3.2.17 Distance travelled

|  | test | Difference | p\_value | Significance |
| --- | --- | --- | --- | --- |
| c2 | WT/WT vs Δ HEL/WT | 10.8654 | 0.9996 | FALSE |
| c3 | Female vs Male | 9.6069 | 1.0000 | FALSE |
| c4 | Female WT/WT vs Female Δ HEL/WT | -203.7555 | 0.5502 | FALSE |
| c5 | Male WT/WT vs Male Δ HEL/WT | 225.4862 | 0.4631 | FALSE |
| c6 | Female Δ HEL/WT vs Male Δ HEL/WT | 224.2278 | 0.9339 | FALSE |

##### 3.2.18 Number of center entries

|  | test | Difference | p\_value | Significance |
| --- | --- | --- | --- | --- |
| c2 | WT/WT vs Δ HEL/WT | -2.6436 | 0.4218 | FALSE |
| c3 | Female vs Male | 0.3762 | 0.9999 | FALSE |
| c4 | Female WT/WT vs Female Δ HEL/WT | -6.3519 | 0.0525 | FALSE |
| c5 | Male WT/WT vs Male Δ HEL/WT | 1.0646 | 0.9717 | FALSE |
| c6 | Female Δ HEL/WT vs Male Δ HEL/WT | 4.0845 | 0.9064 | FALSE |

##### 3.2.19 Periphery average speed

|  | test | Difference | p\_value | Significance |
| --- | --- | --- | --- | --- |
| c2 | WT/WT vs Δ HEL/WT | 0.2183 | 0.8657 | FALSE |
| c3 | Female vs Male | 0.0799 | 0.9999 | FALSE |
| c4 | Female WT/WT vs Female Δ HEL/WT | -0.2622 | 0.9125 | FALSE |
| c5 | Male WT/WT vs Male Δ HEL/WT | 0.6988 | 0.3049 | FALSE |
| c6 | Female Δ HEL/WT vs Male Δ HEL/WT | 0.5604 | 0.9597 | FALSE |

##### 3.2.20 Periphery distance travelled

|  | test | Difference | p\_value | Significance |
| --- | --- | --- | --- | --- |
| c2 | WT/WT vs Δ HEL/WT | 44.2225 | 0.9327 | FALSE |
| c3 | Female vs Male | -16.6322 | 0.9999 | FALSE |
| c4 | Female WT/WT vs Female Δ HEL/WT | -73.5711 | 0.8959 | FALSE |
| c5 | Male WT/WT vs Male Δ HEL/WT | 162.0160 | 0.4172 | FALSE |
| c6 | Female Δ HEL/WT vs Male Δ HEL/WT | 101.1614 | 0.9773 | FALSE |

##### 3.2.21 Periphery permanence time

|  | test | Difference | p\_value | Significance |
| --- | --- | --- | --- | --- |
| c2 | WT/WT vs Δ HEL/WT | -0.9067 | 0.9997 | FALSE |
| c3 | Female vs Male | -12.6778 | 0.7858 | FALSE |
| c4 | Female WT/WT vs Female Δ HEL/WT | 8.5407 | 0.9216 | FALSE |
| c5 | Male WT/WT vs Male Δ HEL/WT | -10.3541 | 0.8697 | FALSE |
| c6 | Female Δ HEL/WT vs Male Δ HEL/WT | -22.1253 | 0.5451 | FALSE |

##### 3.2.22 Periphery resting time

|  | test | Difference | p\_value | Significance |
| --- | --- | --- | --- | --- |
| c2 | WT/WT vs Δ HEL/WT | -2.1205 | 0.9965 | FALSE |
| c3 | Female vs Male | -9.6907 | 0.9004 | FALSE |
| c4 | Female WT/WT vs Female Δ HEL/WT | 9.5286 | 0.9053 | FALSE |
| c5 | Male WT/WT vs Male Δ HEL/WT | -13.7696 | 0.7620 | FALSE |
| c6 | Female Δ HEL/WT vs Male Δ HEL/WT | -21.3397 | 0.6040 | FALSE |

##### 3.2.23 Whole arena average speed

|  | test | Difference | p\_value | Significance |
| --- | --- | --- | --- | --- |
| c2 | WT/WT vs Δ HEL/WT | 0.0275 | 0.9998 | FALSE |
| c3 | Female vs Male | 0.0210 | 1.0000 | FALSE |
| c4 | Female WT/WT vs Female Δ HEL/WT | -0.6865 | 0.5463 | FALSE |
| c5 | Male WT/WT vs Male Δ HEL/WT | 0.7416 | 0.4803 | FALSE |
| c6 | Female Δ HEL/WT vs Male Δ HEL/WT | 0.7351 | 0.7923 | FALSE |

##### 3.2.24 Whole arena resting time

|  | test | Difference | p\_value | Significance |
| --- | --- | --- | --- | --- |
| c2 | WT/WT vs Δ HEL/WT | 1.6251 | 0.9814 | FALSE |
| c3 | Female vs Male | 0.8029 | 0.9999 | FALSE |
| c4 | Female WT/WT vs Female Δ HEL/WT | 9.9993 | 0.3619 | FALSE |
| c5 | Male WT/WT vs Male Δ HEL/WT | -6.7491 | 0.6849 | FALSE |
| c6 | Female Δ HEL/WT vs Male Δ HEL/WT | -7.5713 | 0.9499 | FALSE |

##### 3.2.25 Latency to center entry (s)

|  | test | Difference | p\_value | Significance |
| --- | --- | --- | --- | --- |
| c2 | WT/WT vs Δ HEL/WT | 32.4563 | 0.1763 | FALSE |
| c3 | Female vs Male | 17.7312 | 0.8025 | FALSE |
| c4 | Female WT/WT vs Female Δ HEL/WT | 52.5375 | 0.0944 | FALSE |
| c5 | Male WT/WT vs Male Δ HEL/WT | 12.3750 | 0.9446 | FALSE |
| c6 | Female Δ HEL/WT vs Male Δ HEL/WT | -2.3500 | 0.9997 | FALSE |

##### 3.2.26 Percentage center time - total (%)

|  | test | Difference | p\_value | Significance |
| --- | --- | --- | --- | --- |
| c2 | WT/WT vs Δ HEL/WT | 0.3458 | 0.9997 | FALSE |
| c3 | Female vs Male | 2.9333 | 0.8676 | FALSE |
| c4 | Female WT/WT vs Female Δ HEL/WT | -0.5031 | 0.9997 | FALSE |
| c5 | Male WT/WT vs Male Δ HEL/WT | 1.1948 | 0.9960 | FALSE |
| c6 | Female Δ HEL/WT vs Male Δ HEL/WT | 3.7823 | 0.8945 | FALSE |

#### 3.3 Shirpa and Dismorphology

**Number of animals**

| STRAIN | GENDER | count |
| --- | --- | --- |
| WT/WT | female | 8 |
| WT/WT | male | 8 |
| Δ HEL/WT | female | 8 |
| Δ HEL/WT | male | 8 |

##### 3.3.1 Activity (body position)

##### 3.3.2 Aggression

##### 3.3.3 Coat - color - abdomen

##### 3.3.4 Coat - color - back

##### 3.3.5 Coat - color - head

##### 3.3.6 Coat - color pattern - abdomen

##### 3.3.7 Coat - color pattern - back

##### 3.3.8 Coat - color pattern - head

##### 3.3.9 Coat - hair distribution - abdomen

##### 3.3.10 Coat - hair distribution - back

##### 3.3.11 Coat - hair distribution - head

##### 3.3.12 Coat - hair texture\_appearance - abdomen

##### 3.3.13 Coat - hair texture\_appearance - back

##### 3.3.14 Coat - hair texture\_appearance - head

##### 3.3.15 Contact righting

##### 3.3.16 Ears

##### 3.3.17 Forelimb digit - number

##### 3.3.18 Forelimb digit - shape

##### 3.3.19 Forelimb digit - size

##### 3.3.20 Forelimb nail - length

##### 3.3.21 Forelimb nail - number

##### 3.3.22 Forelimb nail - shape

##### 3.3.23 Forelimbs - position

##### 3.3.24 Forelimbs - shape

##### 3.3.25 Forelimbs - size

##### 3.3.26 Forepaw - shape

##### 3.3.27 Forepaw - size

##### 3.3.28 Gait

##### 3.3.29 Genitalia - morphology

##### 3.3.30 Genitalia - presence

##### 3.3.31 Genitalia - size

##### 3.3.32 Head bobbing

##### 3.3.33 Head morphology

##### 3.3.34 Head size

##### 3.3.35 Hindlimb digit - number

##### 3.3.36 Hindlimb digit - shape

##### 3.3.37 Hindlimb digit - size

##### 3.3.38 Hindlimb nail - length

##### 3.3.39 Hindlimb nail - number

##### 3.3.40 Hindlimb nail - shape

##### 3.3.41 Hindlimbs - position

##### 3.3.42 Hindlimbs - shape

##### 3.3.43 Hindlimbs - size

##### 3.3.44 Hindpaw - shape

##### 3.3.45 Hindpaw - size

##### 3.3.46 Limb grasp

##### 3.3.47 Lower lip morphology

##### 3.3.48 Lower teeth appearance

##### 3.3.49 Mouth morphology

##### 3.3.50 Skin color - back paws

##### 3.3.51 Skin color - ear

##### 3.3.52 Skin color - front paws

##### 3.3.53 Skin color - snout

##### 3.3.54 Skin color - tail

##### 3.3.55 Skin color - whole body

##### 3.3.56 Skin texture - back paws

##### 3.3.57 Skin texture - ear

##### 3.3.58 Skin texture - front paws

##### 3.3.59 Skin texture - snout

##### 3.3.60 Skin texture - tail

##### 3.3.61 Skin texture - whole body

##### 3.3.62 Snout size

##### 3.3.63 Startle response

##### 3.3.64 Tail - length

##### 3.3.65 Tail - morphology

##### 3.3.66 Tail - presence

##### 3.3.67 Tail - thickness

##### 3.3.68 Tail elevation

##### 3.3.69 Teeth presence

##### 3.3.70 Transfer arousal

|  | female WT/WT | female Δ HEL/WT | male WT/WT |
| --- | --- | --- | --- |
| female Δ HEL/WT | 1 | NA | NA |
| male WT/WT | 1 | 1 | NA |
| male Δ HEL/WT | 1 | 1 | 1 |

##### 3.3.71 Tremor

##### 3.3.72 Trunk curl

##### 3.3.73 Unexpected behaviors

##### 3.3.74 Upper lip morphology

##### 3.3.75 Upper teeth appearance

##### 3.3.76 Vibrissae - appearance

##### 3.3.77 Vibrissae - presence

##### 3.3.78 Vocalization

#### 3.4 Grip Strength

**Number of animals**

| STRAIN | GENDER | count |
| --- | --- | --- |
| WT/WT | female | 8 |
| WT/WT | male | 8 |
| Δ HEL/WT | female | 8 |
| Δ HEL/WT | male | 8 |

|  | test | Difference | p\_value | Significance |
| --- | --- | --- | --- | --- |
| c2 | WT/WT vs Δ HEL/WT | -1.8908 | 0.9975 | FALSE |
| c3 | Female vs Male | -26.1839 | 0.0466 | TRUE |
| c4 | Female WT/WT vs Female Δ HEL/WT | 7.1084 | 0.9565 | FALSE |
| c5 | Male WT/WT vs Male Δ HEL/WT | -10.8900 | 0.8629 | FALSE |
| c6 | Female Δ HEL/WT vs Male Δ HEL/WT | -35.1831 | 0.0645 | FALSE |

|  | test | Difference | p\_value | Significance |
| --- | --- | --- | --- | --- |
| c2 | WT/WT vs Δ HEL/WT | 24.9873 | 0.1319 | FALSE |
| c3 | Female vs Male | -35.8054 | 0.0115 | TRUE |
| c4 | Female WT/WT vs Female Δ HEL/WT | 7.4437 | 0.9665 | FALSE |
| c5 | Male WT/WT vs Male Δ HEL/WT | 42.5309 | 0.0467 | TRUE |
| c6 | Female Δ HEL/WT vs Male Δ HEL/WT | -18.2618 | 0.6667 | FALSE |

#### 3.5 Light Dark Box

**Number of animals**

| STRAIN | GENDER | count |
| --- | --- | --- |
| WT/WT | female | 8 |
| WT/WT | male | 8 |
| Δ HEL/WT | female | 8 |
| Δ HEL/WT | male | 8 |

##### 3.5.1 Dark side time spent

|  | test | Difference | p\_value | Significance |
| --- | --- | --- | --- | --- |
| c2 | WT/WT vs Δ HEL/WT | 19.4188 | 0.3704 | FALSE |
| c3 | Female vs Male | 13.9813 | 0.6373 | FALSE |
| c4 | Female WT/WT vs Female Δ HEL/WT | 49.5500 | 0.0320 | TRUE |
| c5 | Male WT/WT vs Male Δ HEL/WT | -10.7125 | 0.9151 | FALSE |
| c6 | Female Δ HEL/WT vs Male Δ HEL/WT | -16.1500 | 0.7645 | FALSE |

##### 3.5.2 Light side time spent

|  | test | Difference | p\_value | Significance |
| --- | --- | --- | --- | --- |
| c2 | WT/WT vs Δ HEL/WT | -19.8937 | 0.3647 | FALSE |
| c3 | Female vs Male | -17.3438 | 0.4812 | FALSE |
| c4 | Female WT/WT vs Female Δ HEL/WT | -50.0000 | 0.0340 | TRUE |
| c5 | Male WT/WT vs Male Δ HEL/WT | 10.2125 | 0.9287 | FALSE |
| c6 | Female Δ HEL/WT vs Male Δ HEL/WT | 12.7625 | 0.8720 | FALSE |

##### 3.5.3 Latency to first transition

|  | test | Difference | p\_value | Significance |
| --- | --- | --- | --- | --- |
| c2 | WT/WT vs Δ HEL/WT | 3.2437 | 0.8456 | FALSE |
| c3 | Female vs Male | 1.7688 | 0.9697 | FALSE |
| c4 | Female WT/WT vs Female Δ HEL/WT | 11.6500 | 0.1952 | FALSE |
| c5 | Male WT/WT vs Male Δ HEL/WT | -5.1625 | 0.7947 | FALSE |
| c6 | Female Δ HEL/WT vs Male Δ HEL/WT | -6.6375 | 0.6449 | FALSE |

##### 3.5.4 Side changes

|  | test | Difference | p\_value | Significance |
| --- | --- | --- | --- | --- |
| c2 | WT/WT vs Δ HEL/WT | -1.4375 | 0.9746 | FALSE |
| c3 | Female vs Male | -3.1875 | 0.7901 | FALSE |
| c4 | Female WT/WT vs Female Δ HEL/WT | -13.7500 | 0.0454 | TRUE |
| c5 | Male WT/WT vs Male Δ HEL/WT | 10.8750 | 0.1464 | FALSE |
| c6 | Female Δ HEL/WT vs Male Δ HEL/WT | 9.1250 | 0.2684 | FALSE |

##### 3.5.5 Time mobile active dark side

|  | test | Difference | p\_value | Significance |
| --- | --- | --- | --- | --- |
| c2 | WT/WT vs Δ HEL/WT | 7.6271 | 0.2871 | FALSE |
| c3 | Female vs Male | 3.1730 | 0.8696 | FALSE |
| c4 | Female WT/WT vs Female Δ HEL/WT | 6.9588 | 0.6437 | FALSE |
| c5 | Male WT/WT vs Male Δ HEL/WT | 8.2953 | 0.5074 | FALSE |
| c6 | Female Δ HEL/WT vs Male Δ HEL/WT | 3.8413 | 0.9129 | FALSE |

##### 3.5.6 Time mobile active light side

|  | test | Difference | p\_value | Significance |
| --- | --- | --- | --- | --- |
| c2 | WT/WT vs Δ HEL/WT | -0.7371 | 0.9988 | FALSE |
| c3 | Female vs Male | -35.7688 | 0.0000 | TRUE |
| c4 | Female WT/WT vs Female Δ HEL/WT | -4.1914 | 0.9336 | FALSE |
| c5 | Male WT/WT vs Male Δ HEL/WT | 2.7172 | 0.9803 | FALSE |
| c6 | Female Δ HEL/WT vs Male Δ HEL/WT | -32.3145 | 0.0006 | TRUE |

#### 3.6 Acustic startle/PPI

**Number of animals**

| STRAIN | GENDER | count |
| --- | --- | --- |
| WT/WT | female | 8 |
| WT/WT | male | 8 |
| Δ HEL/WT | female | 8 |
| Δ HEL/WT | male | 8 |

##### 3.6.1 % Pre-pulse inhibition - Global

|  | test | Difference | p\_value | Significance |
| --- | --- | --- | --- | --- |
| c2 | WT/WT vs Δ HEL/WT | 11.1966 | 0.6283 | FALSE |
| c3 | Female vs Male | -13.2912 | 0.4881 | FALSE |
| c4 | Female WT/WT vs Female Δ HEL/WT | 20.6555 | 0.4047 | FALSE |
| c5 | Male WT/WT vs Male Δ HEL/WT | 1.7377 | 0.9992 | FALSE |
| c6 | Female Δ HEL/WT vs Male Δ HEL/WT | -22.7501 | 0.3191 | FALSE |

##### 3.6.2 % Pre-pulse inhibition - PPI1

|  | test | Difference | p\_value | Significance |
| --- | --- | --- | --- | --- |
| c2 | WT/WT vs Δ HEL/WT | 9.4707 | 0.8267 | FALSE |
| c3 | Female vs Male | -7.7241 | 0.8970 | FALSE |
| c4 | Female WT/WT vs Female Δ HEL/WT | 24.9917 | 0.3922 | FALSE |
| c5 | Male WT/WT vs Male Δ HEL/WT | -6.0503 | 0.9806 | FALSE |
| c6 | Female Δ HEL/WT vs Male Δ HEL/WT | -23.2451 | 0.4595 | FALSE |

##### 3.6.3 % Pre-pulse inhibition - PPI2

|  | test | Difference | p\_value | Significance |
| --- | --- | --- | --- | --- |
| c2 | WT/WT vs Δ HEL/WT | 11.7113 | 0.4912 | FALSE |
| c3 | Female vs Male | -13.2830 | 0.3795 | FALSE |
| c4 | Female WT/WT vs Female Δ HEL/WT | 20.8892 | 0.2863 | FALSE |
| c5 | Male WT/WT vs Male Δ HEL/WT | 2.5335 | 0.9963 | FALSE |
| c6 | Female Δ HEL/WT vs Male Δ HEL/WT | -22.4608 | 0.2268 | FALSE |

##### 3.6.4 % Pre-pulse inhibition - PPI3

|  | test | Difference | p\_value | Significance |
| --- | --- | --- | --- | --- |
| c2 | WT/WT vs Δ HEL/WT | 16.3316 | 0.5345 | FALSE |
| c3 | Female vs Male | -18.1929 | 0.4408 | FALSE |
| c4 | Female WT/WT vs Female Δ HEL/WT | 24.1389 | 0.4969 | FALSE |
| c5 | Male WT/WT vs Male Δ HEL/WT | 8.5242 | 0.9591 | FALSE |
| c6 | Female Δ HEL/WT vs Male Δ HEL/WT | -26.0002 | 0.4314 | FALSE |

##### 3.6.5 % Pre-pulse inhibition - PPI4

|  | test | Difference | p\_value | Significance |
| --- | --- | --- | --- | --- |
| c2 | WT/WT vs Δ HEL/WT | 7.7521 | 0.7965 | FALSE |
| c3 | Female vs Male | -13.4853 | 0.3899 | FALSE |
| c4 | Female WT/WT vs Female Δ HEL/WT | 14.3288 | 0.6311 | FALSE |
| c5 | Male WT/WT vs Male Δ HEL/WT | 1.1755 | 0.9997 | FALSE |
| c6 | Female Δ HEL/WT vs Male Δ HEL/WT | -20.0620 | 0.3446 | FALSE |

#### 3.7 Fear Conditioning

**Number of animals**

| STRAIN | GENDER | count |
| --- | --- | --- |
| WT/WT | female | 8 |
| WT/WT | male | 8 |
| Δ HEL/WT | female | 8 |
| Δ HEL/WT | male | 8 |

#### 3.8 Indirect Calorimetry

**Number of animals**

| STRAIN | GENDER | count |
| --- | --- | --- |
| WT/WT | male | 7 |
| Δ HEL/WT | male | 8 |

##### 3.8.1 Oxygen consumption

| test | Difference | p\_value | Significance |
| --- | --- | --- | --- |
| CWT/WT vs Δ HEL/WT | 126.2974 | 0.3333 | FALSE |

##### 3.8.2 Respiratory Exchange Ratio

| test | Difference | p\_value | Significance |
| --- | --- | --- | --- |
| CWT/WT vs Δ HEL/WT | 0.2025 | 0.2867 | FALSE |

##### 3.8.3 Heat production (metabolic rate)

| test | Difference | p\_value | Significance |
| --- | --- | --- | --- |
| CWT/WT vs Δ HEL/WT | 1.4017 | 0.4576 | FALSE |

##### 3.8.4 Ambulatory activity (no. of beam cuts)

| test | Difference | p\_value | Significance |
| --- | --- | --- | --- |
| CWT/WT vs Δ HEL/WT | 26.5653 | 0.0394 | TRUE |

##### 3.8.5 Total movement

| test | Difference | p\_value | Significance |
| --- | --- | --- | --- |
| CWT/WT vs Δ HEL/WT | 307.636 | 0.0091 | TRUE |

##### 3.8.6 Cumulative food intake

| test | Difference | p\_value | Significance |
| --- | --- | --- | --- |
| CWT/WT vs Δ HEL/WT | -0.807 | 0.8863 | FALSE |

##### 3.8.7 Cumulative water intake

| test | Difference | p\_value | Significance |
| --- | --- | --- | --- |
| CWT/WT vs Δ HEL/WT | -5.9333 | 0.4876 | FALSE |

##### 3.8.8 Carbon dioxide production per kg

| test | Difference | p\_value | Significance |
| --- | --- | --- | --- |
| CWT/WT vs Δ HEL/WT | 33.0785 | 0.868 | FALSE |

##### 3.8.9 Oxygen consumption per kg

| test | Difference | p\_value | Significance |
| --- | --- | --- | --- |
| CWT/WT vs Δ HEL/WT | 53.074 | 0.8166 | FALSE |

##### 3.8.10 DistD

| test | Difference | p\_value | Significance |
| --- | --- | --- | --- |
| CWT/WT vs Δ HEL/WT | 848.8899 | 0.0678 | FALSE |

##### 3.8.11 DistK

| test | Difference | p\_value | Significance |
| --- | --- | --- | --- |
| CWT/WT vs Δ HEL/WT | 69588.7 | 0.0042 | TRUE |

### 3.8.12 Z

| test | Difference | p\_value | Significance |
| --- | --- | --- | --- |
| CWT/WT vs Δ HEL/WT | 62.8633 | 0.0309 | TRUE |

#### 3.9 Echocardiography

**Number of animals**

| STRAIN | GENDER | count |
| --- | --- | --- |
| WT/WT | female | 8 |
| WT/WT | male | 8 |
| Δ HEL/WT | female | 7 |
| Δ HEL/WT | male | 8 |

##### 3.9.1 Aortic diameter (Dao)

|  | test | Difference | p\_value | Significance |
| --- | --- | --- | --- | --- |
| c2 | WT/WT vs Δ HEL/WT | 0.0413 | 0.5249 | FALSE |
| c3 | Female vs Male | 0.0295 | 0.8560 | FALSE |
| c4 | Female WT/WT vs Female Δ HEL/WT | 0.0398 | 0.7954 | FALSE |
| c5 | Male WT/WT vs Male Δ HEL/WT | 0.0427 | 0.7400 | FALSE |
| c6 | Female Δ HEL/WT vs Male Δ HEL/WT | 0.0309 | 0.9193 | FALSE |

##### 3.9.2 Cardiac Output

|  | test | Difference | p\_value | Significance |
| --- | --- | --- | --- | --- |
| c2 | WT/WT vs Δ HEL/WT | 1.9593 | 0.5580 | FALSE |
| c3 | Female vs Male | -5.0347 | 0.5328 | FALSE |
| c4 | Female WT/WT vs Female Δ HEL/WT | 2.9704 | 0.5138 | FALSE |
| c5 | Male WT/WT vs Male Δ HEL/WT | 0.9482 | 0.9679 | FALSE |
| c6 | Female Δ HEL/WT vs Male Δ HEL/WT | -6.0458 | 0.4427 | FALSE |

##### 3.9.3 Ejection Fraction

|  | test | Difference | p\_value | Significance |
| --- | --- | --- | --- | --- |
| c2 | WT/WT vs Δ HEL/WT | -1.2780 | 0.9543 | FALSE |
| c3 | Female vs Male | 2.2724 | 0.8647 | FALSE |
| c4 | Female WT/WT vs Female Δ HEL/WT | 5.2137 | 0.4616 | FALSE |
| c5 | Male WT/WT vs Male Δ HEL/WT | -7.7696 | 0.1160 | FALSE |
| c6 | Female Δ HEL/WT vs Male Δ HEL/WT | -4.2192 | 0.6986 | FALSE |

##### 3.9.4 End-Diastolic Diameter

|  | test | Difference | p\_value | Significance |
| --- | --- | --- | --- | --- |
| c2 | WT/WT vs Δ HEL/WT | 0.2093 | 0.1172 | FALSE |
| c3 | Female vs Male | -0.3118 | 0.5636 | FALSE |
| c4 | Female WT/WT vs Female Δ HEL/WT | 0.0494 | 0.9824 | FALSE |
| c5 | Male WT/WT vs Male Δ HEL/WT | 0.3693 | 0.0263 | TRUE |
| c6 | Female Δ HEL/WT vs Male Δ HEL/WT | -0.1518 | 0.9348 | FALSE |

##### 3.9.5 End-Systolic Diameter

|  | test | Difference | p\_value | Significance |
| --- | --- | --- | --- | --- |
| c2 | WT/WT vs Δ HEL/WT | 0.1850 | 0.3036 | FALSE |
| c3 | Female vs Male | -0.2809 | 0.7866 | FALSE |
| c4 | Female WT/WT vs Female Δ HEL/WT | -0.1088 | 0.8892 | FALSE |
| c5 | Male WT/WT vs Male Δ HEL/WT | 0.4789 | 0.0084 | TRUE |
| c6 | Female Δ HEL/WT vs Male Δ HEL/WT | 0.0129 | 1.0000 | FALSE |

##### 3.9.6 Fractional Shortening

|  | test | Difference | p\_value | Significance |
| --- | --- | --- | --- | --- |
| c2 | WT/WT vs Δ HEL/WT | -0.7206 | 0.9684 | FALSE |
| c3 | Female vs Male | 1.3830 | 0.8782 | FALSE |
| c4 | Female WT/WT vs Female Δ HEL/WT | 3.4105 | 0.4457 | FALSE |
| c5 | Male WT/WT vs Male Δ HEL/WT | -4.8517 | 0.1326 | FALSE |
| c6 | Female Δ HEL/WT vs Male Δ HEL/WT | -2.7481 | 0.6846 | FALSE |

##### 3.9.7 Heart rate

|  | test | Difference | p\_value | Significance |
| --- | --- | --- | --- | --- |
| c2 | WT/WT vs Δ HEL/WT | 0.9054 | 1.0000 | FALSE |
| c3 | Female vs Male | -52.1438 | 0.5883 | FALSE |
| c4 | Female WT/WT vs Female Δ HEL/WT | 6.4704 | 0.9945 | FALSE |
| c5 | Male WT/WT vs Male Δ HEL/WT | -4.6595 | 0.9977 | FALSE |
| c6 | Female Δ HEL/WT vs Male Δ HEL/WT | -57.7088 | 0.5818 | FALSE |

##### 3.9.8 Left ventricle anterior wall (diastole)

|  | test | Difference | p\_value | Significance |
| --- | --- | --- | --- | --- |
| c2 | WT/WT vs Δ HEL/WT | -0.0297 | 0.8977 | FALSE |
| c3 | Female vs Male | 0.0287 | 0.9283 | FALSE |
| c4 | Female WT/WT vs Female Δ HEL/WT | -0.0187 | 0.9901 | FALSE |
| c5 | Male WT/WT vs Male Δ HEL/WT | -0.0407 | 0.9012 | FALSE |
| c6 | Female Δ HEL/WT vs Male Δ HEL/WT | 0.0177 | 0.9927 | FALSE |

##### 3.9.9 Left ventricle anterior wall (systole)

|  | test | Difference | p\_value | Significance |
| --- | --- | --- | --- | --- |
| c2 | WT/WT vs Δ HEL/WT | -0.0328 | 0.9196 | FALSE |
| c3 | Female vs Male | 0.0432 | 0.8882 | FALSE |
| c4 | Female WT/WT vs Female Δ HEL/WT | 0.0659 | 0.8106 | FALSE |
| c5 | Male WT/WT vs Male Δ HEL/WT | -0.1315 | 0.2662 | FALSE |
| c6 | Female Δ HEL/WT vs Male Δ HEL/WT | -0.0555 | 0.8997 | FALSE |

##### 3.9.10 Left ventricle interior diameter (diastole)

|  | test | Difference | p\_value | Significance |
| --- | --- | --- | --- | --- |
| c2 | WT/WT vs Δ HEL/WT | 0.2352 | 0.0971 | FALSE |
| c3 | Female vs Male | -0.2409 | 0.5384 | FALSE |
| c4 | Female WT/WT vs Female Δ HEL/WT | 0.0248 | 0.9982 | FALSE |
| c5 | Male WT/WT vs Male Δ HEL/WT | 0.4457 | 0.0102 | TRUE |
| c6 | Female Δ HEL/WT vs Male Δ HEL/WT | -0.0305 | 0.9988 | FALSE |

##### 3.9.11 Left ventricle interior diameter (systole)

|  | test | Difference | p\_value | Significance |
| --- | --- | --- | --- | --- |
| c2 | WT/WT vs Δ HEL/WT | 0.2234 | 0.3106 | FALSE |
| c3 | Female vs Male | -0.1744 | 0.7280 | FALSE |
| c4 | Female WT/WT vs Female Δ HEL/WT | -0.1575 | 0.8278 | FALSE |
| c5 | Male WT/WT vs Male Δ HEL/WT | 0.6043 | 0.0054 | TRUE |
| c6 | Female Δ HEL/WT vs Male Δ HEL/WT | 0.2065 | 0.7697 | FALSE |

##### 3.9.12 Left ventricle posterior wall (diastole)

|  | test | Difference | p\_value | Significance |
| --- | --- | --- | --- | --- |
| c2 | WT/WT vs Δ HEL/WT | 0.0234 | 0.9727 | FALSE |
| c3 | Female vs Male | -0.0977 | 0.6470 | FALSE |
| c4 | Female WT/WT vs Female Δ HEL/WT | 0.1245 | 0.3876 | FALSE |
| c5 | Male WT/WT vs Male Δ HEL/WT | -0.0777 | 0.7313 | FALSE |
| c6 | Female Δ HEL/WT vs Male Δ HEL/WT | -0.1988 | 0.2066 | FALSE |

##### 3.9.13 Left ventricle posterior wall (systole)

|  | test | Difference | p\_value | Significance |
| --- | --- | --- | --- | --- |
| c2 | WT/WT vs Δ HEL/WT | -0.0409 | 0.9520 | FALSE |
| c3 | Female vs Male | -0.1064 | 0.8706 | FALSE |
| c4 | Female WT/WT vs Female Δ HEL/WT | 0.1464 | 0.5579 | FALSE |
| c5 | Male WT/WT vs Male Δ HEL/WT | -0.2281 | 0.1578 | FALSE |
| c6 | Female Δ HEL/WT vs Male Δ HEL/WT | -0.2936 | 0.2708 | FALSE |

##### 3.9.14 Stroke Volume

|  | test | Difference | p\_value | Significance |
| --- | --- | --- | --- | --- |
| c2 | WT/WT vs Δ HEL/WT | 4.0597 | 0.3638 | FALSE |
| c3 | Female vs Male | -5.8412 | 0.5835 | FALSE |
| c4 | Female WT/WT vs Female Δ HEL/WT | 5.5459 | 0.4099 | FALSE |
| c5 | Male WT/WT vs Male Δ HEL/WT | 2.5734 | 0.8769 | FALSE |
| c6 | Female Δ HEL/WT vs Male Δ HEL/WT | -7.3275 | 0.5075 | FALSE |

#### 3.10 ECG

**Number of animals**

| STRAIN | GENDER | count |
| --- | --- | --- |
| WT/WT | female | 8 |
| WT/WT | male | 8 |
| Δ HEL/WT | female | 7 |
| Δ HEL/WT | male | 7 |

##### 3.10.1 Heart rate

|  | test | Difference | p\_value | Significance |
| --- | --- | --- | --- | --- |
| c2 | WT/WT vs Δ HEL/WT | 14.2857 | 0.8952 | FALSE |
| c3 | Female vs Male | -50.1964 | 0.0993 | FALSE |
| c4 | Female WT/WT vs Female Δ HEL/WT | 14.2321 | 0.9594 | FALSE |
| c5 | Male WT/WT vs Male Δ HEL/WT | 14.3393 | 0.9585 | FALSE |
| c6 | Female Δ HEL/WT vs Male Δ HEL/WT | -50.1429 | 0.3605 | FALSE |

##### 3.10.2 HRV

|  | test | Difference | p\_value | Significance |
| --- | --- | --- | --- | --- |
| c2 | WT/WT vs Δ HEL/WT | -3.8089 | 0.7640 | FALSE |
| c3 | Female vs Male | 8.8911 | 0.1387 | FALSE |
| c4 | Female WT/WT vs Female Δ HEL/WT | -1.0750 | 0.9973 | FALSE |
| c5 | Male WT/WT vs Male Δ HEL/WT | -6.5429 | 0.6428 | FALSE |
| c6 | Female Δ HEL/WT vs Male Δ HEL/WT | 6.1571 | 0.7050 | FALSE |

##### 3.10.3 Mean R amplitude

|  | test | Difference | p\_value | Significance |
| --- | --- | --- | --- | --- |
| c2 | WT/WT vs Δ HEL/WT | -0.0258 | 0.9793 | FALSE |
| c3 | Female vs Male | -0.0119 | 0.9979 | FALSE |
| c4 | Female WT/WT vs Female Δ HEL/WT | -0.0577 | 0.9257 | FALSE |
| c5 | Male WT/WT vs Male Δ HEL/WT | 0.0061 | 0.9999 | FALSE |
| c6 | Female Δ HEL/WT vs Male Δ HEL/WT | 0.0200 | 0.9968 | FALSE |

##### 3.10.4 Mean SR amplitude

|  | test | Difference | p\_value | Significance |
| --- | --- | --- | --- | --- |
| c2 | WT/WT vs Δ HEL/WT | -0.0391 | 0.9886 | FALSE |
| c3 | Female vs Male | 0.0654 | 0.9507 | FALSE |
| c4 | Female WT/WT vs Female Δ HEL/WT | -0.0395 | 0.9958 | FALSE |
| c5 | Male WT/WT vs Male Δ HEL/WT | -0.0388 | 0.9960 | FALSE |
| c6 | Female Δ HEL/WT vs Male Δ HEL/WT | 0.0657 | 0.9829 | FALSE |

### 3.10.5 PQ

|  | test | Difference | p\_value | Significance |
| --- | --- | --- | --- | --- |
| c2 | WT/WT vs Δ HEL/WT | -0.1054 | 0.9994 | FALSE |
| c3 | Female vs Male | 1.8018 | 0.2346 | FALSE |
| c4 | Female WT/WT vs Female Δ HEL/WT | 0.1679 | 0.9992 | FALSE |
| c5 | Male WT/WT vs Male Δ HEL/WT | -0.3786 | 0.9909 | FALSE |
| c6 | Female Δ HEL/WT vs Male Δ HEL/WT | 1.5286 | 0.6677 | FALSE |

### 3.10.6 PR

|  | test | Difference | p\_value | Significance |
| --- | --- | --- | --- | --- |
| c2 | WT/WT vs Δ HEL/WT | -0.1295 | 0.9995 | FALSE |
| c3 | Female vs Male | 1.8170 | 0.4183 | FALSE |
| c4 | Female WT/WT vs Female Δ HEL/WT | 0.0161 | 1.0000 | FALSE |
| c5 | Male WT/WT vs Male Δ HEL/WT | -0.2750 | 0.9982 | FALSE |
| c6 | Female Δ HEL/WT vs Male Δ HEL/WT | 1.6714 | 0.7565 | FALSE |

##### 3.10.7 QRS

|  | test | Difference | p\_value | Significance |
| --- | --- | --- | --- | --- |
| c2 | WT/WT vs Δ HEL/WT | -0.1134 | 0.9783 | FALSE |
| c3 | Female vs Male | -0.0812 | 0.9917 | FALSE |
| c4 | Female WT/WT vs Female Δ HEL/WT | 0.1054 | 0.9936 | FALSE |
| c5 | Male WT/WT vs Male Δ HEL/WT | -0.3321 | 0.8444 | FALSE |
| c6 | Female Δ HEL/WT vs Male Δ HEL/WT | -0.3000 | 0.8891 | FALSE |

### 3.10.8 QT

|  | test | Difference | p\_value | Significance |
| --- | --- | --- | --- | --- |
| c2 | WT/WT vs Δ HEL/WT | -0.6661 | 0.9536 | FALSE |
| c3 | Female vs Male | 2.8036 | 0.1661 | FALSE |
| c4 | Female WT/WT vs Female Δ HEL/WT | -0.1196 | 0.9999 | FALSE |
| c5 | Male WT/WT vs Male Δ HEL/WT | -1.2125 | 0.9082 | FALSE |
| c6 | Female Δ HEL/WT vs Male Δ HEL/WT | 2.2571 | 0.6343 | FALSE |

##### 3.10.9 QT\_dispersion

|  | test | Difference | p\_value | Significance |
| --- | --- | --- | --- | --- |
| c2 | WT/WT vs Δ HEL/WT | -0.0964 | 0.9999 | FALSE |
| c3 | Female vs Male | 5.7964 | 0.0090 | TRUE |
| c4 | Female WT/WT vs Female Δ HEL/WT | 0.8571 | 0.9812 | FALSE |
| c5 | Male WT/WT vs Male Δ HEL/WT | -1.0500 | 0.9665 | FALSE |
| c6 | Female Δ HEL/WT vs Male Δ HEL/WT | 4.8429 | 0.2043 | FALSE |

##### 3.10.10 QTc

|  | test | Difference | p\_value | Significance |
| --- | --- | --- | --- | --- |
| c2 | WT/WT vs Δ HEL/WT | -0.2902 | 0.9875 | FALSE |
| c3 | Female vs Male | 1.4330 | 0.3972 | FALSE |
| c4 | Female WT/WT vs Female Δ HEL/WT | 0.3143 | 0.9943 | FALSE |
| c5 | Male WT/WT vs Male Δ HEL/WT | -0.8946 | 0.8909 | FALSE |
| c6 | Female Δ HEL/WT vs Male Δ HEL/WT | 0.8286 | 0.9180 | FALSE |

##### 3.10.11 QTc Dispersion

|  | test | Difference | p\_value | Significance |
| --- | --- | --- | --- | --- |
| c2 | WT/WT vs Δ HEL/WT | 0.1982 | 0.9991 | FALSE |
| c3 | Female vs Male | 5.1554 | 0.0106 | TRUE |
| c4 | Female WT/WT vs Female Δ HEL/WT | 0.9679 | 0.9655 | FALSE |
| c5 | Male WT/WT vs Male Δ HEL/WT | -0.5714 | 0.9924 | FALSE |
| c6 | Female Δ HEL/WT vs Male Δ HEL/WT | 4.3857 | 0.2087 | FALSE |

### 3.10.12 RR

|  | test | Difference | p\_value | Significance |
| --- | --- | --- | --- | --- |
| c2 | WT/WT vs Δ HEL/WT | -1.5911 | 0.9164 | FALSE |
| c3 | Female vs Male | 5.9786 | 0.1083 | FALSE |
| c4 | Female WT/WT vs Female Δ HEL/WT | -1.6696 | 0.9631 | FALSE |
| c5 | Male WT/WT vs Male Δ HEL/WT | -1.5125 | 0.9721 | FALSE |
| c6 | Female Δ HEL/WT vs Male Δ HEL/WT | 6.0571 | 0.3648 | FALSE |

#### 3.11 Lung screen

**Number of animals**

| STRAIN | GENDER | count |
| --- | --- | --- |
| WT/WT | female | 8 |
| WT/WT | male | 8 |
| Δ HEL/WT | female | 8 |
| Δ HEL/WT | male | 8 |

##### 3.11.1 Compliance

|  | test | Difference | p\_value | Significance |
| --- | --- | --- | --- | --- |
| c2 | WT/WT vs Δ HEL/WT | 0.0043 | 0.3763 | FALSE |
| c3 | Female vs Male | 0.0011 | 0.9859 | FALSE |
| c4 | Female WT/WT vs Female Δ HEL/WT | 0.0060 | 0.2549 | FALSE |
| c5 | Male WT/WT vs Male Δ HEL/WT | 0.0026 | 0.8633 | FALSE |
| c6 | Female Δ HEL/WT vs Male Δ HEL/WT | -0.0005 | 0.9988 | FALSE |

##### 3.11.2 Elastance

|  | test | Difference | p\_value | Significance |
| --- | --- | --- | --- | --- |
| c2 | WT/WT vs Δ HEL/WT | -1.0075 | 0.4927 | FALSE |
| c3 | Female vs Male | -0.1942 | 0.9962 | FALSE |
| c4 | Female WT/WT vs Female Δ HEL/WT | -1.4336 | 0.3471 | FALSE |
| c5 | Male WT/WT vs Male Δ HEL/WT | -0.5814 | 0.9182 | FALSE |
| c6 | Female Δ HEL/WT vs Male Δ HEL/WT | 0.2319 | 0.9954 | FALSE |

##### 3.11.3 Inspiratory Capacity

|  | test | Difference | p\_value | Significance |
| --- | --- | --- | --- | --- |
| c2 | WT/WT vs Δ HEL/WT | 0.0366 | 0.7723 | FALSE |
| c3 | Female vs Male | 0.0257 | 0.9529 | FALSE |
| c4 | Female WT/WT vs Female Δ HEL/WT | 0.0365 | 0.8493 | FALSE |
| c5 | Male WT/WT vs Male Δ HEL/WT | 0.0367 | 0.8832 | FALSE |
| c6 | Female Δ HEL/WT vs Male Δ HEL/WT | 0.0258 | 0.9648 | FALSE |

##### 3.11.4 Resistance

|  | test | Difference | p\_value | Significance |
| --- | --- | --- | --- | --- |
| c2 | WT/WT vs Δ HEL/WT | 0.0191 | 0.8756 | FALSE |
| c3 | Female vs Male | -0.0265 | 0.8560 | FALSE |
| c4 | Female WT/WT vs Female Δ HEL/WT | 0.0403 | 0.5675 | FALSE |
| c5 | Male WT/WT vs Male Δ HEL/WT | -0.0021 | 0.9999 | FALSE |
| c6 | Female Δ HEL/WT vs Male Δ HEL/WT | -0.0477 | 0.5927 | FALSE |

##### 3.11.5 Resistance of the airways (Prime8)

|  | test | Difference | p\_value | Significance |
| --- | --- | --- | --- | --- |
| c2 | WT/WT vs Δ HEL/WT | 0.0303 | 0.3004 | FALSE |
| c3 | Female vs Male | -0.0238 | 0.6902 | FALSE |
| c4 | Female WT/WT vs Female Δ HEL/WT | 0.0388 | 0.2441 | FALSE |
| c5 | Male WT/WT vs Male Δ HEL/WT | 0.0218 | 0.7508 | FALSE |
| c6 | Female Δ HEL/WT vs Male Δ HEL/WT | -0.0323 | 0.5499 | FALSE |

##### 3.11.6 Resistance of the airways (QuickPrime3)

|  | test | Difference | p\_value | Significance |
| --- | --- | --- | --- | --- |
| c2 | WT/WT vs Δ HEL/WT | 0.0359 | 0.2681 | FALSE |
| c3 | Female vs Male | 0.0008 | 1.0000 | FALSE |
| c4 | Female WT/WT vs Female Δ HEL/WT | 0.0425 | 0.2710 | FALSE |
| c5 | Male WT/WT vs Male Δ HEL/WT | 0.0292 | 0.6477 | FALSE |
| c6 | Female Δ HEL/WT vs Male Δ HEL/WT | -0.0059 | 0.9962 | FALSE |

##### 3.11.7 Tissue damping (Prime8)

|  | test | Difference | p\_value | Significance |
| --- | --- | --- | --- | --- |
| c2 | WT/WT vs Δ HEL/WT | -0.1933 | 0.2515 | FALSE |
| c3 | Female vs Male | -0.1052 | 0.8424 | FALSE |
| c4 | Female WT/WT vs Female Δ HEL/WT | -0.2624 | 0.1630 | FALSE |
| c5 | Male WT/WT vs Male Δ HEL/WT | -0.1241 | 0.7757 | FALSE |
| c6 | Female Δ HEL/WT vs Male Δ HEL/WT | -0.0361 | 0.9939 | FALSE |

##### 3.11.8 Tissue damping (QuickPrime3)

|  | test | Difference | p\_value | Significance |
| --- | --- | --- | --- | --- |
| c2 | WT/WT vs Δ HEL/WT | -0.1222 | 0.4226 | FALSE |
| c3 | Female vs Male | -0.1640 | 0.4005 | FALSE |
| c4 | Female WT/WT vs Female Δ HEL/WT | -0.1577 | 0.3562 | FALSE |
| c5 | Male WT/WT vs Male Δ HEL/WT | -0.0867 | 0.8289 | FALSE |
| c6 | Female Δ HEL/WT vs Male Δ HEL/WT | -0.1285 | 0.6696 | FALSE |

##### 3.11.9 Tissue elastance (Prime8)

|  | test | Difference | p\_value | Significance |
| --- | --- | --- | --- | --- |
| c2 | WT/WT vs Δ HEL/WT | -0.8563 | 0.7318 | FALSE |
| c3 | Female vs Male | -0.4213 | 0.9785 | FALSE |
| c4 | Female WT/WT vs Female Δ HEL/WT | -1.3815 | 0.5207 | FALSE |
| c5 | Male WT/WT vs Male Δ HEL/WT | -0.3311 | 0.9897 | FALSE |
| c6 | Female Δ HEL/WT vs Male Δ HEL/WT | 0.1039 | 0.9998 | FALSE |

##### 3.11.10 Tissue elastance (QuickPrime3)

|  | test | Difference | p\_value | Significance |
| --- | --- | --- | --- | --- |
| c2 | WT/WT vs Δ HEL/WT | -1.1877 | 0.4685 | FALSE |
| c3 | Female vs Male | 0.3121 | 0.9898 | FALSE |
| c4 | Female WT/WT vs Female Δ HEL/WT | -1.6666 | 0.3344 | FALSE |
| c5 | Male WT/WT vs Male Δ HEL/WT | -0.7088 | 0.9034 | FALSE |
| c6 | Female Δ HEL/WT vs Male Δ HEL/WT | 0.7909 | 0.8995 | FALSE |

#### 3.12 Intraperitoneal Glucose Tolerance Test

**Number of animals**

| STRAIN | GENDER | count |
| --- | --- | --- |
| WT/WT | female | 7 |
| WT/WT | male | 7 |
| Δ HEL/WT | female | 7 |
| Δ HEL/WT | male | 6 |

|  | test | Difference | p\_value | Significance |
| --- | --- | --- | --- | --- |
| c2 | WT/WT vs Δ HEL/WT | 3.3349 | 0.9763 | FALSE |
| c3 | Female vs Male | 3.5408 | 0.9871 | FALSE |
| c4 | Female WT/WT vs Female Δ HEL/WT | 13.9165 | 0.4830 | FALSE |
| c5 | Male WT/WT vs Male Δ HEL/WT | -7.2468 | 0.9094 | FALSE |
| c6 | Female Δ HEL/WT vs Male Δ HEL/WT | -7.0409 | 0.9313 | FALSE |

[1] 0

##### 3.12.1 AUC

|  | test | Difference | p\_value | Significance |
| --- | --- | --- | --- | --- |
| c2 | WT/WT vs Δ HEL/WT | 948.3586 | 0.3963 | FALSE |
| c3 | Female vs Male | -607.6252 | 0.8253 | FALSE |
| c4 | Female WT/WT vs Female Δ HEL/WT | 2322.3928 | 0.0325 | TRUE |
| c5 | Male WT/WT vs Male Δ HEL/WT | -425.6755 | 0.9604 | FALSE |
| c6 | Female Δ HEL/WT vs Male Δ HEL/WT | -1981.6594 | 0.1629 | FALSE |

#### 3.13 Auditory Brain Stem Response

**Number of animals**

| STRAIN | GENDER | count |
| --- | --- | --- |
| WT/WT | female | 8 |
| WT/WT | male | 8 |
| Δ HEL/WT | female | 7 |
| Δ HEL/WT | male | 7 |

##### 3.13.1 Click-evoked ABR threshold

|  | test | Difference | p\_value | Significance |
| --- | --- | --- | --- | --- |
| c2 | WT/WT vs Δ HEL/WT | 0.7397 | 0.9941 | FALSE |
| c3 | Female vs Male | 2.4166 | 0.8310 | FALSE |
| c4 | Female WT/WT vs Female Δ HEL/WT | 2.0730 | 0.9570 | FALSE |
| c5 | Male WT/WT vs Male Δ HEL/WT | -0.5937 | 0.9989 | FALSE |
| c6 | Female Δ HEL/WT vs Male Δ HEL/WT | 1.0833 | 0.9938 | FALSE |

##### 3.13.2 6kHz-evoked ABR Threshold

|  | test | Difference | p\_value | Significance |
| --- | --- | --- | --- | --- |
| c2 | WT/WT vs Δ HEL/WT | -5.1786 | 0.1808 | FALSE |
| c3 | Female vs Male | -6.5179 | 0.0565 | FALSE |
| c4 | Female WT/WT vs Female Δ HEL/WT | -2.4107 | 0.9058 | FALSE |
| c5 | Male WT/WT vs Male Δ HEL/WT | -7.9464 | 0.1273 | FALSE |
| c6 | Female Δ HEL/WT vs Male Δ HEL/WT | -9.2857 | 0.0662 | FALSE |

##### 3.13.3 12kHz-evoked ABR Threshold

|  | test | Difference | p\_value | Significance |
| --- | --- | --- | --- | --- |
| c2 | WT/WT vs Δ HEL/WT | 7.4754 | 0.6537 | FALSE |
| c3 | Female vs Male | -6.2340 | 0.7587 | FALSE |
| c4 | Female WT/WT vs Female Δ HEL/WT | 10.3399 | 0.6775 | FALSE |
| c5 | Male WT/WT vs Male Δ HEL/WT | 4.6109 | 0.9534 | FALSE |
| c6 | Female Δ HEL/WT vs Male Δ HEL/WT | -9.0985 | 0.7507 | FALSE |

##### 3.13.4 18kHz-evoked ABR Threshold

|  | test | Difference | p\_value | Significance |
| --- | --- | --- | --- | --- |
| c2 | WT/WT vs Δ HEL/WT | 14.2806 | 0.5406 | FALSE |
| c3 | Female vs Male | -11.6980 | 0.6836 | FALSE |
| c4 | Female WT/WT vs Female Δ HEL/WT | 19.3486 | 0.3441 | FALSE |
| c5 | Male WT/WT vs Male Δ HEL/WT | 9.2125 | 0.9535 | FALSE |
| c6 | Female Δ HEL/WT vs Male Δ HEL/WT | -16.7661 | 0.7024 | FALSE |

#### 3.14 Body Composition

**Number of animals**

| STRAIN | GENDER | count |
| --- | --- | --- |
| WT/WT | female | 8 |
| WT/WT | male | 8 |
| Δ HEL/WT | female | 8 |
| Δ HEL/WT | male | 6 |

##### 3.14.1 BMC/Body weight CCP

|  | test | Difference | p\_value | Significance |
| --- | --- | --- | --- | --- |
| c2 | WT/WT vs Δ HEL/WT | -2e-04 | 0.0036 | TRUE |
| c3 | Female vs Male | 4e-04 | 0.0000 | TRUE |
| c4 | Female WT/WT vs Female Δ HEL/WT | -2e-04 | 0.0149 | TRUE |
| c5 | Male WT/WT vs Male Δ HEL/WT | -2e-04 | 0.1305 | FALSE |
| c6 | Female Δ HEL/WT vs Male Δ HEL/WT | 5e-04 | 0.0000 | TRUE |

##### 3.14.2 Bone Area CCP

|  | test | Difference | p\_value | Significance |
| --- | --- | --- | --- | --- |
| c2 | WT/WT vs Δ HEL/WT | 13.8142 | 0.9163 | FALSE |
| c3 | Female vs Male | -62.8955 | 0.0388 | TRUE |
| c4 | Female WT/WT vs Female Δ HEL/WT | 5.6458 | 0.9973 | FALSE |
| c5 | Male WT/WT vs Male Δ HEL/WT | 21.9826 | 0.8965 | FALSE |
| c6 | Female Δ HEL/WT vs Male Δ HEL/WT | -54.7271 | 0.3355 | FALSE |

##### 3.14.3 Bone Mineral Content (excluding skull) CCP

|  | test | Difference | p\_value | Significance |
| --- | --- | --- | --- | --- |
| c2 | WT/WT vs Δ HEL/WT | -0.0011 | 0.9319 | FALSE |
| c3 | Female vs Male | 0.0017 | 0.8029 | FALSE |
| c4 | Female WT/WT vs Female Δ HEL/WT | -0.0016 | 0.9237 | FALSE |
| c5 | Male WT/WT vs Male Δ HEL/WT | -0.0006 | 0.9952 | FALSE |
| c6 | Female Δ HEL/WT vs Male Δ HEL/WT | 0.0021 | 0.8599 | FALSE |

##### 3.14.4 Bone Mineral Density CCP

|  | test | Difference | p\_value | Significance |
| --- | --- | --- | --- | --- |
| c2 | WT/WT vs Δ HEL/WT | -0.0020 | 0.0228 | TRUE |
| c3 | Female vs Male | 0.0057 | 0.0000 | TRUE |
| c4 | Female WT/WT vs Female Δ HEL/WT | -0.0020 | 0.1225 | FALSE |
| c5 | Male WT/WT vs Male Δ HEL/WT | -0.0020 | 0.1742 | FALSE |
| c6 | Female Δ HEL/WT vs Male Δ HEL/WT | 0.0057 | 0.0000 | TRUE |

##### 3.14.5 Fat mass CCP

|  | test | Difference | p\_value | Significance |
| --- | --- | --- | --- | --- |
| c2 | WT/WT vs Δ HEL/WT | 0.2226 | 0.7457 | FALSE |
| c3 | Female vs Male | -0.3445 | 0.4236 | FALSE |
| c4 | Female WT/WT vs Female Δ HEL/WT | 0.3228 | 0.7069 | FALSE |
| c5 | Male WT/WT vs Male Δ HEL/WT | 0.1224 | 0.9809 | FALSE |
| c6 | Female Δ HEL/WT vs Male Δ HEL/WT | -0.4447 | 0.5304 | FALSE |

##### 3.14.6 Fat/Body weight

|  | test | Difference | p\_value | Significance |
| --- | --- | --- | --- | --- |
| c2 | WT/WT vs Δ HEL/WT | 0.0009 | 0.9989 | FALSE |
| c3 | Female vs Male | 0.0024 | 0.9779 | FALSE |
| c4 | Female WT/WT vs Female Δ HEL/WT | 0.0044 | 0.9453 | FALSE |
| c5 | Male WT/WT vs Male Δ HEL/WT | -0.0027 | 0.9892 | FALSE |
| c6 | Female Δ HEL/WT vs Male Δ HEL/WT | -0.0012 | 0.9990 | FALSE |

##### 3.14.7 Lean mass CCP

|  | test | Difference | p\_value | Significance |
| --- | --- | --- | --- | --- |
| c2 | WT/WT vs Δ HEL/WT | 2.3427 | 0.0067 | TRUE |
| c3 | Female vs Male | -5.9169 | 0.0000 | TRUE |
| c4 | Female WT/WT vs Female Δ HEL/WT | 2.1566 | 0.0900 | FALSE |
| c5 | Male WT/WT vs Male Δ HEL/WT | 2.5288 | 0.0589 | FALSE |
| c6 | Female Δ HEL/WT vs Male Δ HEL/WT | -5.7308 | 0.0000 | TRUE |

##### 3.14.8 Lean/Body weight CCP

|  | test | Difference | p\_value | Significance |
| --- | --- | --- | --- | --- |
| c2 | WT/WT vs Δ HEL/WT | 0.0090 | 0.6934 | FALSE |
| c3 | Female vs Male | -0.0349 | 0.0016 | TRUE |
| c4 | Female WT/WT vs Female Δ HEL/WT | 0.0120 | 0.7034 | FALSE |
| c5 | Male WT/WT vs Male Δ HEL/WT | 0.0059 | 0.9585 | FALSE |
| c6 | Female Δ HEL/WT vs Male Δ HEL/WT | -0.0379 | 0.0228 | TRUE |

##### 3.14.9 relative V bone

|  | test | Difference | p\_value | Significance |
| --- | --- | --- | --- | --- |
| c2 | WT/WT vs Δ HEL/WT | -0.3108 | 0.0037 | TRUE |
| c3 | Female vs Male | 0.6713 | 0.0000 | TRUE |
| c4 | Female WT/WT vs Female Δ HEL/WT | -0.3980 | 0.0059 | TRUE |
| c5 | Male WT/WT vs Male Δ HEL/WT | -0.2236 | 0.2411 | FALSE |
| c6 | Female Δ HEL/WT vs Male Δ HEL/WT | 0.7586 | 0.0000 | TRUE |

##### 3.14.10 relative V fat

|  | test | Difference | p\_value | Significance |
| --- | --- | --- | --- | --- |
| c2 | WT/WT vs Δ HEL/WT | 0.0141 | 1.0000 | FALSE |
| c3 | Female vs Male | 0.6374 | 0.8666 | FALSE |
| c4 | Female WT/WT vs Female Δ HEL/WT | 0.3969 | 0.9844 | FALSE |
| c5 | Male WT/WT vs Male Δ HEL/WT | -0.3686 | 0.9899 | FALSE |
| c6 | Female Δ HEL/WT vs Male Δ HEL/WT | 0.2547 | 0.9966 | FALSE |

##### 3.14.11 relative V lean

|  | test | Difference | p\_value | Significance |
| --- | --- | --- | --- | --- |
| c2 | WT/WT vs Δ HEL/WT | 0.4627 | 0.9370 | FALSE |
| c3 | Female vs Male | -1.4402 | 0.3055 | FALSE |
| c4 | Female WT/WT vs Female Δ HEL/WT | 0.0167 | 1.0000 | FALSE |
| c5 | Male WT/WT vs Male Δ HEL/WT | 0.9087 | 0.8648 | FALSE |
| c6 | Female Δ HEL/WT vs Male Δ HEL/WT | -0.9941 | 0.8312 | FALSE |

##### 3.14.12 Tissue Mineral Density

|  | test | Difference | p\_value | Significance |
| --- | --- | --- | --- | --- |
| c2 | WT/WT vs Δ HEL/WT | -0.0029 | 0.8450 | FALSE |
| c3 | Female vs Male | 0.0240 | 0.0000 | TRUE |
| c4 | Female WT/WT vs Female Δ HEL/WT | 0.0004 | 0.9998 | FALSE |
| c5 | Male WT/WT vs Male Δ HEL/WT | -0.0063 | 0.6385 | FALSE |
| c6 | Female Δ HEL/WT vs Male Δ HEL/WT | 0.0206 | 0.0040 | TRUE |

#### 3.15 Skeleton morfology

**Number of animals**

| STRAIN | GENDER | count |
| --- | --- | --- |
| WT/WT | female | 8 |
| WT/WT | male | 8 |
| Δ HEL/WT | female | 8 |
| Δ HEL/WT | male | 6 |

##### 3.15.1 Tibia length

|  | test | Difference | p\_value | Significance |
| --- | --- | --- | --- | --- |
| c2 | WT/WT vs Δ HEL/WT | 0.0660 | 0.9509 | FALSE |
| c3 | Female vs Male | 0.0517 | 0.9745 | FALSE |
| c4 | Female WT/WT vs Female Δ HEL/WT | 0.0912 | 0.9512 | FALSE |
| c5 | Male WT/WT vs Male Δ HEL/WT | 0.0408 | 0.9959 | FALSE |
| c6 | Female Δ HEL/WT vs Male Δ HEL/WT | 0.0265 | 0.9989 | FALSE |

##### 3.15.2 Body length

|  | test | Difference | p\_value | Significance |
| --- | --- | --- | --- | --- |
| c2 | WT/WT vs Δ HEL/WT | 1.2192 | 0.3420 | FALSE |
| c3 | Female vs Male | -2.5346 | 0.0044 | TRUE |
| c4 | Female WT/WT vs Female Δ HEL/WT | 0.8225 | 0.8369 | FALSE |
| c5 | Male WT/WT vs Male Δ HEL/WT | 1.6158 | 0.4327 | FALSE |
| c6 | Female Δ HEL/WT vs Male Δ HEL/WT | -2.1379 | 0.1958 | FALSE |

##### 3.15.3 Length of hindpaw

|  | test | Difference | p\_value | Significance |
| --- | --- | --- | --- | --- |
| c2 | WT/WT vs Δ HEL/WT | 0.4210 | 0.0679 | FALSE |
| c3 | Female vs Male | -0.3287 | 0.1871 | FALSE |
| c4 | Female WT/WT vs Female Δ HEL/WT | 0.2606 | 0.7023 | FALSE |
| c5 | Male WT/WT vs Male Δ HEL/WT | 0.5814 | 0.0792 | FALSE |
| c6 | Female Δ HEL/WT vs Male Δ HEL/WT | -0.1683 | 0.8957 | FALSE |

##### 3.15.4 Length of cranium

|  | test | Difference | p\_value | Significance |
| --- | --- | --- | --- | --- |
| c2 | WT/WT vs Δ HEL/WT | 0.6776 | 0.0083 | TRUE |
| c3 | Female vs Male | -0.2109 | 0.6962 | FALSE |
| c4 | Female WT/WT vs Female Δ HEL/WT | 1.0806 | 0.0035 | TRUE |
| c5 | Male WT/WT vs Male Δ HEL/WT | 0.2746 | 0.7716 | FALSE |
| c6 | Female Δ HEL/WT vs Male Δ HEL/WT | -0.6139 | 0.1569 | FALSE |

##### 3.15.5 Number of caudal vertebrae

|  | 26:27 | 26:28 | 26:29 | 27:28 | 27:29 | 28:29 |
| --- | --- | --- | --- | --- | --- | --- |
| female WT/WT:female Δ HEL/WT | 1 | 0.9 | 1 | 0.9 | 1 | 0.9 |
| female WT/WT:male WT/WT | 1 | 0.9 | 1 | 0.9 | 1 | 0.9 |
| female WT/WT:male Δ HEL/WT | 1 | 1.0 | 1 | 1.0 | 1 | 1.0 |
| female Δ HEL/WT:male WT/WT | 1 | 1.0 | 1 | 1.0 | 1 | 1.0 |
| female Δ HEL/WT:male Δ HEL/WT | 1 | 0.9 | 1 | 0.9 | 1 | 1.0 |
| male WT/WT:male Δ HEL/WT | 1 | 0.9 | 1 | 0.9 | 1 | 1.0 |

##### 3.15.6 Number of cervical vertebrae

##### 3.15.7 Number of lumbar vertebrae

|  | female WT/WT | female Δ HEL/WT | male WT/WT |
| --- | --- | --- | --- |
| female Δ HEL/WT | 1 | NA | NA |
| male WT/WT | 1 | 1 | NA |
| male Δ HEL/WT | 1 | 1 | 1 |

##### 3.15.8 Number of pelvic vertebrae

|  | 3:4 | 3:5 | 4:5 |
| --- | --- | --- | --- |
| female WT/WT:female Δ HEL/WT | 1 | 1 | 1 |
| female WT/WT:male WT/WT | 1 | 1 | 1 |
| female WT/WT:male Δ HEL/WT | 1 | 1 | 1 |
| female Δ HEL/WT:male WT/WT | 1 | 1 | 1 |
| female Δ HEL/WT:male Δ HEL/WT | 1 | 1 | 1 |
| male WT/WT:male Δ HEL/WT | 1 | 1 | 1 |

##### 3.15.9 Number of thoracic vertebrae

|  | female WT/WT | female Δ HEL/WT | male WT/WT |
| --- | --- | --- | --- |
| female Δ HEL/WT | 0.6000 | NA | NA |
| male WT/WT | 1.0000 | 0.6000 | NA |
| male Δ HEL/WT | 0.6429 | 0.6965 | 0.6429 |

##### 3.15.10 Number of digits

##### 3.15.11 Number of digits in forepaws

##### 3.15.12 Number of digits lo hindpaws

##### 3.15.13 Number of ribs left

|  | female WT/WT | female Δ HEL/WT | male WT/WT |
| --- | --- | --- | --- |
| female Δ HEL/WT | 0.2308 | NA | NA |
| male WT/WT | 1.0000 | 0.2308 | NA |
| male Δ HEL/WT | 0.2473 | 0.7524 | 0.2473 |

##### 3.15.14 Number of ribs right

|  | female WT/WT | female Δ HEL/WT | male WT/WT |
| --- | --- | --- | --- |
| female Δ HEL/WT | 0.6000 | NA | NA |
| male WT/WT | 1.0000 | 0.6000 | NA |
| male Δ HEL/WT | 0.6429 | 0.6965 | 0.6429 |

##### 3.15.15 Brachydactyly

##### 3.15.16 Caudal processes

##### 3.15.17 Cervical processes

##### 3.15.18 Clavicle

##### 3.15.19 Digit integrity

##### 3.15.20 Femur

##### 3.15.21 Fibula

##### 3.15.22 Fusion of processes

##### 3.15.23 Fusion of ribs

##### 3.15.24 Fusion of vertebrae

##### 3.15.25 Humerus

##### 3.15.26 Joints

##### 3.15.27 Kyphosis

##### 3.15.28 Lordosis

##### 3.15.29 Lumbar processes

|  | female WT/WT | female Δ HEL/WT | male WT/WT |
| --- | --- | --- | --- |
| female Δ HEL/WT | 0.2378 | NA | NA |
| male WT/WT | 1.0000 | 0.1538 | NA |
| male Δ HEL/WT | 0.3671 | 1.0000 | 0.1648 |

##### 3.15.30 Mandibles

##### 3.15.31 Maxilla/Pre-maxilla

##### 3.15.32 Missing cranial rib

##### 3.15.33 Pelvis

##### 3.15.34 Polysyndactylism

##### 3.15.35 Presence of baculum bone

|  | female WT/WT | female Δ HEL/WT | male WT/WT |
| --- | --- | --- | --- |
| female Δ HEL/WT | 1e+00 | NA | NA |
| male WT/WT | 9e-04 | 0.0028 | NA |
| male Δ HEL/WT | 1e-03 | 0.0070 | 1 |

##### 3.15.36 Processes on vertebrae

|  | female WT/WT | female Δ HEL/WT | male WT/WT |
| --- | --- | --- | --- |
| female Δ HEL/WT | 0.0811 | NA | NA |
| male WT/WT | 1.0000 | 0.042 | NA |
| male Δ HEL/WT | 0.1364 | 1.000 | 0.045 |

##### 3.15.37 Quality of enamel on incisors

##### 3.15.38 Quality of enamel on molars

##### 3.15.39 Radius

##### 3.15.40 Sacral processes

|  | female WT/WT | female Δ HEL/WT | male WT/WT |
| --- | --- | --- | --- |
| female Δ HEL/WT | 1 | NA | NA |
| male WT/WT | 1 | 1 | NA |
| male Δ HEL/WT | 1 | 1 | 1 |

##### 3.15.41 Scapulae

##### 3.15.42 Scoliosis

|  | female WT/WT | female Δ HEL/WT | male WT/WT |
| --- | --- | --- | --- |
| female Δ HEL/WT | 1 | NA | NA |
| male WT/WT | 1 | 1 | NA |
| male Δ HEL/WT | 1 | 1 | 1 |

##### 3.15.43 Shape of ribcage

|  | female WT/WT | female Δ HEL/WT | male WT/WT |
| --- | --- | --- | --- |
| female Δ HEL/WT | 1 | NA | NA |
| male WT/WT | 1 | 1 | NA |
| male Δ HEL/WT | 1 | 1 | 1 |

##### 3.15.44 Shape of ribs

|  | female WT/WT | female Δ HEL/WT | male WT/WT |
| --- | --- | --- | --- |
| female Δ HEL/WT | 1 | NA | NA |
| male WT/WT | 1 | 1 | NA |
| male Δ HEL/WT | 1 | 1 | 1 |

##### 3.15.45 Shape of spine

|  | female WT/WT | female Δ HEL/WT | male WT/WT |
| --- | --- | --- | --- |
| female Δ HEL/WT | 1 | NA | NA |
| male WT/WT | 1 | 1 | NA |
| male Δ HEL/WT | 1 | 1 | 1 |

##### 3.15.46 Shape of vertebrae

|  | female WT/WT | female Δ HEL/WT | male WT/WT |
| --- | --- | --- | --- |
| female Δ HEL/WT | 1 | NA | NA |
| male WT/WT | 1 | 1 | NA |
| male Δ HEL/WT | 1 | 1 | 1 |

##### 3.15.47 Skull shape

##### 3.15.48 Syndactylism

##### 3.15.49 Teeth

|  | female WT/WT | female Δ HEL/WT | male WT/WT |
| --- | --- | --- | --- |
| female Δ HEL/WT | 1 | NA | NA |
| male WT/WT | 1 | 1 | NA |
| male Δ HEL/WT | 1 | 1 | 1 |

##### 3.15.50 Thoracic processes

|  | female WT/WT | female Δ HEL/WT | male WT/WT |
| --- | --- | --- | --- |
| female Δ HEL/WT | 0.7 | NA | NA |
| male WT/WT | 1.0 | 0.7 | NA |
| male Δ HEL/WT | 0.7 | 1.0 | 0.7 |

##### 3.15.51 Tibia

##### 3.15.52 Transitional vertebrae

|  | female WT/WT | female Δ HEL/WT | male WT/WT |
| --- | --- | --- | --- |
| female Δ HEL/WT | 0.8706 | NA | NA |
| male WT/WT | 1.0000 | 0.8706 | NA |
| male Δ HEL/WT | 1.0000 | 0.8706 | 0.8706 |

##### 3.15.53 Ulna

##### 28Y-25534 Dorso Ventral Dorso Ventral Lateral Orientation Forepaw Skull Lateral Orientation Skull Dorso Ventral Orientation Hind Leg and Hip Hind Leg and Hip uCT image: detail 1 28Y-25536 Dorso Ventral Dorso Ventral Lateral Orientation Forepaw Skull Lateral Orientation Skull Dorso Ventral Orientation Hind Leg and Hip Hind Leg and Hip uCT image: detail 2 uCT image: detail 1 28Y-25537 Dorso Ventral Dorso Ventral Lateral Orientation Forepaw Skull Lateral Orientation Skull Dorso Ventral Orientation Hind Leg and Hip Hind Leg and Hip uCT image: detail 1 28Y-25576 Dorso Ventral Dorso Ventral Lateral Orientation Forepaw Skull Lateral Orientation Skull Dorso Ventral Orientation Hind Leg and Hip Hind Leg and Hip uCT image: detail 1 28Y-25577 Dorso Ventral Dorso Ventral Lateral Orientation Forepaw Skull Lateral Orientation Skull Dorso Ventral Orientation Hind Leg and Hip Hind Leg and Hip uCT image: detail 1 28Y-25579 Dorso Ventral Dorso Ventral Lateral Orientation Forepaw Skull Lateral Orientation Skull Dorso Ventral Orientation Hind Leg and Hip Hind Leg and Hip 28Y-25580 Dorso Ventral Dorso Ventral Lateral Orientation Forepaw Skull Lateral Orientation Skull Dorso Ventral Orientation Hind Leg and Hip Hind Leg and Hip uCT image: detail 2 uCT image: detail 1 28Y-25588 Dorso Ventral Dorso Ventral Lateral Orientation Forepaw Skull Lateral Orientation Skull Dorso Ventral Orientation Hind Leg and Hip Hind Leg and Hip uCT image: detail 2 uCT image: detail 1 28Y-25589 Dorso Ventral Dorso Ventral Lateral Orientation Forepaw Skull Lateral Orientation Skull Dorso Ventral Orientation Hind Leg and Hip Hind Leg and Hip uCT image: detail 1 28Y-25590 Dorso Ventral Dorso Ventral Lateral Orientation Forepaw Skull Lateral Orientation Skull Dorso Ventral Orientation Hind Leg and Hip Hind Leg and Hip uCT image: detail 1 28Y-25592 Dorso Ventral Dorso Ventral Lateral Orientation Forepaw Skull Lateral Orientation Skull Dorso Ventral Orientation Hind Leg and Hip Hind Leg and Hip uCT image: detail 1

#### 3.16 Vision screen

**Number of animals**

| STRAIN | GENDER | count |
| --- | --- | --- |
| WT/WT | female | 8 |
| WT/WT | male | 8 |
| Δ HEL/WT | female | 7 |
| Δ HEL/WT | male | 6 |

##### 3.16.1 Corneal opacity

|  | female WT/WT | female Δ HEL/WT | male WT/WT |
| --- | --- | --- | --- |
| female Δ HEL/WT | 0.1217 | NA | NA |
| male WT/WT | 1.0000 | 0.2378 | NA |
| male Δ HEL/WT | 0.5670 | 0.0280 | 0.3132 |

##### 3.16.2 Fusion between cornea and lens

##### 3.16.3 Lens Opacity

|  | female WT/WT | female Δ HEL/WT | male WT/WT |
| --- | --- | --- | --- |
| female Δ HEL/WT | 1 | NA | NA |
| male WT/WT | 1 | 1 | NA |
| male Δ HEL/WT | 1 | 1 | 1 |

##### 3.16.4 Fusion between cornea and lens

##### 3.16.5 Lens

|  | female WT/WT | female Δ HEL/WT | male WT/WT |
| --- | --- | --- | --- |
| female Δ HEL/WT | 0.5600 | NA | NA |
| male WT/WT | 0.2308 | 0.4231 | NA |
| male Δ HEL/WT | 0.2308 | 0.4231 | 1 |

##### 3.16.6 Optic Disc

##### 3.16.7 Retina Complete

##### 3.16.8 Retinal Blood Vessels

##### 3.16.9 Retinal Blood Vessels Pattern

##### 3.16.10 Retinal Blood Vessels Structure

#### 3.17 Hematology

**Number of animals**

| STRAIN | GENDER | count |
| --- | --- | --- |
| WT/WT | female | 8 |
| WT/WT | male | 8 |
| Δ HEL/WT | female | 7 |
| Δ HEL/WT | male | 6 |

##### 3.17.1 Eosinophil cell count

|  | test | Difference | p\_value | Significance |
| --- | --- | --- | --- | --- |
| c2 | WT/WT vs Δ HEL/WT | 0.0074 | 0.9462 | FALSE |
| c3 | Female vs Male | -0.0110 | 0.9442 | FALSE |
| c4 | Female WT/WT vs Female Δ HEL/WT | -0.0034 | 0.9979 | FALSE |
| c5 | Male WT/WT vs Male Δ HEL/WT | 0.0182 | 0.7755 | FALSE |
| c6 | Female Δ HEL/WT vs Male Δ HEL/WT | -0.0002 | 1.0000 | FALSE |

##### 3.17.2 Eosinophil differential count

|  | test | Difference | p\_value | Significance |
| --- | --- | --- | --- | --- |
| c2 | WT/WT vs Δ HEL/WT | -0.0585 | 0.9983 | FALSE |
| c3 | Female vs Male | -0.1592 | 0.9899 | FALSE |
| c4 | Female WT/WT vs Female Δ HEL/WT | -0.4482 | 0.7978 | FALSE |
| c5 | Male WT/WT vs Male Δ HEL/WT | 0.3312 | 0.9075 | FALSE |
| c6 | Female Δ HEL/WT vs Male Δ HEL/WT | 0.2306 | 0.9870 | FALSE |

##### 3.17.3 Hematocrit

|  | test | Difference | p\_value | Significance |
| --- | --- | --- | --- | --- |
| c2 | WT/WT vs Δ HEL/WT | 0.3077 | 0.9788 | FALSE |
| c3 | Female vs Male | 1.5256 | 0.2171 | FALSE |
| c4 | Female WT/WT vs Female Δ HEL/WT | 0.3071 | 0.9918 | FALSE |
| c5 | Male WT/WT vs Male Δ HEL/WT | 0.3083 | 0.9927 | FALSE |
| c6 | Female Δ HEL/WT vs Male Δ HEL/WT | 1.5262 | 0.5572 | FALSE |

##### 3.17.4 Hemoglobin

|  | test | Difference | p\_value | Significance |
| --- | --- | --- | --- | --- |
| c2 | WT/WT vs Δ HEL/WT | 0.1355 | 0.9378 | FALSE |
| c3 | Female vs Male | 0.5689 | 0.0971 | FALSE |
| c4 | Female WT/WT vs Female Δ HEL/WT | 0.1036 | 0.9885 | FALSE |
| c5 | Male WT/WT vs Male Δ HEL/WT | 0.1674 | 0.9594 | FALSE |
| c6 | Female Δ HEL/WT vs Male Δ HEL/WT | 0.6008 | 0.3377 | FALSE |

##### 3.17.5 Lymphocyte cell count

|  | test | Difference | p\_value | Significance |
| --- | --- | --- | --- | --- |
| c2 | WT/WT vs Δ HEL/WT | 0.5021 | 0.6928 | FALSE |
| c3 | Female vs Male | -0.4250 | 0.7895 | FALSE |
| c4 | Female WT/WT vs Female Δ HEL/WT | 1.0409 | 0.3640 | FALSE |
| c5 | Male WT/WT vs Male Δ HEL/WT | -0.0367 | 0.9999 | FALSE |
| c6 | Female Δ HEL/WT vs Male Δ HEL/WT | -0.9638 | 0.4957 | FALSE |

##### 3.17.6 Lymphocyte differential count

|  | test | Difference | p\_value | Significance |
| --- | --- | --- | --- | --- |
| c2 | WT/WT vs Δ HEL/WT | -4.1440 | 0.0736 | FALSE |
| c3 | Female vs Male | 1.8528 | 0.7211 | FALSE |
| c4 | Female WT/WT vs Female Δ HEL/WT | 1.4054 | 0.9291 | FALSE |
| c5 | Male WT/WT vs Male Δ HEL/WT | -9.6934 | 0.0005 | TRUE |
| c6 | Female Δ HEL/WT vs Male Δ HEL/WT | -3.6966 | 0.4826 | FALSE |

##### 3.17.7 Mean cell hemoglobin concentration

|  | test | Difference | p\_value | Significance |
| --- | --- | --- | --- | --- |
| c2 | WT/WT vs Δ HEL/WT | 0.1003 | 0.9266 | FALSE |
| c3 | Female vs Male | 0.2164 | 0.5502 | FALSE |
| c4 | Female WT/WT vs Female Δ HEL/WT | 0.0214 | 0.9997 | FALSE |
| c5 | Male WT/WT vs Male Δ HEL/WT | 0.1792 | 0.8712 | FALSE |
| c6 | Female Δ HEL/WT vs Male Δ HEL/WT | 0.2952 | 0.6186 | FALSE |

##### 3.17.8 Mean cell volume

|  | test | Difference | p\_value | Significance |
| --- | --- | --- | --- | --- |
| c2 | WT/WT vs Δ HEL/WT | 0.4420 | 0.5811 | FALSE |
| c3 | Female vs Male | 0.7205 | 0.1715 | FALSE |
| c4 | Female WT/WT vs Female Δ HEL/WT | 0.3839 | 0.8528 | FALSE |
| c5 | Male WT/WT vs Male Δ HEL/WT | 0.5000 | 0.7492 | FALSE |
| c6 | Female Δ HEL/WT vs Male Δ HEL/WT | 0.7786 | 0.4371 | FALSE |

##### 3.17.9 Mean corpuscular hemoglobin

|  | test | Difference | p\_value | Significance |
| --- | --- | --- | --- | --- |
| c2 | WT/WT vs Δ HEL/WT | 0.1970 | 0.3485 | FALSE |
| c3 | Female vs Male | 0.3113 | 0.0497 | TRUE |
| c4 | Female WT/WT vs Female Δ HEL/WT | 0.1607 | 0.7578 | FALSE |
| c5 | Male WT/WT vs Male Δ HEL/WT | 0.2333 | 0.5227 | FALSE |
| c6 | Female Δ HEL/WT vs Male Δ HEL/WT | 0.3476 | 0.2067 | FALSE |

##### 3.17.10 Mean platelet volume

|  | test | Difference | p\_value | Significance |
| --- | --- | --- | --- | --- |
| c2 | WT/WT vs Δ HEL/WT | -0.0581 | 0.7772 | FALSE |
| c3 | Female vs Male | 0.1293 | 0.2916 | FALSE |
| c4 | Female WT/WT vs Female Δ HEL/WT | -0.0807 | 0.7768 | FALSE |
| c5 | Male WT/WT vs Male Δ HEL/WT | -0.0354 | 0.9768 | FALSE |
| c6 | Female Δ HEL/WT vs Male Δ HEL/WT | 0.1520 | 0.4470 | FALSE |

##### 3.17.11 Monocyte cell count

|  | test | Difference | p\_value | Significance |
| --- | --- | --- | --- | --- |
| c2 | WT/WT vs Δ HEL/WT | 0.0617 | 0.2470 | FALSE |
| c3 | Female vs Male | -0.0692 | 0.1614 | FALSE |
| c4 | Female WT/WT vs Female Δ HEL/WT | 0.0296 | 0.9134 | FALSE |
| c5 | Male WT/WT vs Male Δ HEL/WT | 0.0938 | 0.2070 | FALSE |
| c6 | Female Δ HEL/WT vs Male Δ HEL/WT | -0.0371 | 0.8705 | FALSE |

##### 3.17.12 Monocyte differential count

|  | test | Difference | p\_value | Significance |
| --- | --- | --- | --- | --- |
| c2 | WT/WT vs Δ HEL/WT | 0.2521 | 0.9387 | FALSE |
| c3 | Female vs Male | -0.5979 | 0.5262 | FALSE |
| c4 | Female WT/WT vs Female Δ HEL/WT | -0.4125 | 0.9036 | FALSE |
| c5 | Male WT/WT vs Male Δ HEL/WT | 0.9167 | 0.4747 | FALSE |
| c6 | Female Δ HEL/WT vs Male Δ HEL/WT | 0.0667 | 0.9996 | FALSE |

##### 3.17.13 Neutrophil cell count

|  | test | Difference | p\_value | Significance |
| --- | --- | --- | --- | --- |
| c2 | WT/WT vs Δ HEL/WT | 0.3410 | 0.0009 | TRUE |
| c3 | Female vs Male | -0.1514 | 0.3303 | FALSE |
| c4 | Female WT/WT vs Female Δ HEL/WT | 0.1398 | 0.6669 | FALSE |
| c5 | Male WT/WT vs Male Δ HEL/WT | 0.5421 | 0.0002 | TRUE |
| c6 | Female Δ HEL/WT vs Male Δ HEL/WT | 0.0498 | 0.9814 | FALSE |

##### 3.17.14 Neutrophil differential count

|  | test | Difference | p\_value | Significance |
| --- | --- | --- | --- | --- |
| c2 | WT/WT vs Δ HEL/WT | 3.7905 | 0.0640 | FALSE |
| c3 | Female vs Male | -1.0488 | 0.8954 | FALSE |
| c4 | Female WT/WT vs Female Δ HEL/WT | -0.7607 | 0.9827 | FALSE |
| c5 | Male WT/WT vs Male Δ HEL/WT | 8.3417 | 0.0008 | TRUE |
| c6 | Female Δ HEL/WT vs Male Δ HEL/WT | 3.5024 | 0.4005 | FALSE |

##### 3.17.15 Platelet count

|  | test | Difference | p\_value | Significance |
| --- | --- | --- | --- | --- |
| c2 | WT/WT vs Δ HEL/WT | -2.1310 | 0.9998 | FALSE |
| c3 | Female vs Male | -78.0774 | 0.0338 | TRUE |
| c4 | Female WT/WT vs Female Δ HEL/WT | -3.3036 | 0.9998 | FALSE |
| c5 | Male WT/WT vs Male Δ HEL/WT | -0.9583 | 1.0000 | FALSE |
| c6 | Female Δ HEL/WT vs Male Δ HEL/WT | -76.9048 | 0.2617 | FALSE |

##### 3.17.16 Platelet crit

|  | test | Difference | p\_value | Significance |
| --- | --- | --- | --- | --- |
| c2 | WT/WT vs Δ HEL/WT | -0.0455 | 0.9858 | FALSE |
| c3 | Female vs Male | -0.2653 | 0.1993 | FALSE |
| c4 | Female WT/WT vs Female Δ HEL/WT | -0.0539 | 0.9910 | FALSE |
| c5 | Male WT/WT vs Male Δ HEL/WT | -0.0371 | 0.9974 | FALSE |
| c6 | Female Δ HEL/WT vs Male Δ HEL/WT | -0.2569 | 0.5646 | FALSE |

##### 3.17.17 Platelet distribution width

|  | test | Difference | p\_value | Significance |
| --- | --- | --- | --- | --- |
| c2 | WT/WT vs Δ HEL/WT | -0.0295 | 0.9707 | FALSE |
| c3 | Female vs Male | 0.1420 | 0.1559 | FALSE |
| c4 | Female WT/WT vs Female Δ HEL/WT | -0.0089 | 0.9997 | FALSE |
| c5 | Male WT/WT vs Male Δ HEL/WT | -0.0500 | 0.9538 | FALSE |
| c6 | Female Δ HEL/WT vs Male Δ HEL/WT | 0.1214 | 0.6138 | FALSE |

##### 3.17.18 Red blood cell count

|  | test | Difference | p\_value | Significance |
| --- | --- | --- | --- | --- |
| c2 | WT/WT vs Δ HEL/WT | -0.0189 | 0.9993 | FALSE |
| c3 | Female vs Male | 0.1714 | 0.6647 | FALSE |
| c4 | Female WT/WT vs Female Δ HEL/WT | -0.0154 | 0.9999 | FALSE |
| c5 | Male WT/WT vs Male Δ HEL/WT | -0.0225 | 0.9996 | FALSE |
| c6 | Female Δ HEL/WT vs Male Δ HEL/WT | 0.1679 | 0.8749 | FALSE |

##### 3.17.19 Red blood cell distribution width

|  | test | Difference | p\_value | Significance |
| --- | --- | --- | --- | --- |
| c2 | WT/WT vs Δ HEL/WT | -0.5701 | 0.0320 | TRUE |
| c3 | Female vs Male | 0.1930 | 0.9189 | FALSE |
| c4 | Female WT/WT vs Female Δ HEL/WT | -0.3155 | 0.6805 | FALSE |
| c5 | Male WT/WT vs Male Δ HEL/WT | -0.8247 | 0.0265 | TRUE |
| c6 | Female Δ HEL/WT vs Male Δ HEL/WT | -0.0616 | 0.9987 | FALSE |

##### 3.17.20 White blood cell count

|  | test | Difference | p\_value | Significance |
| --- | --- | --- | --- | --- |
| c2 | WT/WT vs Δ HEL/WT | 0.9212 | 0.3082 | FALSE |
| c3 | Female vs Male | -0.6608 | 0.5944 | FALSE |
| c4 | Female WT/WT vs Female Δ HEL/WT | 1.2195 | 0.3469 | FALSE |
| c5 | Male WT/WT vs Male Δ HEL/WT | 0.6229 | 0.8443 | FALSE |
| c6 | Female Δ HEL/WT vs Male Δ HEL/WT | -0.9590 | 0.6135 | FALSE |

#### 3.18 Biochemistry

**Number of animals**

| STRAIN | GENDER | count |
| --- | --- | --- |
| WT/WT | female | 8 |
| WT/WT | male | 8 |
| Δ HEL/WT | female | 7 |
| Δ HEL/WT | male | 6 |

##### 3.18.1 Alanine aminotransferase

|  | test | Difference | p\_value | Significance |
| --- | --- | --- | --- | --- |
| c2 | WT/WT vs Δ HEL/WT | -4.6461 | 0.8817 | FALSE |
| c3 | Female vs Male | -22.1320 | 0.1035 | FALSE |
| c4 | Female WT/WT vs Female Δ HEL/WT | 1.0388 | 0.9994 | FALSE |
| c5 | Male WT/WT vs Male Δ HEL/WT | -10.3310 | 0.6559 | FALSE |
| c6 | Female Δ HEL/WT vs Male Δ HEL/WT | -27.8168 | 0.1333 | FALSE |

##### 3.18.2 Albumin

|  | test | Difference | p\_value | Significance |
| --- | --- | --- | --- | --- |
| c2 | WT/WT vs Δ HEL/WT | -0.8243 | 0.1999 | FALSE |
| c3 | Female vs Male | 1.5684 | 0.0044 | TRUE |
| c4 | Female WT/WT vs Female Δ HEL/WT | -0.4333 | 0.8716 | FALSE |
| c5 | Male WT/WT vs Male Δ HEL/WT | -1.2154 | 0.1794 | FALSE |
| c6 | Female Δ HEL/WT vs Male Δ HEL/WT | 1.1773 | 0.2766 | FALSE |

##### 3.18.3 Alkaline phosphatase

|  | test | Difference | p\_value | Significance |
| --- | --- | --- | --- | --- |
| c2 | WT/WT vs Δ HEL/WT | 10.7298 | 0.1809 | FALSE |
| c3 | Female vs Male | 3.1120 | 0.9844 | FALSE |
| c4 | Female WT/WT vs Female Δ HEL/WT | 22.1661 | 0.0198 | TRUE |
| c5 | Male WT/WT vs Male Δ HEL/WT | -0.7064 | 0.9997 | FALSE |
| c6 | Female Δ HEL/WT vs Male Δ HEL/WT | -8.3242 | 0.8839 | FALSE |

##### 3.18.4 Aspartate aminotransferase

|  | test | Difference | p\_value | Significance |
| --- | --- | --- | --- | --- |
| c2 | WT/WT vs Δ HEL/WT | -1.7798 | 0.9688 | FALSE |
| c3 | Female vs Male | -0.6369 | 0.9985 | FALSE |
| c4 | Female WT/WT vs Female Δ HEL/WT | -0.8929 | 0.9984 | FALSE |
| c5 | Male WT/WT vs Male Δ HEL/WT | -2.6667 | 0.9654 | FALSE |
| c6 | Female Δ HEL/WT vs Male Δ HEL/WT | -1.5238 | 0.9937 | FALSE |

##### 3.18.5 Calcium

|  | test | Difference | p\_value | Significance |
| --- | --- | --- | --- | --- |
| c2 | WT/WT vs Δ HEL/WT | -0.0088 | 0.9993 | FALSE |
| c3 | Female vs Male | 0.0955 | 0.5249 | FALSE |
| c4 | Female WT/WT vs Female Δ HEL/WT | -0.0343 | 0.9844 | FALSE |
| c5 | Male WT/WT vs Male Δ HEL/WT | 0.0167 | 0.9984 | FALSE |
| c6 | Female Δ HEL/WT vs Male Δ HEL/WT | 0.1210 | 0.6514 | FALSE |

##### 3.18.6 Chloride

|  | test | Difference | p\_value | Significance |
| --- | --- | --- | --- | --- |
| c2 | WT/WT vs Δ HEL/WT | -0.5636 | 0.6617 | FALSE |
| c3 | Female vs Male | 1.5770 | 0.0340 | TRUE |
| c4 | Female WT/WT vs Female Δ HEL/WT | -0.2912 | 0.9733 | FALSE |
| c5 | Male WT/WT vs Male Δ HEL/WT | -0.8361 | 0.6344 | FALSE |
| c6 | Female Δ HEL/WT vs Male Δ HEL/WT | 1.3045 | 0.3685 | FALSE |

##### 3.18.7 Creatinine

|  | test | Difference | p\_value | Significance |
| --- | --- | --- | --- | --- |
| c2 | WT/WT vs Δ HEL/WT | 0.0003 | 0.9998 | FALSE |
| c3 | Female vs Male | 0.0188 | 0.0000 | TRUE |
| c4 | Female WT/WT vs Female Δ HEL/WT | 0.0009 | 0.9978 | FALSE |
| c5 | Male WT/WT vs Male Δ HEL/WT | -0.0003 | 0.9999 | FALSE |
| c6 | Female Δ HEL/WT vs Male Δ HEL/WT | 0.0182 | 0.0047 | TRUE |

##### 3.18.8 Glucose

|  | test | Difference | p\_value | Significance |
| --- | --- | --- | --- | --- |
| c2 | WT/WT vs Δ HEL/WT | -3.5371 | 0.9544 | FALSE |
| c3 | Female vs Male | -23.8878 | 0.0038 | TRUE |
| c4 | Female WT/WT vs Female Δ HEL/WT | -2.5772 | 0.9928 | FALSE |
| c5 | Male WT/WT vs Male Δ HEL/WT | -4.4970 | 0.9682 | FALSE |
| c6 | Female Δ HEL/WT vs Male Δ HEL/WT | -24.8477 | 0.0801 | FALSE |

##### 3.18.9 HDL-cholesterol

|  | test | Difference | p\_value | Significance |
| --- | --- | --- | --- | --- |
| c2 | WT/WT vs Δ HEL/WT | 4.7982 | 0.4993 | FALSE |
| c3 | Female vs Male | -27.8113 | 0.0000 | TRUE |
| c4 | Female WT/WT vs Female Δ HEL/WT | 7.2660 | 0.4307 | FALSE |
| c5 | Male WT/WT vs Male Δ HEL/WT | 2.3304 | 0.9616 | FALSE |
| c6 | Female Δ HEL/WT vs Male Δ HEL/WT | -30.2791 | 0.0000 | TRUE |

##### 3.18.10 Iron

|  | test | Difference | p\_value | Significance |
| --- | --- | --- | --- | --- |
| c2 | WT/WT vs Δ HEL/WT | -0.0223 | 0.0905 | FALSE |
| c3 | Female vs Male | 0.0126 | 0.8192 | FALSE |
| c4 | Female WT/WT vs Female Δ HEL/WT | -0.0317 | 0.0839 | FALSE |
| c5 | Male WT/WT vs Male Δ HEL/WT | -0.0128 | 0.7598 | FALSE |
| c6 | Female Δ HEL/WT vs Male Δ HEL/WT | 0.0221 | 0.6548 | FALSE |

##### 3.18.11 Phosphate

|  | test | Difference | p\_value | Significance |
| --- | --- | --- | --- | --- |
| c2 | WT/WT vs Δ HEL/WT | 0.1518 | 0.9303 | FALSE |
| c3 | Female vs Male | 1.0721 | 0.0814 | FALSE |
| c4 | Female WT/WT vs Female Δ HEL/WT | 0.0308 | 0.9998 | FALSE |
| c5 | Male WT/WT vs Male Δ HEL/WT | 0.2727 | 0.8678 | FALSE |
| c6 | Female Δ HEL/WT vs Male Δ HEL/WT | 1.1931 | 0.1659 | FALSE |

##### 3.18.12 Potassium

|  | test | Difference | p\_value | Significance |
| --- | --- | --- | --- | --- |
| c2 | WT/WT vs Δ HEL/WT | 0.4311 | 0.4775 | FALSE |
| c3 | Female vs Male | -1.4582 | 0.0000 | TRUE |
| c4 | Female WT/WT vs Female Δ HEL/WT | 0.1205 | 0.9912 | FALSE |
| c5 | Male WT/WT vs Male Δ HEL/WT | 0.7417 | 0.3228 | FALSE |
| c6 | Female Δ HEL/WT vs Male Δ HEL/WT | -1.1476 | 0.0566 | FALSE |

##### 3.18.13 Sodium

|  | test | Difference | p\_value | Significance |
| --- | --- | --- | --- | --- |
| c2 | WT/WT vs Δ HEL/WT | -1.1190 | 0.2090 | FALSE |
| c3 | Female vs Male | 0.2024 | 0.9841 | FALSE |
| c4 | Female WT/WT vs Female Δ HEL/WT | -0.8214 | 0.7213 | FALSE |
| c5 | Male WT/WT vs Male Δ HEL/WT | -1.4167 | 0.3169 | FALSE |
| c6 | Female Δ HEL/WT vs Male Δ HEL/WT | -0.0952 | 0.9995 | FALSE |

##### 3.18.14 Total bilirubin

|  | test | Difference | p\_value | Significance |
| --- | --- | --- | --- | --- |
| c2 | WT/WT vs Δ HEL/WT | -0.0092 | 0.8957 | FALSE |
| c3 | Female vs Male | -0.0467 | 0.0596 | FALSE |
| c4 | Female WT/WT vs Female Δ HEL/WT | -0.0098 | 0.9496 | FALSE |
| c5 | Male WT/WT vs Male Δ HEL/WT | -0.0085 | 0.9666 | FALSE |
| c6 | Female Δ HEL/WT vs Male Δ HEL/WT | -0.0461 | 0.2465 | FALSE |

##### 3.18.15 Total cholesterol

|  | test | Difference | p\_value | Significance |
| --- | --- | --- | --- | --- |
| c2 | WT/WT vs Δ HEL/WT | 6.0794 | 0.6482 | FALSE |
| c3 | Female vs Male | -35.7339 | 0.0001 | TRUE |
| c4 | Female WT/WT vs Female Δ HEL/WT | 10.9660 | 0.4441 | FALSE |
| c5 | Male WT/WT vs Male Δ HEL/WT | 1.1928 | 0.9984 | FALSE |
| c6 | Female Δ HEL/WT vs Male Δ HEL/WT | -40.6205 | 0.0013 | TRUE |

##### 3.18.16 Total protein

|  | test | Difference | p\_value | Significance |
| --- | --- | --- | --- | --- |
| c2 | WT/WT vs Δ HEL/WT | -1.0705 | 0.3106 | FALSE |
| c3 | Female vs Male | 0.8678 | 0.7898 | FALSE |
| c4 | Female WT/WT vs Female Δ HEL/WT | -1.0104 | 0.6430 | FALSE |
| c5 | Male WT/WT vs Male Δ HEL/WT | -1.1305 | 0.5604 | FALSE |
| c6 | Female Δ HEL/WT vs Male Δ HEL/WT | 0.8077 | 0.9128 | FALSE |

##### 3.18.17 Triglycerides

|  | test | Difference | p\_value | Significance |
| --- | --- | --- | --- | --- |
| c2 | WT/WT vs Δ HEL/WT | 9.3123 | 0.8387 | FALSE |
| c3 | Female vs Male | -19.3012 | 0.6810 | FALSE |
| c4 | Female WT/WT vs Female Δ HEL/WT | 14.9870 | 0.7763 | FALSE |
| c5 | Male WT/WT vs Male Δ HEL/WT | 3.6376 | 0.9956 | FALSE |
| c6 | Female Δ HEL/WT vs Male Δ HEL/WT | -24.9758 | 0.6931 | FALSE |

##### 3.18.18 Urea (Blood Urea Nitrogen - BUN)

|  | test | Difference | p\_value | Significance |
| --- | --- | --- | --- | --- |
| c2 | WT/WT vs Δ HEL/WT | 1.7526 | 0.4573 | FALSE |
| c3 | Female vs Male | -10.2962 | 0.0000 | TRUE |
| c4 | Female WT/WT vs Female Δ HEL/WT | 1.5923 | 0.7751 | FALSE |
| c5 | Male WT/WT vs Male Δ HEL/WT | 1.9130 | 0.6611 | FALSE |
| c6 | Female Δ HEL/WT vs Male Δ HEL/WT | -10.1359 | 0.0012 | TRUE |

#### 3.19 Organ weight

**Number of animals**

| STRAIN | GENDER | count |
| --- | --- | --- |
| WT/WT | female | 8 |
| WT/WT | male | 8 |
| Δ HEL/WT | female | 7 |
| Δ HEL/WT | male | 6 |

##### 3.19.1 Heart weight

|  | test | Difference | p\_value | Significance |
| --- | --- | --- | --- | --- |
| c2 | WT/WT vs Δ HEL/WT | 15.5833 | 0.2437 | FALSE |
| c3 | Female vs Male | -54.1667 | 0.0000 | TRUE |
| c4 | Female WT/WT vs Female Δ HEL/WT | 24.2500 | 0.1570 | FALSE |
| c5 | Male WT/WT vs Male Δ HEL/WT | 6.9167 | 0.9370 | FALSE |
| c6 | Female Δ HEL/WT vs Male Δ HEL/WT | -62.8333 | 0.0000 | TRUE |

##### 3.19.2 Heart weight normalised against body weight

|  | test | Difference | p\_value | Significance |
| --- | --- | --- | --- | --- |
| c2 | WT/WT vs Δ HEL/WT | -2e-04 | 0.8668 | FALSE |
| c3 | Female vs Male | -6e-04 | 0.0841 | FALSE |
| c4 | Female WT/WT vs Female Δ HEL/WT | 1e-04 | 0.9921 | FALSE |
| c5 | Male WT/WT vs Male Δ HEL/WT | -5e-04 | 0.5438 | FALSE |
| c6 | Female Δ HEL/WT vs Male Δ HEL/WT | -9e-04 | 0.0867 | FALSE |

##### 3.19.3 Tibia length

|  | test | Difference | p\_value | Significance |
| --- | --- | --- | --- | --- |
| c2 | WT/WT vs Δ HEL/WT | 0.8542 | 0.1772 | FALSE |
| c3 | Female vs Male | -0.3958 | 0.7739 | FALSE |
| c4 | Female WT/WT vs Female Δ HEL/WT | 0.6250 | 0.6957 | FALSE |
| c5 | Male WT/WT vs Male Δ HEL/WT | 1.0833 | 0.2782 | FALSE |
| c6 | Female Δ HEL/WT vs Male Δ HEL/WT | -0.1667 | 0.9929 | FALSE |

##### 3.19.4 Heart weight normalised against tibia length

|  | test | Difference | p\_value | Significance |
| --- | --- | --- | --- | --- |
| c2 | WT/WT vs Δ HEL/WT | 0.3676 | 0.7997 | FALSE |
| c3 | Female vs Male | -2.3860 | 0.0000 | TRUE |
| c4 | Female WT/WT vs Female Δ HEL/WT | 0.8863 | 0.3962 | FALSE |
| c5 | Male WT/WT vs Male Δ HEL/WT | -0.1512 | 0.9938 | FALSE |
| c6 | Female Δ HEL/WT vs Male Δ HEL/WT | -2.9048 | 0.0000 | TRUE |

##### 3.19.5 Body weight

|  | test | Difference | p\_value | Significance |
| --- | --- | --- | --- | --- |
| c2 | WT/WT vs Δ HEL/WT | 3.7634 | 0.0000 | TRUE |
| c3 | Female vs Male | -5.9259 | 0.0000 | TRUE |
| c4 | Female WT/WT vs Female Δ HEL/WT | 3.5518 | 0.0060 | TRUE |
| c5 | Male WT/WT vs Male Δ HEL/WT | 3.9750 | 0.0028 | TRUE |
| c6 | Female Δ HEL/WT vs Male Δ HEL/WT | -5.7143 | 0.0000 | TRUE |

##### 3.19.6 Left kidney weight

|  | test | Difference | p\_value | Significance |
| --- | --- | --- | --- | --- |
| c2 | WT/WT vs Δ HEL/WT | 0.0460 | 0e+00 | TRUE |
| c3 | Female vs Male | -0.1170 | 0e+00 | TRUE |
| c4 | Female WT/WT vs Female Δ HEL/WT | 0.0397 | 3e-04 | TRUE |
| c5 | Male WT/WT vs Male Δ HEL/WT | 0.0524 | 0e+00 | TRUE |
| c6 | Female Δ HEL/WT vs Male Δ HEL/WT | -0.1106 | 0e+00 | TRUE |

##### 3.19.7 Liver weight

|  | test | Difference | p\_value | Significance |
| --- | --- | --- | --- | --- |
| c2 | WT/WT vs Δ HEL/WT | 0.1684 | 0.0671 | FALSE |
| c3 | Female vs Male | -0.3739 | 0.0001 | TRUE |
| c4 | Female WT/WT vs Female Δ HEL/WT | 0.2072 | 0.1282 | FALSE |
| c5 | Male WT/WT vs Male Δ HEL/WT | 0.1296 | 0.5243 | FALSE |
| c6 | Female Δ HEL/WT vs Male Δ HEL/WT | -0.4127 | 0.0037 | TRUE |

##### 3.19.8 Lung weight

|  | test | Difference | p\_value | Significance |
| --- | --- | --- | --- | --- |
| c2 | WT/WT vs Δ HEL/WT | 0.0268 | 0.2309 | FALSE |
| c3 | Female vs Male | -0.0254 | 0.2751 | FALSE |
| c4 | Female WT/WT vs Female Δ HEL/WT | 0.0095 | 0.9598 | FALSE |
| c5 | Male WT/WT vs Male Δ HEL/WT | 0.0440 | 0.1387 | FALSE |
| c6 | Female Δ HEL/WT vs Male Δ HEL/WT | -0.0081 | 0.9793 | FALSE |

##### 3.19.9 Right kidney weight

|  | test | Difference | p\_value | Significance |
| --- | --- | --- | --- | --- |
| c2 | WT/WT vs Δ HEL/WT | 0.0424 | 0.0000 | TRUE |
| c3 | Female vs Male | -0.1244 | 0.0000 | TRUE |
| c4 | Female WT/WT vs Female Δ HEL/WT | 0.0363 | 0.0085 | TRUE |
| c5 | Male WT/WT vs Male Δ HEL/WT | 0.0485 | 0.0003 | TRUE |
| c6 | Female Δ HEL/WT vs Male Δ HEL/WT | -0.1183 | 0.0000 | TRUE |

##### 3.19.10 Spleen weight

|  | test | Difference | p\_value | Significance |
| --- | --- | --- | --- | --- |
| c2 | WT/WT vs Δ HEL/WT | 0.0109 | 0.2000 | FALSE |
| c3 | Female vs Male | 0.0373 | 0.0004 | TRUE |
| c4 | Female WT/WT vs Female Δ HEL/WT | 0.0182 | 0.0953 | FALSE |
| c5 | Male WT/WT vs Male Δ HEL/WT | 0.0036 | 0.9648 | FALSE |
| c6 | Female Δ HEL/WT vs Male Δ HEL/WT | 0.0300 | 0.0630 | FALSE |

##### 3.19.11 Testes weight (sin.)

| test | x | p\_value | Significance |
| --- | --- | --- | --- |
| WT vs KO | -0.021708333 | 0.007173557 | TRUE |

##### 3.19.12 Testes weight (dx.)

| test | x | p\_value | Significance |
| --- | --- | --- | --- |
| WT vs KO | -0.021416667 | 0.008770578 | TRUE |

#### 3.20 Immunology

**Number of animals**

| STRAIN | GENDER | count |
| --- | --- | --- |
| WT/WT | female | 8 |
| WT/WT | male | 8 |
| Δ HEL/WT | female | 7 |
| Δ HEL/WT | male | 6 |

##### 3.20.1 B cells - % of live leukocytes (Panel B)

|  | test | Difference | p\_value | Significance |
| --- | --- | --- | --- | --- |
| c2 | WT/WT vs Δ HEL/WT | 4.4207 | 0.0791 | FALSE |
| c3 | Female vs Male | -3.0688 | 0.7309 | FALSE |
| c4 | Female WT/WT vs Female Δ HEL/WT | 3.5416 | 0.4749 | FALSE |
| c5 | Male WT/WT vs Male Δ HEL/WT | 5.2998 | 0.1955 | FALSE |
| c6 | Female Δ HEL/WT vs Male Δ HEL/WT | -2.1897 | 0.9397 | FALSE |

##### 3.20.2 B1a cells - % of B cells (Panel B)

|  | test | Difference | p\_value | Significance |
| --- | --- | --- | --- | --- |
| c2 | WT/WT vs Δ HEL/WT | 0.5632 | 0.3923 | FALSE |
| c3 | Female vs Male | 0.0950 | 0.9937 | FALSE |
| c4 | Female WT/WT vs Female Δ HEL/WT | 0.6935 | 0.4725 | FALSE |
| c5 | Male WT/WT vs Male Δ HEL/WT | 0.4329 | 0.8400 | FALSE |
| c6 | Female Δ HEL/WT vs Male Δ HEL/WT | -0.0353 | 0.9999 | FALSE |

##### 3.20.3 B1a cells - % of live(Panel B)

|  | test | Difference | p\_value | Significance |
| --- | --- | --- | --- | --- |
| c2 | WT/WT vs Δ HEL/WT | 0.3166 | 0.2847 | FALSE |
| c3 | Female vs Male | -0.0220 | 0.9993 | FALSE |
| c4 | Female WT/WT vs Female Δ HEL/WT | 0.3271 | 0.5177 | FALSE |
| c5 | Male WT/WT vs Male Δ HEL/WT | 0.3060 | 0.6432 | FALSE |
| c6 | Female Δ HEL/WT vs Male Δ HEL/WT | -0.0326 | 0.9993 | FALSE |

##### 3.20.4 CD11b- NK cells - % of live leukocytes (Panel B)

|  | test | Difference | p\_value | Significance |
| --- | --- | --- | --- | --- |
| c2 | WT/WT vs Δ HEL/WT | -0.4117 | 0.4862 | FALSE |
| c3 | Female vs Male | 0.1674 | 0.9724 | FALSE |
| c4 | Female WT/WT vs Female Δ HEL/WT | -0.0975 | 0.9943 | FALSE |
| c5 | Male WT/WT vs Male Δ HEL/WT | -0.7259 | 0.3212 | FALSE |
| c6 | Female Δ HEL/WT vs Male Δ HEL/WT | -0.1468 | 0.9920 | FALSE |

##### 3.20.5 CD11b- NK cells - % of NK cells (Panel B)

|  | test | Difference | p\_value | Significance |
| --- | --- | --- | --- | --- |
| c2 | WT/WT vs Δ HEL/WT | 1.2300 | 0.9994 | FALSE |
| c3 | Female vs Male | -13.9414 | 0.5355 | FALSE |
| c4 | Female WT/WT vs Female Δ HEL/WT | 2.6429 | 0.9975 | FALSE |
| c5 | Male WT/WT vs Male Δ HEL/WT | -0.1829 | 1.0000 | FALSE |
| c6 | Female Δ HEL/WT vs Male Δ HEL/WT | -15.3543 | 0.7457 | FALSE |

##### 3.20.6 CD11b-high cDC - % of conventional DC (Panel B)

|  | test | Difference | p\_value | Significance |
| --- | --- | --- | --- | --- |
| c2 | WT/WT vs Δ HEL/WT | -0.1714 | 0.9998 | FALSE |
| c3 | Female vs Male | 1.3000 | 0.9216 | FALSE |
| c4 | Female WT/WT vs Female Δ HEL/WT | -0.0143 | 1.0000 | FALSE |
| c5 | Male WT/WT vs Male Δ HEL/WT | -0.3286 | 0.9995 | FALSE |
| c6 | Female Δ HEL/WT vs Male Δ HEL/WT | 1.1429 | 0.9817 | FALSE |

##### 3.20.7 CD11b-high cDC - % of live leukocytes (Panel B)

|  | test | Difference | p\_value | Significance |
| --- | --- | --- | --- | --- |
| c2 | WT/WT vs Δ HEL/WT | 0.1680 | 0.0028 | TRUE |
| c3 | Female vs Male | -0.0249 | 0.9832 | FALSE |
| c4 | Female WT/WT vs Female Δ HEL/WT | 0.2979 | 0.0000 | TRUE |
| c5 | Male WT/WT vs Male Δ HEL/WT | 0.0381 | 0.9424 | FALSE |
| c6 | Female Δ HEL/WT vs Male Δ HEL/WT | -0.1548 | 0.3284 | FALSE |

##### 3.20.8 CD11b-low cDC - % of conventional DC (Panel B)

|  | test | Difference | p\_value | Significance |
| --- | --- | --- | --- | --- |
| c2 | WT/WT vs Δ HEL/WT | 0.6428 | 0.9754 | FALSE |
| c3 | Female vs Male | -0.7371 | 0.9752 | FALSE |
| c4 | Female WT/WT vs Female Δ HEL/WT | 2.0422 | 0.7613 | FALSE |
| c5 | Male WT/WT vs Male Δ HEL/WT | -0.7567 | 0.9870 | FALSE |
| c6 | Female Δ HEL/WT vs Male Δ HEL/WT | -2.1366 | 0.8259 | FALSE |

##### 3.20.9 CD11b-low cDC - % of of live leukocytes (Panel B)

|  | test | Difference | p\_value | Significance |
| --- | --- | --- | --- | --- |
| c2 | WT/WT vs Δ HEL/WT | 0.0301 | 0.4404 | FALSE |
| c3 | Female vs Male | -0.0068 | 0.9935 | FALSE |
| c4 | Female WT/WT vs Female Δ HEL/WT | 0.0642 | 0.0937 | FALSE |
| c5 | Male WT/WT vs Male Δ HEL/WT | -0.0039 | 0.9991 | FALSE |
| c6 | Female Δ HEL/WT vs Male Δ HEL/WT | -0.0409 | 0.6557 | FALSE |

##### 3.20.10 CD11b+ NK cells - % of live leukocytes (Panel B)

|  | test | Difference | p\_value | Significance |
| --- | --- | --- | --- | --- |
| c2 | WT/WT vs Δ HEL/WT | -0.7143 | 0.7375 | FALSE |
| c3 | Female vs Male | 1.6164 | 0.3633 | FALSE |
| c4 | Female WT/WT vs Female Δ HEL/WT | -0.3065 | 0.9880 | FALSE |
| c5 | Male WT/WT vs Male Δ HEL/WT | -1.1222 | 0.6918 | FALSE |
| c6 | Female Δ HEL/WT vs Male Δ HEL/WT | 1.2086 | 0.7914 | FALSE |

##### 3.20.11 CD11b+ NK cells - % of NK cells (Panel B)

|  | test | Difference | p\_value | Significance |
| --- | --- | --- | --- | --- |
| c2 | WT/WT vs Δ HEL/WT | -1.2357 | 0.9994 | FALSE |
| c3 | Female vs Male | 13.9414 | 0.5356 | FALSE |
| c4 | Female WT/WT vs Female Δ HEL/WT | -2.6486 | 0.9975 | FALSE |
| c5 | Male WT/WT vs Male Δ HEL/WT | 0.1771 | 1.0000 | FALSE |
| c6 | Female Δ HEL/WT vs Male Δ HEL/WT | 15.3543 | 0.7457 | FALSE |

##### 3.20.12 CD161+ B cells - % of B cells (Panel B)

|  | test | Difference | p\_value | Significance |
| --- | --- | --- | --- | --- |
| c2 | WT/WT vs Δ HEL/WT | -0.0658 | 0.2064 | FALSE |
| c3 | Female vs Male | 0.1236 | 0.1120 | FALSE |
| c4 | Female WT/WT vs Female Δ HEL/WT | -0.0411 | 0.7942 | FALSE |
| c5 | Male WT/WT vs Male Δ HEL/WT | -0.0904 | 0.2532 | FALSE |
| c6 | Female Δ HEL/WT vs Male Δ HEL/WT | 0.0990 | 0.4905 | FALSE |

##### 3.20.13 CD161+ B cells - % of live leukocytes (Panel B)

|  | test | Difference | p\_value | Significance |
| --- | --- | --- | --- | --- |
| c2 | WT/WT vs Δ HEL/WT | -0.0293 | 0.2818 | FALSE |
| c3 | Female vs Male | 0.0425 | 0.3456 | FALSE |
| c4 | Female WT/WT vs Female Δ HEL/WT | -0.0176 | 0.8523 | FALSE |
| c5 | Male WT/WT vs Male Δ HEL/WT | -0.0411 | 0.3171 | FALSE |
| c6 | Female Δ HEL/WT vs Male Δ HEL/WT | 0.0307 | 0.7862 | FALSE |

##### 3.20.14 CD4 T cells - % of live leukocytes (Panel A)

|  | test | Difference | p\_value | Significance |
| --- | --- | --- | --- | --- |
| c2 | WT/WT vs Δ HEL/WT | -2.4586 | 0.3210 | FALSE |
| c3 | Female vs Male | -0.3507 | 0.9967 | FALSE |
| c4 | Female WT/WT vs Female Δ HEL/WT | -1.8941 | 0.7491 | FALSE |
| c5 | Male WT/WT vs Male Δ HEL/WT | -3.0231 | 0.4832 | FALSE |
| c6 | Female Δ HEL/WT vs Male Δ HEL/WT | -0.9152 | 0.9802 | FALSE |

##### 3.20.15 CD4- NKT cells - % of live leukocytes (Panel A)

|  | test | Difference | p\_value | Significance |
| --- | --- | --- | --- | --- |
| c2 | WT/WT vs Δ HEL/WT | -0.0104 | 0.9986 | FALSE |
| c3 | Female vs Male | 0.1502 | 0.1532 | FALSE |
| c4 | Female WT/WT vs Female Δ HEL/WT | -0.0317 | 0.9834 | FALSE |
| c5 | Male WT/WT vs Male Δ HEL/WT | 0.0110 | 0.9995 | FALSE |
| c6 | Female Δ HEL/WT vs Male Δ HEL/WT | 0.1715 | 0.3500 | FALSE |

##### 3.20.16 CD4- NKT cells - % of NKT cells (Panel A)

|  | test | Difference | p\_value | Significance |
| --- | --- | --- | --- | --- |
| c2 | WT/WT vs Δ HEL/WT | -11.3318 | 0.2448 | FALSE |
| c3 | Female vs Male | 8.6945 | 0.4951 | FALSE |
| c4 | Female WT/WT vs Female Δ HEL/WT | -15.9909 | 0.1963 | FALSE |
| c5 | Male WT/WT vs Male Δ HEL/WT | -6.6728 | 0.8781 | FALSE |
| c6 | Female Δ HEL/WT vs Male Δ HEL/WT | 13.3536 | 0.4658 | FALSE |

##### 3.20.17 CD4+ NKT cells - % of live leukocytes (Panel A)

|  | test | Difference | p\_value | Significance |
| --- | --- | --- | --- | --- |
| c2 | WT/WT vs Δ HEL/WT | 0.1391 | 0.1181 | FALSE |
| c3 | Female vs Male | 0.1009 | 0.3641 | FALSE |
| c4 | Female WT/WT vs Female Δ HEL/WT | 0.2114 | 0.0552 | FALSE |
| c5 | Male WT/WT vs Male Δ HEL/WT | 0.0669 | 0.8856 | FALSE |
| c6 | Female Δ HEL/WT vs Male Δ HEL/WT | 0.0286 | 0.9894 | FALSE |

##### 3.20.18 CD4+ NKT cells - % of NKT cells (Panel A)

|  | test | Difference | p\_value | Significance |
| --- | --- | --- | --- | --- |
| c2 | WT/WT vs Δ HEL/WT | 11.3224 | 0.2507 | FALSE |
| c3 | Female vs Male | -8.5824 | 0.5085 | FALSE |
| c4 | Female WT/WT vs Female Δ HEL/WT | 15.8749 | 0.2064 | FALSE |
| c5 | Male WT/WT vs Male Δ HEL/WT | 6.7700 | 0.8757 | FALSE |
| c6 | Female Δ HEL/WT vs Male Δ HEL/WT | -13.1348 | 0.4836 | FALSE |

##### 3.20.19 CD4+ T cells - % of T cells (Panel A)

|  | test | Difference | p\_value | Significance |
| --- | --- | --- | --- | --- |
| c2 | WT/WT vs Δ HEL/WT | -0.8317 | 0.9943 | FALSE |
| c3 | Female vs Male | 2.6819 | 0.9087 | FALSE |
| c4 | Female WT/WT vs Female Δ HEL/WT | 0.2302 | 0.9999 | FALSE |
| c5 | Male WT/WT vs Male Δ HEL/WT | -1.8936 | 0.9800 | FALSE |
| c6 | Female Δ HEL/WT vs Male Δ HEL/WT | 1.6200 | 0.9916 | FALSE |

##### 3.20.20 CD4+ T helper cells - % of CD4 T cells (Panel A)

|  | test | Difference | p\_value | Significance |
| --- | --- | --- | --- | --- |
| c2 | WT/WT vs Δ HEL/WT | -0.9725 | 0.1080 | FALSE |
| c3 | Female vs Male | 0.5275 | 0.5968 | FALSE |
| c4 | Female WT/WT vs Female Δ HEL/WT | 0.2464 | 0.9708 | FALSE |
| c5 | Male WT/WT vs Male Δ HEL/WT | -2.1914 | 0.0046 | TRUE |
| c6 | Female Δ HEL/WT vs Male Δ HEL/WT | -0.6914 | 0.6935 | FALSE |

##### 3.20.21 CD4+ T helper cells - %oflive leukocytes (Panel A

|  | test | Difference | p\_value | Significance |
| --- | --- | --- | --- | --- |
| c2 | WT/WT vs Δ HEL/WT | -2.4832 | 0.2692 | FALSE |
| c3 | Female vs Male | -0.2931 | 0.9977 | FALSE |
| c4 | Female WT/WT vs Female Δ HEL/WT | -1.6931 | 0.7819 | FALSE |
| c5 | Male WT/WT vs Male Δ HEL/WT | -3.2734 | 0.3688 | FALSE |
| c6 | Female Δ HEL/WT vs Male Δ HEL/WT | -1.0832 | 0.9622 | FALSE |

##### 3.20.22 CD8+ T cells - % of live leukocytes (Panel A)

|  | test | Difference | p\_value | Significance |
| --- | --- | --- | --- | --- |
| c2 | WT/WT vs Δ HEL/WT | -0.7481 | 0.8629 | FALSE |
| c3 | Female vs Male | -0.5224 | 0.9474 | FALSE |
| c4 | Female WT/WT vs Female Δ HEL/WT | -1.2111 | 0.7760 | FALSE |
| c5 | Male WT/WT vs Male Δ HEL/WT | -0.2851 | 0.9972 | FALSE |
| c6 | Female Δ HEL/WT vs Male Δ HEL/WT | -0.0594 | 1.0000 | FALSE |

##### 3.20.23 CD8+ T cells - % of T cells (Panel A)

|  | test | Difference | p\_value | Significance |
| --- | --- | --- | --- | --- |
| c2 | WT/WT vs Δ HEL/WT | 0.6451 | 0.9972 | FALSE |
| c3 | Female vs Male | -2.6172 | 0.9083 | FALSE |
| c4 | Female WT/WT vs Female Δ HEL/WT | -0.5025 | 0.9994 | FALSE |
| c5 | Male WT/WT vs Male Δ HEL/WT | 1.7927 | 0.9822 | FALSE |
| c6 | Female Δ HEL/WT vs Male Δ HEL/WT | -1.4697 | 0.9933 | FALSE |

##### 3.20.24 Conventional DC - % of live leukocytes (Panel B)

|  | test | Difference | p\_value | Significance |
| --- | --- | --- | --- | --- |
| c2 | WT/WT vs Δ HEL/WT | 0.2170 | 0.0014 | TRUE |
| c3 | Female vs Male | -0.0300 | 0.9810 | FALSE |
| c4 | Female WT/WT vs Female Δ HEL/WT | 0.3796 | 0.0000 | TRUE |
| c5 | Male WT/WT vs Male Δ HEL/WT | 0.0544 | 0.9096 | FALSE |
| c6 | Female Δ HEL/WT vs Male Δ HEL/WT | -0.1926 | 0.2739 | FALSE |

##### 3.20.25 Effector CD4- NKT c. - % of CD4- NKT c. (Panel A)

|  | test | Difference | p\_value | Significance |
| --- | --- | --- | --- | --- |
| c2 | WT/WT vs Δ HEL/WT | 9.0932 | 0.2433 | FALSE |
| c3 | Female vs Male | -4.9282 | 0.7347 | FALSE |
| c4 | Female WT/WT vs Female Δ HEL/WT | 12.3893 | 0.2212 | FALSE |
| c5 | Male WT/WT vs Male Δ HEL/WT | 5.7971 | 0.8512 | FALSE |
| c6 | Female Δ HEL/WT vs Male Δ HEL/WT | -8.2243 | 0.6648 | FALSE |

##### 3.20.26 Effector CD4- NKT cells - % of live (Panel A)

|  | test | Difference | p\_value | Significance |
| --- | --- | --- | --- | --- |
| c2 | WT/WT vs Δ HEL/WT | 0.0307 | 0.5536 | FALSE |
| c3 | Female vs Male | 0.0605 | 0.0841 | FALSE |
| c4 | Female WT/WT vs Female Δ HEL/WT | 0.0385 | 0.5991 | FALSE |
| c5 | Male WT/WT vs Male Δ HEL/WT | 0.0229 | 0.9101 | FALSE |
| c6 | Female Δ HEL/WT vs Male Δ HEL/WT | 0.0527 | 0.4762 | FALSE |

##### 3.20.27 Effector CD4+ NKT c. - % of CD4+ NKT c. (Panel A)

|  | test | Difference | p\_value | Significance |
| --- | --- | --- | --- | --- |
| c2 | WT/WT vs Δ HEL/WT | 13.7384 | 0.3673 | FALSE |
| c3 | Female vs Male | -3.6259 | 0.9725 | FALSE |
| c4 | Female WT/WT vs Female Δ HEL/WT | 13.2339 | 0.6351 | FALSE |
| c5 | Male WT/WT vs Male Δ HEL/WT | 14.2429 | 0.6709 | FALSE |
| c6 | Female Δ HEL/WT vs Male Δ HEL/WT | -3.1214 | 0.9945 | FALSE |

##### 3.20.28 Effector CD4+ NKT cells - % of live (Panel A)

|  | test | Difference | p\_value | Significance |
| --- | --- | --- | --- | --- |
| c2 | WT/WT vs Δ HEL/WT | 0.1498 | 0.0413 | TRUE |
| c3 | Female vs Male | 0.0789 | 0.4865 | FALSE |
| c4 | Female WT/WT vs Female Δ HEL/WT | 0.2056 | 0.0322 | TRUE |
| c5 | Male WT/WT vs Male Δ HEL/WT | 0.0940 | 0.6680 | FALSE |
| c6 | Female Δ HEL/WT vs Male Δ HEL/WT | 0.0230 | 0.9923 | FALSE |

##### 3.20.29 Effector CD4+ T help c.-% of CD4+ T help (Panel A)

|  | test | Difference | p\_value | Significance |
| --- | --- | --- | --- | --- |
| c2 | WT/WT vs Δ HEL/WT | 6.4015 | 0.0008 | TRUE |
| c3 | Female vs Male | 10.3422 | 0.0001 | TRUE |
| c4 | Female WT/WT vs Female Δ HEL/WT | 4.4402 | 0.1801 | FALSE |
| c5 | Male WT/WT vs Male Δ HEL/WT | 8.3628 | 0.0037 | TRUE |
| c6 | Female Δ HEL/WT vs Male Δ HEL/WT | 12.3036 | 0.0008 | TRUE |

##### 3.20.30 Effector CD4+ T helper cells - % of live (Panel A)

|  | test | Difference | p\_value | Significance |
| --- | --- | --- | --- | --- |
| c2 | WT/WT vs Δ HEL/WT | 0.4829 | 0.3511 | FALSE |
| c3 | Female vs Male | 2.5807 | 0.0000 | TRUE |
| c4 | Female WT/WT vs Female Δ HEL/WT | 0.0867 | 0.9960 | FALSE |
| c5 | Male WT/WT vs Male Δ HEL/WT | 0.8791 | 0.1778 | FALSE |
| c6 | Female Δ HEL/WT vs Male Δ HEL/WT | 2.9769 | 0.0001 | TRUE |

##### 3.20.31 Effector CD8+ T cells - % of CD8+ T cells (Panel A

|  | test | Difference | p\_value | Significance |
| --- | --- | --- | --- | --- |
| c2 | WT/WT vs Δ HEL/WT | 1.1576 | 0.7706 | FALSE |
| c3 | Female vs Male | 8.3663 | 0.0004 | TRUE |
| c4 | Female WT/WT vs Female Δ HEL/WT | 1.4299 | 0.8161 | FALSE |
| c5 | Male WT/WT vs Male Δ HEL/WT | 0.8854 | 0.9576 | FALSE |
| c6 | Female Δ HEL/WT vs Male Δ HEL/WT | 8.0940 | 0.0180 | TRUE |

##### 3.20.32 Effector CD8+ T cells - %oflive leukocytes (Pane A

|  | test | Difference | p\_value | Significance |
| --- | --- | --- | --- | --- |
| c2 | WT/WT vs Δ HEL/WT | 0.0556 | 0.7539 | FALSE |
| c3 | Female vs Male | 0.2329 | 0.0715 | FALSE |
| c4 | Female WT/WT vs Female Δ HEL/WT | 0.0899 | 0.6399 | FALSE |
| c5 | Male WT/WT vs Male Δ HEL/WT | 0.0213 | 0.9937 | FALSE |
| c6 | Female Δ HEL/WT vs Male Δ HEL/WT | 0.1986 | 0.3736 | FALSE |

##### 3.20.33 Effector NK cells - % of live leukocytes (Panel A)

|  | test | Difference | p\_value | Significance |
| --- | --- | --- | --- | --- |
| c2 | WT/WT vs Δ HEL/WT | 0.0816 | 0.9017 | FALSE |
| c3 | Female vs Male | 0.5073 | 0.0452 | TRUE |
| c4 | Female WT/WT vs Female Δ HEL/WT | 0.1738 | 0.7032 | FALSE |
| c5 | Male WT/WT vs Male Δ HEL/WT | -0.0107 | 0.9999 | FALSE |
| c6 | Female Δ HEL/WT vs Male Δ HEL/WT | 0.4151 | 0.3526 | FALSE |

##### 3.20.34 Effector NK cells - % of NK cells (Panel A)

|  | test | Difference | p\_value | Significance |
| --- | --- | --- | --- | --- |
| c2 | WT/WT vs Δ HEL/WT | 5.4861 | 0.0000 | TRUE |
| c3 | Female vs Male | 1.1614 | 0.9061 | FALSE |
| c4 | Female WT/WT vs Female Δ HEL/WT | 4.8049 | 0.0132 | TRUE |
| c5 | Male WT/WT vs Male Δ HEL/WT | 6.1673 | 0.0022 | TRUE |
| c6 | Female Δ HEL/WT vs Male Δ HEL/WT | 1.8426 | 0.8525 | FALSE |

##### 3.20.35 Effector Treg cells - % of live leukocytes (Panel A)

|  | test | Difference | p\_value | Significance |
| --- | --- | --- | --- | --- |
| c2 | WT/WT vs Δ HEL/WT | 0.0744 | 0.2147 | FALSE |
| c3 | Female vs Male | 0.1026 | 0.3018 | FALSE |
| c4 | Female WT/WT vs Female Δ HEL/WT | -0.0062 | 0.9993 | FALSE |
| c5 | Male WT/WT vs Male Δ HEL/WT | 0.1549 | 0.0353 | TRUE |
| c6 | Female Δ HEL/WT vs Male Δ HEL/WT | 0.1831 | 0.0968 | FALSE |

##### 3.20.36 Effector Treg cells - % of Treg cells (Panel A)

|  | test | Difference | p\_value | Significance |
| --- | --- | --- | --- | --- |
| c2 | WT/WT vs Δ HEL/WT | 5.4316 | 0.0097 | TRUE |
| c3 | Female vs Male | 7.0321 | 0.0327 | TRUE |
| c4 | Female WT/WT vs Female Δ HEL/WT | 5.0730 | 0.1227 | FALSE |
| c5 | Male WT/WT vs Male Δ HEL/WT | 5.7903 | 0.0983 | FALSE |
| c6 | Female Δ HEL/WT vs Male Δ HEL/WT | 7.3907 | 0.1325 | FALSE |

##### 3.20.37 Eosinophils - % of live leukocytes (Panel B)

|  | test | Difference | p\_value | Significance |
| --- | --- | --- | --- | --- |
| c2 | WT/WT vs Δ HEL/WT | 0.0504 | 0.9241 | FALSE |
| c3 | Female vs Male | 0.2242 | 0.1683 | FALSE |
| c4 | Female WT/WT vs Female Δ HEL/WT | 0.0643 | 0.9347 | FALSE |
| c5 | Male WT/WT vs Male Δ HEL/WT | 0.0364 | 0.9897 | FALSE |
| c6 | Female Δ HEL/WT vs Male Δ HEL/WT | 0.2102 | 0.4726 | FALSE |

##### 3.20.38 Follicular B cells - % of B cells (Panel B)

|  | test | Difference | p\_value | Significance |
| --- | --- | --- | --- | --- |
| c2 | WT/WT vs Δ HEL/WT | -0.3857 | 0.9713 | FALSE |
| c3 | Female vs Male | -1.2571 | 0.4862 | FALSE |
| c4 | Female WT/WT vs Female Δ HEL/WT | -1.7571 | 0.4548 | FALSE |
| c5 | Male WT/WT vs Male Δ HEL/WT | 0.9857 | 0.8718 | FALSE |
| c6 | Female Δ HEL/WT vs Male Δ HEL/WT | 0.1143 | 0.9998 | FALSE |

##### 3.20.39 Follicular B cells - % of B cells LIVE (Panel B)

|  | test | Difference | p\_value | Significance |
| --- | --- | --- | --- | --- |
| c2 | WT/WT vs Δ HEL/WT | 3.1091 | 0.1866 | FALSE |
| c3 | Female vs Male | -3.5527 | 0.4842 | FALSE |
| c4 | Female WT/WT vs Female Δ HEL/WT | 1.8494 | 0.8054 | FALSE |
| c5 | Male WT/WT vs Male Δ HEL/WT | 4.3688 | 0.2146 | FALSE |
| c6 | Female Δ HEL/WT vs Male Δ HEL/WT | -2.2929 | 0.8891 | FALSE |

##### 3.20.40 Granulocytes - % of live leukocytes (Panel B)

|  | test | Difference | p\_value | Significance |
| --- | --- | --- | --- | --- |
| c2 | WT/WT vs Δ HEL/WT | 0.2886 | 0.6983 | FALSE |
| c3 | Female vs Male | 0.8961 | 0.1651 | FALSE |
| c4 | Female WT/WT vs Female Δ HEL/WT | 0.3716 | 0.7313 | FALSE |
| c5 | Male WT/WT vs Male Δ HEL/WT | 0.2055 | 0.9507 | FALSE |
| c6 | Female Δ HEL/WT vs Male Δ HEL/WT | 0.8130 | 0.4605 | FALSE |

##### 3.20.41 Live leukocytes (Panel A) - % of total events

|  | test | Difference | p\_value | Significance |
| --- | --- | --- | --- | --- |
| c2 | WT/WT vs Δ HEL/WT | 4.0820 | 0.1556 | FALSE |
| c3 | Female vs Male | -4.2895 | 0.1243 | FALSE |
| c4 | Female WT/WT vs Female Δ HEL/WT | 1.2125 | 0.9633 | FALSE |
| c5 | Male WT/WT vs Male Δ HEL/WT | 6.9514 | 0.0837 | FALSE |
| c6 | Female Δ HEL/WT vs Male Δ HEL/WT | -1.4200 | 0.9596 | FALSE |

##### 3.20.42 Live leukocytes (Panel B) - % of total events

|  | test | Difference | p\_value | Significance |
| --- | --- | --- | --- | --- |
| c2 | WT/WT vs Δ HEL/WT | 4.6603 | 0.2605 | FALSE |
| c3 | Female vs Male | -8.6319 | 0.0791 | FALSE |
| c4 | Female WT/WT vs Female Δ HEL/WT | 1.5898 | 0.9652 | FALSE |
| c5 | Male WT/WT vs Male Δ HEL/WT | 7.7308 | 0.1647 | FALSE |
| c6 | Female Δ HEL/WT vs Male Δ HEL/WT | -5.5614 | 0.6375 | FALSE |

##### 3.20.43 Ly6C+ CD11b- NK cells - % of live (Panel B)

|  | test | Difference | p\_value | Significance |
| --- | --- | --- | --- | --- |
| c2 | WT/WT vs Δ HEL/WT | -0.0755 | 0.5610 | FALSE |
| c3 | Female vs Male | -0.0216 | 0.9876 | FALSE |
| c4 | Female WT/WT vs Female Δ HEL/WT | 0.0188 | 0.9949 | FALSE |
| c5 | Male WT/WT vs Male Δ HEL/WT | -0.1698 | 0.1985 | FALSE |
| c6 | Female Δ HEL/WT vs Male Δ HEL/WT | -0.1159 | 0.5972 | FALSE |

##### 3.20.44 Ly6C+ CD11b- NK cells - % of NK cells (Panel B)

|  | test | Difference | p\_value | Significance |
| --- | --- | --- | --- | --- |
| c2 | WT/WT vs Δ HEL/WT | -0.0970 | 0.9995 | FALSE |
| c3 | Female vs Male | -1.9856 | 0.1038 | FALSE |
| c4 | Female WT/WT vs Female Δ HEL/WT | 0.6257 | 0.9477 | FALSE |
| c5 | Male WT/WT vs Male Δ HEL/WT | -0.8197 | 0.9149 | FALSE |
| c6 | Female Δ HEL/WT vs Male Δ HEL/WT | -2.7083 | 0.1506 | FALSE |

##### 3.20.45 Ly6C+ CD11b+ NK cells - % of live (Panel B)

|  | test | Difference | p\_value | Significance |
| --- | --- | --- | --- | --- |
| c2 | WT/WT vs Δ HEL/WT | -0.1469 | 0.9500 | FALSE |
| c3 | Female vs Male | 0.6043 | 0.4077 | FALSE |
| c4 | Female WT/WT vs Female Δ HEL/WT | 0.0318 | 0.9998 | FALSE |
| c5 | Male WT/WT vs Male Δ HEL/WT | -0.3257 | 0.8487 | FALSE |
| c6 | Female Δ HEL/WT vs Male Δ HEL/WT | 0.4256 | 0.8404 | FALSE |

##### 3.20.46 Ly6C+ CD11b+ NK cells - % of NK cells (Panel B)

|  | test | Difference | p\_value | Significance |
| --- | --- | --- | --- | --- |
| c2 | WT/WT vs Δ HEL/WT | 2.1306 | 0.9675 | FALSE |
| c3 | Female vs Male | 4.8706 | 0.7215 | FALSE |
| c4 | Female WT/WT vs Female Δ HEL/WT | 1.9971 | 0.9886 | FALSE |
| c5 | Male WT/WT vs Male Δ HEL/WT | 2.2640 | 0.9874 | FALSE |
| c6 | Female Δ HEL/WT vs Male Δ HEL/WT | 5.0040 | 0.8845 | FALSE |

##### 3.20.47 Ly6C+ NKT cells - % of live leukocytes (Panel B)

|  | test | Difference | p\_value | Significance |
| --- | --- | --- | --- | --- |
| c2 | WT/WT vs Δ HEL/WT | 0.0628 | 0.8536 | FALSE |
| c3 | Female vs Male | 0.1749 | 0.1284 | FALSE |
| c4 | Female WT/WT vs Female Δ HEL/WT | 0.1389 | 0.5598 | FALSE |
| c5 | Male WT/WT vs Male Δ HEL/WT | -0.0132 | 0.9995 | FALSE |
| c6 | Female Δ HEL/WT vs Male Δ HEL/WT | 0.0988 | 0.8289 | FALSE |

##### 3.20.48 Ly6C+ NKT cells - % of NKT cells (Panel B)

|  | test | Difference | p\_value | Significance |
| --- | --- | --- | --- | --- |
| c2 | WT/WT vs Δ HEL/WT | -6.9440 | 0.7296 | FALSE |
| c3 | Female vs Male | 0.7360 | 0.9995 | FALSE |
| c4 | Female WT/WT vs Female Δ HEL/WT | 0.9343 | 0.9996 | FALSE |
| c5 | Male WT/WT vs Male Δ HEL/WT | -14.8223 | 0.4472 | FALSE |
| c6 | Female Δ HEL/WT vs Male Δ HEL/WT | -7.1423 | 0.8881 | FALSE |

##### 3.20.49 Marginal zone B cells - % B cells LIVE (Panel B)

|  | test | Difference | p\_value | Significance |
| --- | --- | --- | --- | --- |
| c2 | WT/WT vs Δ HEL/WT | 0.1682 | 0.9570 | FALSE |
| c3 | Female vs Male | 0.7197 | 0.5992 | FALSE |
| c4 | Female WT/WT vs Female Δ HEL/WT | 0.4694 | 0.7300 | FALSE |
| c5 | Male WT/WT vs Male Δ HEL/WT | -0.1330 | 0.9927 | FALSE |
| c6 | Female Δ HEL/WT vs Male Δ HEL/WT | 0.4185 | 0.9407 | FALSE |

##### 3.20.50 Marginal zone B cells - % of B cells (Panel B)

|  | test | Difference | p\_value | Significance |
| --- | --- | --- | --- | --- |
| c2 | WT/WT vs Δ HEL/WT | -0.4758 | 0.8752 | FALSE |
| c3 | Female vs Male | 2.1288 | 0.2108 | FALSE |
| c4 | Female WT/WT vs Female Δ HEL/WT | 0.2088 | 0.9949 | FALSE |
| c5 | Male WT/WT vs Male Δ HEL/WT | -1.1604 | 0.5974 | FALSE |
| c6 | Female Δ HEL/WT vs Male Δ HEL/WT | 1.4442 | 0.7225 | FALSE |

##### 3.20.51 Monocytes - % of live leukocytes (Panel B)

|  | test | Difference | p\_value | Significance |
| --- | --- | --- | --- | --- |
| c2 | WT/WT vs Δ HEL/WT | 0.0883 | 0.8989 | FALSE |
| c3 | Female vs Male | -0.2189 | 0.3240 | FALSE |
| c4 | Female WT/WT vs Female Δ HEL/WT | 0.0900 | 0.9527 | FALSE |
| c5 | Male WT/WT vs Male Δ HEL/WT | 0.0866 | 0.9672 | FALSE |
| c6 | Female Δ HEL/WT vs Male Δ HEL/WT | -0.2206 | 0.6461 | FALSE |

##### 3.20.52 Naive CD8+ T cells - % of CD8+ T cells (Panel A)

|  | test | Difference | p\_value | Significance |
| --- | --- | --- | --- | --- |
| c2 | WT/WT vs Δ HEL/WT | -5.3214 | 0.3534 | FALSE |
| c3 | Female vs Male | -17.4441 | 0.0072 | TRUE |
| c4 | Female WT/WT vs Female Δ HEL/WT | -6.1069 | 0.4970 | FALSE |
| c5 | Male WT/WT vs Male Δ HEL/WT | -4.5358 | 0.7653 | FALSE |
| c6 | Female Δ HEL/WT vs Male Δ HEL/WT | -16.6585 | 0.0842 | FALSE |

##### 3.20.53 Naive CD8+ T cells - % of live leukocytes (Panel A

|  | test | Difference | p\_value | Significance |
| --- | --- | --- | --- | --- |
| c2 | WT/WT vs Δ HEL/WT | -0.9382 | 0.6791 | FALSE |
| c3 | Female vs Male | -1.9330 | 0.3458 | FALSE |
| c4 | Female WT/WT vs Female Δ HEL/WT | -1.2120 | 0.7027 | FALSE |
| c5 | Male WT/WT vs Male Δ HEL/WT | -0.6643 | 0.9484 | FALSE |
| c6 | Female Δ HEL/WT vs Male Δ HEL/WT | -1.6591 | 0.7146 | FALSE |

##### 3.20.54 NK cells (Panel A) - % of live l(Panel A)

|  | test | Difference | p\_value | Significance |
| --- | --- | --- | --- | --- |
| c2 | WT/WT vs Δ HEL/WT | -1.3504 | 0.3775 | FALSE |
| c3 | Female vs Male | 2.0114 | 0.3451 | FALSE |
| c4 | Female WT/WT vs Female Δ HEL/WT | -0.8611 | 0.8660 | FALSE |
| c5 | Male WT/WT vs Male Δ HEL/WT | -1.8397 | 0.4431 | FALSE |
| c6 | Female Δ HEL/WT vs Male Δ HEL/WT | 1.5221 | 0.7797 | FALSE |

##### 3.20.55 NK cells (Panel B) - % of live leukocytes (Panel B

|  | test | Difference | p\_value | Significance |
| --- | --- | --- | --- | --- |
| c2 | WT/WT vs Δ HEL/WT | -1.1345 | 0.6203 | FALSE |
| c3 | Female vs Male | 1.8564 | 0.4980 | FALSE |
| c4 | Female WT/WT vs Female Δ HEL/WT | -0.4260 | 0.9864 | FALSE |
| c5 | Male WT/WT vs Male Δ HEL/WT | -1.8430 | 0.5354 | FALSE |
| c6 | Female Δ HEL/WT vs Male Δ HEL/WT | 1.1479 | 0.9123 | FALSE |

##### 3.20.56 NKT cells (panel A) - % of live (Panel A)

|  | test | Difference | p\_value | Significance |
| --- | --- | --- | --- | --- |
| c2 | WT/WT vs Δ HEL/WT | 0.1348 | 0.6815 | FALSE |
| c3 | Female vs Male | 0.2457 | 0.1890 | FALSE |
| c4 | Female WT/WT vs Female Δ HEL/WT | 0.1892 | 0.6407 | FALSE |
| c5 | Male WT/WT vs Male Δ HEL/WT | 0.0804 | 0.9703 | FALSE |
| c6 | Female Δ HEL/WT vs Male Δ HEL/WT | 0.1913 | 0.7170 | FALSE |

##### 3.20.57 NKT cells (panel B) - % of live leukocytes (Panel

|  | test | Difference | p\_value | Significance |
| --- | --- | --- | --- | --- |
| c2 | WT/WT vs Δ HEL/WT | 0.2175 | 0.4931 | FALSE |
| c3 | Female vs Male | 0.3993 | 0.0546 | FALSE |
| c4 | Female WT/WT vs Female Δ HEL/WT | 0.3036 | 0.4628 | FALSE |
| c5 | Male WT/WT vs Male Δ HEL/WT | 0.1313 | 0.9373 | FALSE |
| c6 | Female Δ HEL/WT vs Male Δ HEL/WT | 0.3131 | 0.5151 | FALSE |

##### 3.20.58 Resting CD4- NKT c. - % of CD4- NKT c. (Panel A)

|  | test | Difference | p\_value | Significance |
| --- | --- | --- | --- | --- |
| c2 | WT/WT vs Δ HEL/WT | -7.2358 | 0.6605 | FALSE |
| c3 | Female vs Male | 12.4642 | 0.2082 | FALSE |
| c4 | Female WT/WT vs Female Δ HEL/WT | -10.5804 | 0.5862 | FALSE |
| c5 | Male WT/WT vs Male Δ HEL/WT | -3.8911 | 0.9759 | FALSE |
| c6 | Female Δ HEL/WT vs Male Δ HEL/WT | 15.8089 | 0.3466 | FALSE |

##### 3.20.59 Resting CD4- NKT cells - % of live (Panel A)

|  | test | Difference | p\_value | Significance |
| --- | --- | --- | --- | --- |
| c2 | WT/WT vs Δ HEL/WT | -0.0408 | 0.7922 | FALSE |
| c3 | Female vs Male | 0.0858 | 0.2561 | FALSE |
| c4 | Female WT/WT vs Female Δ HEL/WT | -0.0719 | 0.6127 | FALSE |
| c5 | Male WT/WT vs Male Δ HEL/WT | -0.0097 | 0.9988 | FALSE |
| c6 | Female Δ HEL/WT vs Male Δ HEL/WT | 0.1169 | 0.3235 | FALSE |

##### 3.20.60 Resting CD4+ NKT c. - % of CD4+ NKT c. (Panel A)

|  | test | Difference | p\_value | Significance |
| --- | --- | --- | --- | --- |
| c2 | WT/WT vs Δ HEL/WT | -11.1394 | 0.0358 | TRUE |
| c3 | Female vs Male | 2.3206 | 0.9380 | FALSE |
| c4 | Female WT/WT vs Female Δ HEL/WT | -10.5788 | 0.2048 | FALSE |
| c5 | Male WT/WT vs Male Δ HEL/WT | -11.7000 | 0.2220 | FALSE |
| c6 | Female Δ HEL/WT vs Male Δ HEL/WT | 1.7600 | 0.9911 | FALSE |

##### 3.20.61 Resting CD4+ NKT cells - % of live (Panel A)

|  | test | Difference | p\_value | Significance |
| --- | --- | --- | --- | --- |
| c2 | WT/WT vs Δ HEL/WT | -0.0081 | 0.6488 | FALSE |
| c3 | Female vs Male | 0.0162 | 0.1025 | FALSE |
| c4 | Female WT/WT vs Female Δ HEL/WT | 0.0033 | 0.9838 | FALSE |
| c5 | Male WT/WT vs Male Δ HEL/WT | -0.0195 | 0.2500 | FALSE |
| c6 | Female Δ HEL/WT vs Male Δ HEL/WT | 0.0048 | 0.9674 | FALSE |

##### 3.20.62 Resting CD4+ T help c. - % of CD4+ T help (Panel

|  | test | Difference | p\_value | Significance |
| --- | --- | --- | --- | --- |
| c2 | WT/WT vs Δ HEL/WT | -6.5498 | 0.0004 | TRUE |
| c3 | Female vs Male | -11.3385 | 0.0000 | TRUE |
| c4 | Female WT/WT vs Female Δ HEL/WT | -4.9817 | 0.1004 | FALSE |
| c5 | Male WT/WT vs Male Δ HEL/WT | -8.1179 | 0.0041 | TRUE |
| c6 | Female Δ HEL/WT vs Male Δ HEL/WT | -12.9066 | 0.0005 | TRUE |

##### 3.20.63 Resting CD4+ T helper cells - % of live (Panel A)

|  | test | Difference | p\_value | Significance |
| --- | --- | --- | --- | --- |
| c2 | WT/WT vs Δ HEL/WT | -3.1115 | 0.0392 | TRUE |
| c3 | Female vs Male | -2.1172 | 0.2568 | FALSE |
| c4 | Female WT/WT vs Female Δ HEL/WT | -2.2288 | 0.4570 | FALSE |
| c5 | Male WT/WT vs Male Δ HEL/WT | -3.9943 | 0.1003 | FALSE |
| c6 | Female Δ HEL/WT vs Male Δ HEL/WT | -3.0000 | 0.3040 | FALSE |

##### 3.20.64 Resting CD8+ T cells - % of CD8+ T cells (Panel A)

|  | test | Difference | p\_value | Significance |
| --- | --- | --- | --- | --- |
| c2 | WT/WT vs Δ HEL/WT | 3.3695 | 0.5316 | FALSE |
| c3 | Female vs Male | 0.5480 | 0.9961 | FALSE |
| c4 | Female WT/WT vs Female Δ HEL/WT | 3.2732 | 0.7524 | FALSE |
| c5 | Male WT/WT vs Male Δ HEL/WT | 3.4657 | 0.7879 | FALSE |
| c6 | Female Δ HEL/WT vs Male Δ HEL/WT | 0.6443 | 0.9981 | FALSE |

##### 3.20.65 Resting CD8+ T cells - %of live (Panel A)

|  | test | Difference | p\_value | Significance |
| --- | --- | --- | --- | --- |
| c2 | WT/WT vs Δ HEL/WT | 0.1514 | 0.7694 | FALSE |
| c3 | Female vs Male | -0.0072 | 1.0000 | FALSE |
| c4 | Female WT/WT vs Female Δ HEL/WT | 0.0634 | 0.9900 | FALSE |
| c5 | Male WT/WT vs Male Δ HEL/WT | 0.2394 | 0.7391 | FALSE |
| c6 | Female Δ HEL/WT vs Male Δ HEL/WT | 0.0809 | 0.9858 | FALSE |

##### 3.20.66 Resting NK cells - % of live leukocytes (Panel A)

|  | test | Difference | p\_value | Significance |
| --- | --- | --- | --- | --- |
| c2 | WT/WT vs Δ HEL/WT | -1.4129 | 0.2235 | FALSE |
| c3 | Female vs Male | 1.3755 | 0.5174 | FALSE |
| c4 | Female WT/WT vs Female Δ HEL/WT | -1.0233 | 0.7178 | FALSE |
| c5 | Male WT/WT vs Male Δ HEL/WT | -1.8025 | 0.3406 | FALSE |
| c6 | Female Δ HEL/WT vs Male Δ HEL/WT | 0.9859 | 0.8848 | FALSE |

##### 3.20.67 Resting NK cells - % of NK cells (Panel A)

|  | test | Difference | p\_value | Significance |
| --- | --- | --- | --- | --- |
| c2 | WT/WT vs Δ HEL/WT | -8.8541 | 0.3036 | FALSE |
| c3 | Female vs Male | -1.6684 | 0.9872 | FALSE |
| c4 | Female WT/WT vs Female Δ HEL/WT | -6.8339 | 0.7346 | FALSE |
| c5 | Male WT/WT vs Male Δ HEL/WT | -10.8743 | 0.4788 | FALSE |
| c6 | Female Δ HEL/WT vs Male Δ HEL/WT | -3.6886 | 0.9609 | FALSE |

##### 3.20.68 Resting Treg cells - % of live (Panel A)

|  | test | Difference | p\_value | Significance |
| --- | --- | --- | --- | --- |
| c2 | WT/WT vs Δ HEL/WT | -0.0512 | 0.9261 | FALSE |
| c3 | Female vs Male | -0.1483 | 0.2914 | FALSE |
| c4 | Female WT/WT vs Female Δ HEL/WT | -0.2089 | 0.2405 | FALSE |
| c5 | Male WT/WT vs Male Δ HEL/WT | 0.1066 | 0.8266 | FALSE |
| c6 | Female Δ HEL/WT vs Male Δ HEL/WT | 0.0094 | 0.9998 | FALSE |

##### 3.20.69 Resting Treg cells - % of Treg cells (Panel A)

|  | test | Difference | p\_value | Significance |
| --- | --- | --- | --- | --- |
| c2 | WT/WT vs Δ HEL/WT | -5.5197 | 0.0108 | TRUE |
| c3 | Female vs Male | -7.4452 | 0.0266 | TRUE |
| c4 | Female WT/WT vs Female Δ HEL/WT | -5.1598 | 0.1269 | FALSE |
| c5 | Male WT/WT vs Male Δ HEL/WT | -5.8796 | 0.1023 | FALSE |
| c6 | Female Δ HEL/WT vs Male Δ HEL/WT | -7.8052 | 0.1181 | FALSE |

##### 3.20.70 T cells (Panel A) - % of live leukocytes (Panel A)

|  | test | Difference | p\_value | Significance |
| --- | --- | --- | --- | --- |
| c2 | WT/WT vs Δ HEL/WT | -3.1367 | 0.1109 | FALSE |
| c3 | Female vs Male | -1.4087 | 0.8121 | FALSE |
| c4 | Female WT/WT vs Female Δ HEL/WT | -2.9003 | 0.3884 | FALSE |
| c5 | Male WT/WT vs Male Δ HEL/WT | -3.3731 | 0.3493 | FALSE |
| c6 | Female Δ HEL/WT vs Male Δ HEL/WT | -1.6450 | 0.8850 | FALSE |

##### 3.20.71 T cells (panel B) - % of live leukocytes (Panel B)

|  | test | Difference | p\_value | Significance |
| --- | --- | --- | --- | --- |
| c2 | WT/WT vs Δ HEL/WT | -3.5600 | 0.1312 | FALSE |
| c3 | Female vs Male | -1.4606 | 0.8614 | FALSE |
| c4 | Female WT/WT vs Female Δ HEL/WT | -3.9583 | 0.2715 | FALSE |
| c5 | Male WT/WT vs Male Δ HEL/WT | -3.1617 | 0.5401 | FALSE |
| c6 | Female Δ HEL/WT vs Male Δ HEL/WT | -1.0623 | 0.9769 | FALSE |

##### 3.20.72 Treg cells - % of CD4 T cells (Panel A)

|  | test | Difference | p\_value | Significance |
| --- | --- | --- | --- | --- |
| c2 | WT/WT vs Δ HEL/WT | 0.9618 | 0.1108 | FALSE |
| c3 | Female vs Male | -0.5411 | 0.5720 | FALSE |
| c4 | Female WT/WT vs Female Δ HEL/WT | -0.2455 | 0.9706 | FALSE |
| c5 | Male WT/WT vs Male Δ HEL/WT | 2.1691 | 0.0046 | TRUE |
| c6 | Female Δ HEL/WT vs Male Δ HEL/WT | 0.6663 | 0.7137 | FALSE |

##### 3.20.73 Treg cells - % of live leukocytes (Panel A)

|  | test | Difference | p\_value | Significance |
| --- | --- | --- | --- | --- |
| c2 | WT/WT vs Δ HEL/WT | 0.0475 | 0.9779 | FALSE |
| c3 | Female vs Male | -0.1033 | 0.8491 | FALSE |
| c4 | Female WT/WT vs Female Δ HEL/WT | -0.1728 | 0.6920 | FALSE |
| c5 | Male WT/WT vs Male Δ HEL/WT | 0.2678 | 0.4356 | FALSE |
| c6 | Female Δ HEL/WT vs Male Δ HEL/WT | 0.1169 | 0.9221 | FALSE |

#### 3.21 Gross Pathology

**Number of animals**

| STRAIN | GENDER | count |
| --- | --- | --- |
| WT/WT | female | 8 |
| WT/WT | male | 8 |
| Δ HEL/WT | female | 7 |
| Δ HEL/WT | male | 6 |

##### 3.21.1 Adrenal gland

##### 3.21.2 Bone

##### 3.21.3 Bone marrow

##### 3.21.4 Brain

##### 3.21.5 Epididymis

##### 3.21.6 Esophagus

##### 3.21.7 Eye with optic nerve

##### 3.21.8 Femur

##### 3.21.9 Gall bladder

##### 3.21.10 Heart

##### 3.21.11 Kidney

##### 3.21.12 Large intestine

##### 3.21.13 Liver

##### 3.21.14 Lung

##### 3.21.15 Lymph node

##### 3.21.16 Mammary gland

##### 3.21.17 Ovary

##### 3.21.18 Pancreas

##### 3.21.19 Prostate

##### 3.21.20 Salivary gland

##### 3.21.21 Seminal vesicle

##### 3.21.22 Skeletal muscle

##### 3.21.23 Skin

##### 3.21.24 Small intestine

##### 3.21.25 Spinal cord

##### 3.21.26 Spleen

##### 3.21.27 Stomach

##### 3.21.28 Testes

##### 3.21.29 Thymus

|  | female WT/WT | female Δ HEL/WT | male WT/WT |
| --- | --- | --- | --- |
| female Δ HEL/WT | 0.300 | NA | NA |
| male WT/WT | 1.000 | 0.6831 | NA |
| male Δ HEL/WT | 0.018 | 0.2051 | 0.0769 |

##### 3.21.30 Thyroid

##### 3.21.31 Tooth

##### 3.21.32 Trachea

##### 3.21.33 Urinary bladder

##### 3.21.34 Uterus

| SPECIMEN\_ID | GENE | GENDER | DATE | Tissue | Ontology |
| --- | --- | --- | --- | --- | --- |
| 28Y-25531 | deltaHEL1 | male | 2017-07-16 | Thymus | MP:0000703:abnormal thymus morphology |
| 28Y-25532 | deltaHEL1 | male | 2017-07-16 | Thymus | MP:0000703:abnormal thymus morphology |
| 28Y-25576 | deltaHEL1 | male | 2017-07-16 | Thymus | MP:0000703:abnormal thymus morphology |
| 28Y-25577 | deltaHEL1 | male | 2017-07-16 | Thymus | MP:0000703:abnormal thymus morphology |
| 28Y-25579 | deltaHEL1 | male | 2017-07-16 | Thymus | MP:0000703:abnormal thymus morphology |
| 28Y-25589 | deltaHEL1 | female | 2017-07-17 | Thymus | MP:0000703:abnormal thymus morphology |
| 28Y-25592 | deltaHEL1 | female | 2018-07-18 | Thymus | MP:0000703:abnormal thymus morphology |
